# Cryo-EM Structures of Prestin and the Molecular Basis of Outer Hair Cell Electromotility

**DOI:** 10.1101/2021.08.06.455374

**Authors:** Navid Bavi, Michael David Clark, Gustavo F. Contreras, Rong Shen, Bharat Reddy, Wieslawa Milewski, Eduardo Perozo

**Author notes:** Correspondence to: **Eduardo Perozo**.

## Abstract

The voltage-dependent motor protein, Prestin (SLC26A5) is responsible for the electromotive behavior of outer hair cells (OHCs). Here, we determined the structure of dolphin Prestin in six distinct states using single particle cryo-electron microscopy. Structural and functional data suggest that Prestin adopts a unique and complex set of states, tunable by the identity of bound anions. Complexes with the inhibitor salicylate show that it competes for the anion-binding site of Prestin. These conformations reveal a novel mechanism of area expansion that depends on the helix flexibility and conformational transitions at the membrane protein interface and putatively affects the physical state of the surrounding membrane. These observations illuminate the structural basis of Prestin electromotility, a key component of the mammalian cochlear amplifier.

## Introduction

Mammals have evolved a highly sophisticated sense of hearing (*1*) characterized by extraordinary sensitivity and the ability to process high-frequency sounds (*2, 3*). Humans can detect frequencies fivefold higher (20 kHz) than reptiles and birds, while the exceptional ultrasound hearing in bats, whales and some rodents (up to ∼160 kHz) allows them to efficiently echolocate (*4–7*). This evolutionary outcome is the result of a mechanism of amplification that relies on the specialized electromotility of the mammalian outer hair cells (OHCs) (*8–10*). The cochlear amplifier depends on the voltage-dependent longitudinal contractions or elongations of OHCs triggered by the concerted action of millions of fast “motor” proteins in the basolateral membrane (up to ∼7000 copies/µm^2^) (*11*). This molecular motor was identified as Prestin (SLC26A5), a piezoelectric member of the SLC26 family of anion transporters (*5, 8, 12, 13*). Prestin is necessary and sufficient to elicit electromotility by transducing changes in transmembrane potential into mechanical forces, and vice versa (*12*). Importantly, Prestin‘s absence or dysfunctions are associated with non-syndromic hearing loss in mammals (*14–16*). In spite of the intense attention to Prestin since its identification as the OHC motor (*12*), fundamental questions remain unanswered, including its oligomeric state (*17–20*), underlying voltage sensing mechanisms and the molecular basis of electromotility (*21*). The lack of high-resolution structures remains a key missing element in defining OHC functional behavior at a molecular level.

OHCs shrink upon depolarization and expand upon hyperpolarization (*10*). These length changes modify the mechanical properties of the basilar membrane and lead to amplification of sound-evoked vibrations. As a first approximation, OHC electromotility does not appear to depend on the action of cytoskeletal elements (*22, 23*). This points to a mechanism where Prestin might exert lateral forces that directly influence the physical state of the bilayer. As proposed, force transmission through an area motor model (*24–26*) requires Prestin to populate at least two distinct conformations characterized by different intra-membrane cross sectional areas, a compact and an expanded conformation. Thus, given enough copies in the basolateral membrane, the sum of Prestin’s molecular scale two-dimensional area changes would then lead to a cellular-level contraction or elongation of OHCs.

Prestin’s conformational landscape is directly modulated by voltage (*12, 27*), anion-binding (*28, 29*) and membrane tension (*26, 30, 31*). Voltage sensing can be estimated from non-linear capacitance (NLC) signals in OHCs (*32, 33*); these represent the electrical signature of conformational changes in Prestin. However, since neutralization of most of Prestin’s intra-membrane changes have been difficult to interpret in the absence of a structural reference (*28*), the molecular underpinnings of this charge movement remain unclear. In fact, Prestin’s NLC is exquisitely sensitive to the nature and concentration of the intracellular anion, leading to the suggestion that Cl^−^ could act as an extrinsic voltage sensor (*29*). The lack of correlation between Prestin’s apparent elementary charge (z) with the ionic valence of the substituting anion suggests that anions might instead be an important part of an electrostatic network that interacts with yet to be defined fixed charges or dipoles (*24*).

In this study, we have used single particle cryo-electron microscopy (cryo-EM) to determine the structure of Prestin under various ionic conditions and in complex with the reversible inhibitor salicylate. These structures, together with mutagenesis, functional data and electrostatic calculations show that Prestin adopts a defined set of states as part of its electromotility cycle. These states point to novel mechanisms of voltage-dependent area changes, highlight the evolutionary differences with SLC26 transporters, and constrain explicit models that help explain Prestin’s behavior as a piezoelectric motor. Finally, they illuminate the interplay between electric fields and mechanical forces that give rise to the cochlear amplifier.

### Cryo-EM Structure of the SLC26A5 homodimer in Cl^−^

To access a biochemically stable Prestin preparation suitable for single particle cryo-EM, a variety of orthologs were explored. We hypothesized that those adapted to detect high-frequency signals (i.e., bats and whales) might exemplify particularly robust systems. An initial screening by fluorescence-detected size-exclusion chromatography (FSEC) led to the identification of a Prestin candidate from the bottlenose dolphin (*Tursiops truncatus*). HEK293 cells expressing dolphin Prestin display the characteristic bell shape NLC with a V1/2 of −61 ± 0.5 mV (mean ± SEM, n=20, Fig. 1A) and similar voltage sensitivity (1/α= 37±3 mV, mean ± SEM, n=20) as those reported for other mammalian systems. Importantly, HEK293 cells overexpressing Prestin at high levels undergo unusually large voltage-driven cell movements under whole cell patch clamp configuration, consistent with robust levels of electromotility (Fig. 1B; Fig. S1A; Movie S1). Expressed in baculovirus-infected mammalian cells, dolphin Prestin is characterized by a main gel filtration peak around ∼150 kDa (predicted molecular mass of ∼75 kDa), suggesting a stable dimer in solution (Fig. S1B,C). We report no evidence of larger oligomeric assemblies.

**Fig. 1.**
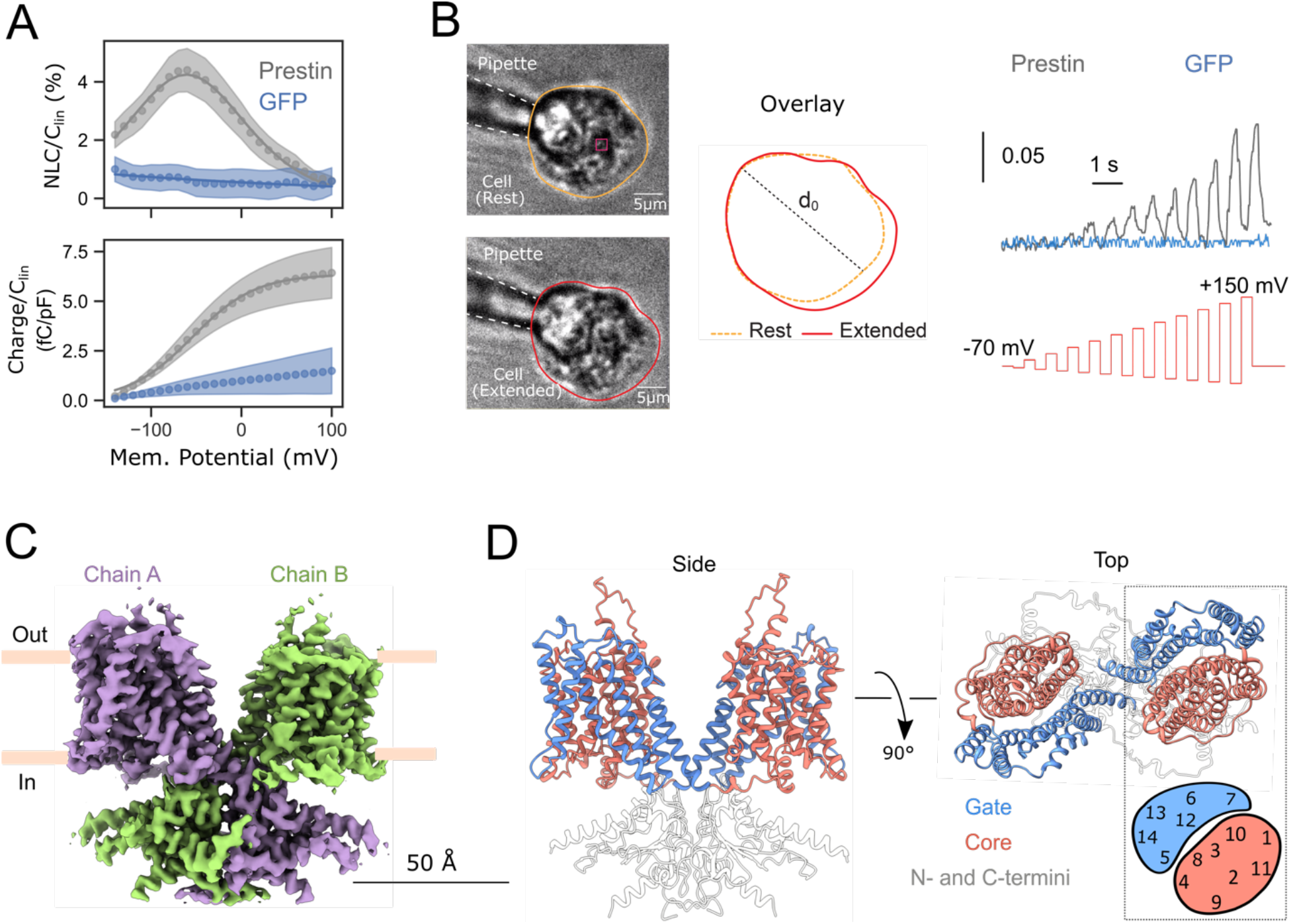
Structure and function of dolphin SLC26A5 homodimer in Cl^−^. (**A**) Normalized change in nonlinear capacitance (NLC=*C-Clin*) and charge movement (right panel) of dolphin Prestin expressed in HEK cells and using patch clamp electrophysiology. Compared to mock (GFP only) transfected cells, dolphin Prestin has a bell shape change in capacitance with V1/2 of −61 ± 0.5 mV (n=20, mean ± SEM). The NLC curve of Prestin (grey) vs transfected cells with GFP alone (blue, n=8) in whole-cell patches. Prestin’s NLC follows a Boltzmann distribution (fit with solid lines). Cell movement is (**B**) Electromotility of Hek293 cells transfected with dolphin Prestin in whole cell configuration (n=7). Cells were lifted off the surface to avoid any possible effect from cell-surface adhesion forces on the cellular movement. Upon changing the membrane potential (the amplitude were increased in each step), the overexpressing dolphin Prestin in HEK293 cells can visibly deform the cells. By tracking the change in the 2D projected area of the cell (red and yellow traces), the cell expands upon hyperpolarization. Cells transfected with GFP only does not sow such response (blue solid line, n=5). Cell movement (right panel) has been normalized with the largest diameter of the cell, d0. (**C**) Cryo-EM density map and the overall structure of the dolphin SLC26A5 homodimer at 3.3 Å nominal resolution when Cl^−^ is the main anion (See methods); the subunits are colored in violet and green. (**D**) Side and top views of Prestin structure colored according to the different domains, with the core domain in red, the gate domain in blue, and the N- and C-terminus and STAS domain in grey. UCSF ChimeraX was used for illustration.

We determined the structure of full-length dolphin Prestin at 3.3 Å using single particle cryo-EM, with Cl^−^ as the main anion (Fig. 1C, D; Fig. S2). It shows Prestin is a symmetric homodimer (C2) with 14 α-helical transmembrane (TM) segments (Fig. S3-S4), divided into two overlapping domains: a core domain (TM segments 5-7and 12-14) and a gate domain (TM segments 1-4 and 8-11) as part of the individual subunit. These domains putatively undergo translational motions relative to each other as expected from an elevator-type transporter mechanism (*34, 35*). The TM regions are domain swapped with the STAS containing -N and -C termini cytoplasmic domains (Fig. 1D). This putative Cl^−^-bound conformation is structurally closest to that of the intermediate state in SLC26A9 (PDB:6RTF) (*35*) (Fig. S5), where a large cavity is observed accessible to the intracellular face of the molecule between gate and core modules. We see no explicit density associated with a bound Cl^−^, as is the case with other SCL26 structures (*34, 35*). When superimposed with SLC26A9 in the intermediate conformation, the core domain displays a rigid body rearrangement characterized by a ∼14° rotation, together with a ∼4 Å translation towards the extracellular side (Fig. S5). This would be equivalent to an elevator movement towards the outward-facing conformation of the transporter. However, in Cl^−^-Prestin, the anion binding pocket is not directly accessible, neither to the intracellular nor to the extracellular face of the molecule.

Interestingly, a comparison of the electrostatic surfaces of SLC26A9 and Prestin points to significant differences in their membrane hydrophobic footprint (Fig. S6). Whereas SLC26A9 displays a broad and almost flat hydrophobic surface (in line with the footprint of a ∼30 Å bilayer thickness) (Fig. S6A), Prestin shows a narrower, more uneven profile tilted some 15° relative to the plane of the bilayer (Fig. S6B). This points to a unique interaction between Prestin’s hydrophobic footprint and the membrane at the lipid-protein interface. The core domain is the site of the canonical SLC family anion-binding pocket, which includes the TM1 helix and the helical macrodipole-oriented TM3 and TM10 pair that buttresses a set of anion coordinating residues (Q97, F101, F137, S398 and R399; Fig. S7). These residues are largely conserved along SLC26 transporter family (Fig. S8). Mutating any of these residues in Prestin leads to major effects on NLC, V1/2 and/or the total charge transfer (*17, 34, 35*).

### SO4^2−^ binding uncovers novel Prestin conformations

Replacing Cl^−^ with SO4^2−^ as the main anion (10 mM Cl^−^ vs. ∼ 130 mM Na2SO4^2−^) reduces dolphin Prestin’s NLC by ∼ 75 % and shifts it ∼80 mV toward positive potentials (Fig. 2A). This is in agreement with previous studies in cetacean Prestin (*36*) and follows the behavior reported in mouse and gerbil (*28, 37–40*). Therefore, at zero mV almost half of its total charge has moved across the electric field (Fig. 2A) and SO4^2−^-Prestin would display maximal conformational heterogeneity. Indeed, the cryoEM structural determination of SO4^2−^-Prestin revealed three distinct new states: two conformations with solvent access to the anion binding pocket (Down I and II) and an Intermediate conformation preceding the Up state (Fig. 2B-D, Fig. S9-S12). When compared with Cl^−^ -Prestin (Up), the major differences among these states were quantitatively described based on two measurements between Core and Gate domains (Fig2. B): *z*, the position of residue R399 (TM10, Core domain) relative to V499 in TM14 in the gate domain; and *d*, the gap that separates the TM3-TM10 helical dipole.

**Fig. 2:**
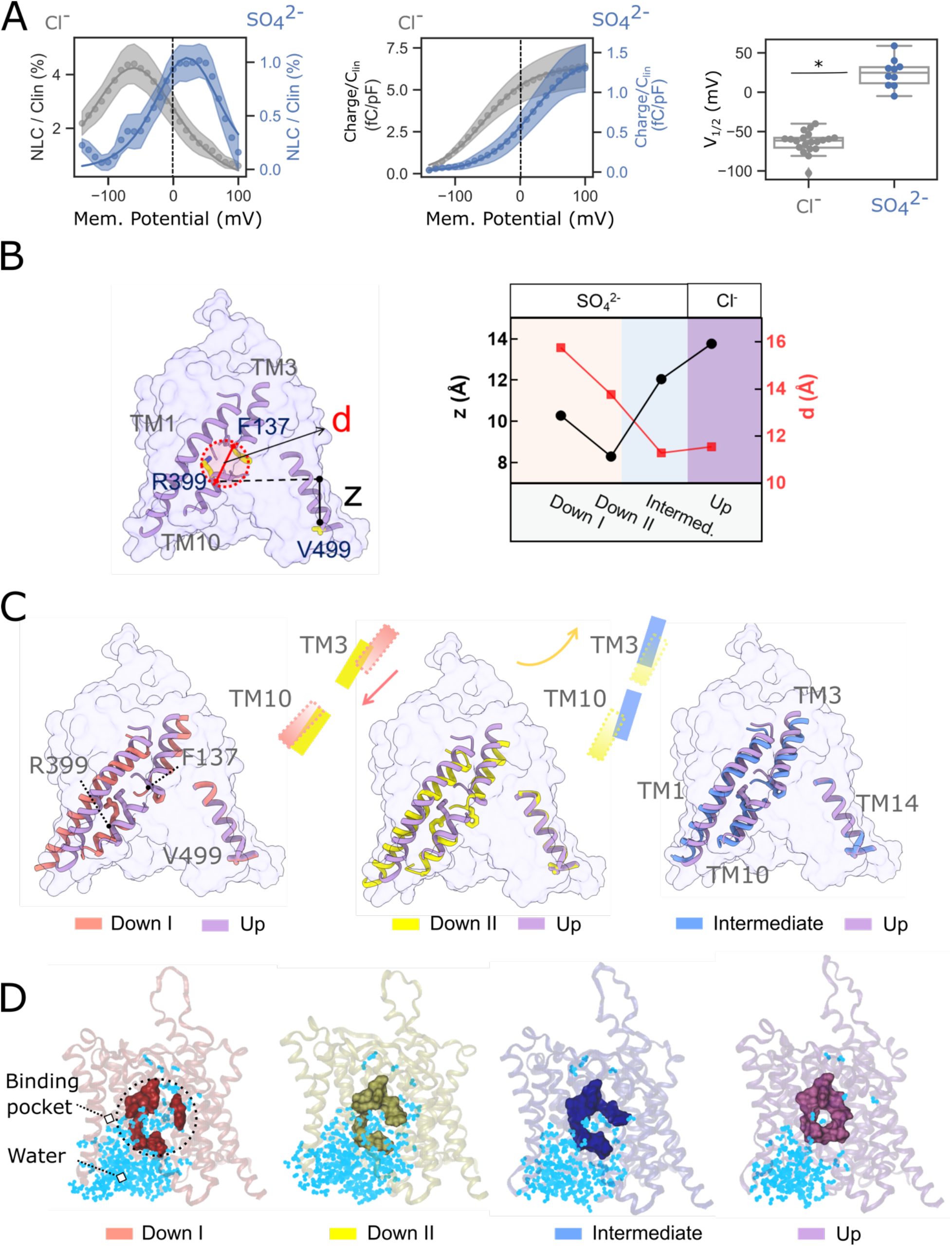
Sulfate (SO4^2−^) drives Prestin towards down and intermediate states at zero membrane potential. (**A**) Normalized NLC and charge transfer measurement of transiently transfected HEK293 cells with dolphin Prestin with Cl^−^ and SO4^2−^. Compared to Cl^−^, when SO4^2−^ is the main internal and external anion, the NLC amplitude is attenuated and the V1/2 is shifted to ∼ +20 ± 3 mV (mean ± SEM, n=10; Student’s t-test, *, P=0.002). (**B**) Wild type Prestin structure in a SO4^2−^-based medium were obtained at three distinct configurations using single particle Cryo-EM technique. We considered two reaction coordinates (*z* and *d*) for determining the structural landscape of Prestin. *z* is the vertical distance between R399 backbone carbon and V499 backbone carbon (a residue on TM14 that resides on the protein-water interface; *d* is the distance between R399 backbone carbon and F137 backbone carbon which is equivalent to the binding pocket diameter, assuming the pocket as an sphere. All three residues are shown in Sticks. *z* and *d* has been measured across different states; Accordingly, from left to right, we have named the Prestin states: Down I, Down II, Intermediate (all SO4^2−^) and Up (Cl^−^). The color code we have chosen for these states are red, yellow, blue and violet respectively. (**C)** Movement of the anion binding site from Down to Up states. From Down to Up, first the electrical field becomes more focused due as the TM3 moves towards TM10 to form a helical dipole (i.e., the pocket becomes tighter, shown by the red arrow); this is followed by an elevator-like movement of the binding site from Down to Up, as shown by the cartoons and the yellow arrow. Down I, Down II and Intermediate are in the presence of SO4^2−^, while in the Up state, Cl^−^ is the main anion. The structures were aligned based on residues 460 to 550 (TM13-TM14, the least mobile part of Prestin). For clarity, only TM14 has been illustrated. UCSF ChimeraX was used for illustration. (**D**) The 1 µs all-atom molecular dynamics simulation of these states in POPC lipid bilayer reveals the dynamics of the anion-binding pocket as well as the water hydration (cyan) at different states. The number of waters in the intracellular cavity of Prestin reduces from Down to Up states (left to right). In addition, the pocket is the most apart in the Down I state (weak anion binding) and most confined in the Up (Cl^−^) state. For clarity, only the water molecules within 8 of the residues Q97, F101, F137 V397, S398, R399, E280 and E404 have been screened and illustrated using VMD.

When linked as part of a trajectory of conformational rearrangements, these states point to the overall working cycle of Prestin, from Down to Up conformations (Figs. 2 and 3). In the Down states the core domain is positioned towards the intracellular face of the membrane (z between 8-10 Å). While in the Up state, the core domain moves ∼ 6 Å towards the extracellular face of the membrane (Fig. 2B,C). This movement (Fig. 2C, Movies S2, and S3) is reminiscent of a partial “elevator” rearrangement seen in other SLC transporters where the uppermost conformation in Prestin would goes beyond SLC26 transporters in an occluded (intermediate) state (*35*). The core domain movements associated with the substitution of Cl^−^ by SO4^2−^ also trigger a widening of the anion binding pocket (parameter *d*). In the transition from Down I to Down II to the Intermediate state, TM3 moves towards TM10 (∼4 Å), forming a stronger helical dipole and a tighter anion-binding pocket (Fig 2C, Movie S2). Long scale (1 µs) all-atom MD simulations shows that these structural changes lead to substantial changes in water penetration, where the anion binding pocket is fully water accessible in states Down I and II but water accessibility is sharply reduced in the Intermediate state (Fig. 2D). In the Down I state, the ∼16 Å gap between TM3 and TM10 helices almost fully hydrates the anion-binding site, weakening the helical dipole between TM3 and TM10.

**Fig. 3:**
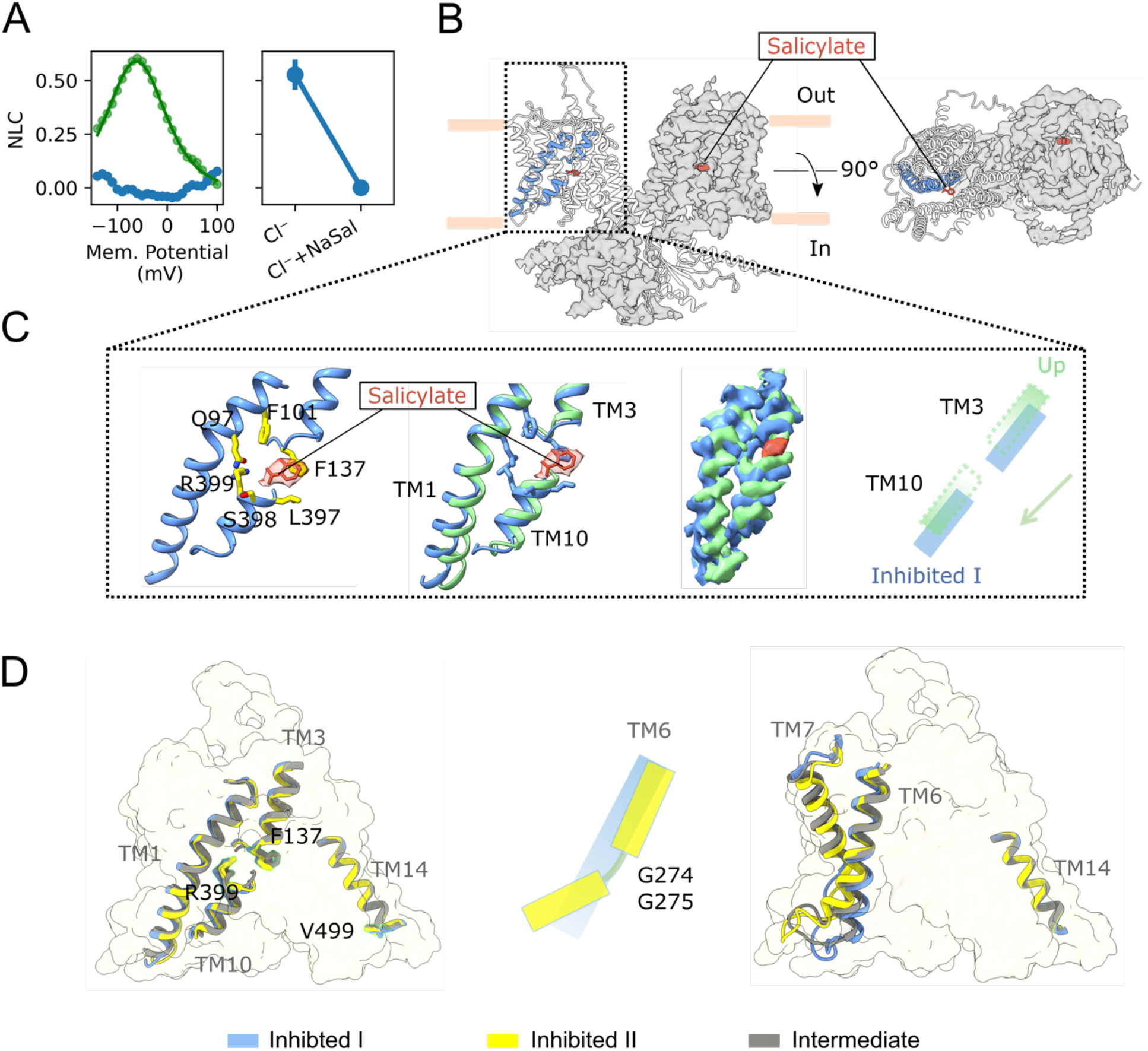
Structural basis of Prestin inhibition by salicylate. (**A**) 10 mM salicylate inhibits Prestin function and flattens NLC and charge transfer when expressed in HEK293 cells (whole cell patch clamp electrophysiology, n=7; Student’s t-test, * P=0.005). (**B**) Resolved structural densities indicates salicylate (shown in red) bound to anion pocket of Prestin, located at the core-gate interface. (**C**) A zoomed inset of the anion-binding site formed by residues Q97, F101, F137, L397, S398 and R399 (shown in yellow), situated in the space confided by TM1 helix and the micro-dipole of TM3 and TM10. An overlay of the EM densities for sensor Up (Cl^−^) versus Inhibited I (Cl^−^+salicylate, left panel). For salicylate to be able bind to the pocket, TM10 helix should move downward to create enough space for accommodating the larger size salicylate compared to Cl^−^. Salicylate molecule and density has been shown in salmon color. (**D**) The Inhibited I (Cl^−^+salicylate) and Inhibited II (Cl^−^+salicylate) structures and the intermediate state are overlapping at the anion-binding site, (TM13-TM14 helices were aligned). The major difference between them is at the TM6-TM7 helical region of the core domain. The TM6 helix is kinked in the inhibited II state (yellow). UCSF ChimeraX was used for the illustration.

### Salicylate traps Prestin in an Intermediate state by occluding the anion-binding pocket

The amphiphilic drug Salicylate is known to cause tinnitus (*52, 53*), abolishing the NLC and electromotility in mammalian OHCs (*12, 29, 54*), while also inhibiting Cl^−^ transport in its non-mammalian homologs (*55*). While it has long been the best-characterized Prestin inhibitor, the molecular basis for Salicylate inhibitory effects remains unclear. Whole-cell patch clamp recordings show that 10 mM Na-salicylate flattens dolphin Prestin’s NLC curve across the physiological voltage range (Fig. 3A, Fig. S13A) in ways reminiscent to that observed in a variety of mammalian systems (*12, 29, 54*). We determined the structures of the Prestin-salicylate complex in the presence of either, Cl^−^ (360 mM) or SO4^2−^ (120 mM) at 3.8 Å and 3.7 Å, respectively (Fig. 3B, Fig. S13B-S15). In each case, 50 mM sodium salicylate was present. Regardless of the available anion, the anion-binding pocket appears to be predominantly occluded by a robust density corresponding to salicylate (Fig. 3B and S13).

Salicylate likely outcompetes any other bound anion at the binding pocket, and given that no anion can be resolved even under saturating Cl^−^ or SO4^2−^ concentrations (Fig 4B), this density helps confirm the overall nature of ion coordination at the anion-binding pocket. In both structures, the salicylate density snuggly fits in the pocket formed by residues Q97, F101, F137, L397, S398 and R399 (Fig. 3C and Fig. S13B). Comparing the anion binding site in the Up conformation with that of the Cl^−^+salicylate structure (Inhibited I), shows how the core domain TM10 moves ∼ 3 Å downward, creating enough space to accommodate the larger salicylate (Fig. 3C). The salicylate-inhibited structures (Cl^−^ and SO4^2−^) appear to converge into an Intermediate state (Fig. S16A) that display common conformations at the anion-binding site (Fig. 3D, Fig. S16B). Important differences are also observed at the TM6-TM7 helices (Fig. 3D, Fig. S16C), in the periphery of the gate domain, facing the lipid bilayer. These represent unique conformations with fundamental roles in defining electromotility (see below). Based on these results, we suggest that salicylate inhibits Prestin’s motor function by immobilizing the relative displacement of the core domain around the anion-binding site.

**Fig. 4:**
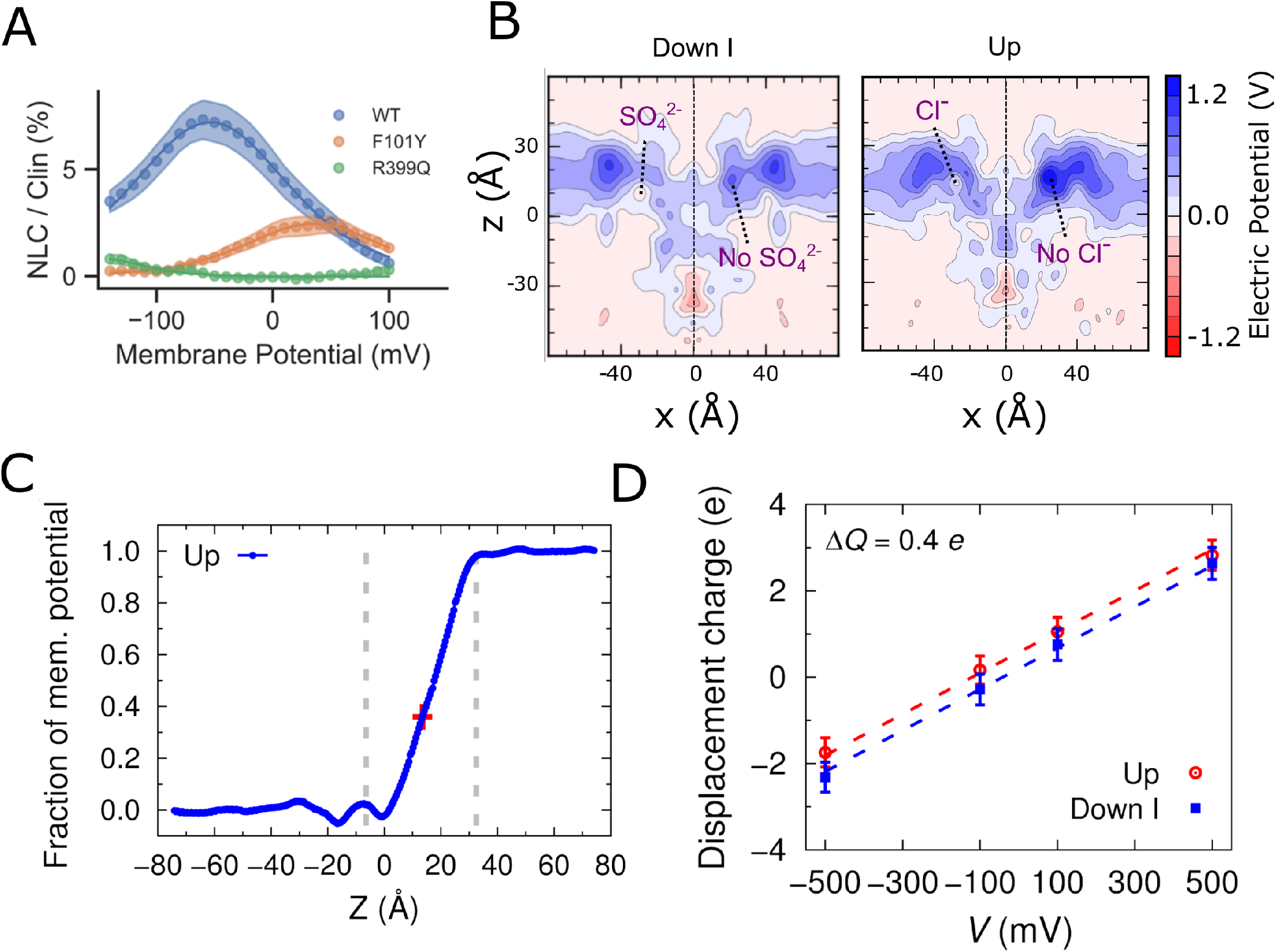
Electrostatic calculations and charge transfer of Prestin across the membrane. (**A**) Mutation of the key residues in the anion binding pocket either completely abolishes the NLC (R399Q) or right shifts the V1/2 by more than 60 mV (F101Y); a similar effect has been observed in other Prestin homologs (51). (**B**) Snapshots from the MD trajectories of the systems, and calculation of the electrostatic potential across the membrane at two states, the Down I state (with SO4^2−^ in the left cavity, and without SO4^2−^ in the right cavity) versus Up (with Cl^−^ in the left cavity and without any Cl^−^ in the right cavity). The *x-z* plane is crossing the two central anion-binding sites. In both models, the positive field is mainly focused around the transmembrane mid-plane and around the anion-binding site, creating an attractive (blue) field for the binding of the anion. However, in the Up state the field is more positive around the mid-plane compared to the corresponding region in the Intermediate state. In both cases, the presence of the anion only partially neutralizes (∼35 %) the positive field around the bilayer mid-plane. Note that the actual size of the simulation box is larger than what has been illustrated here (see methods). (**C**) Averaged 1-D fraction of membrane potential in the z direction along the two central binding sites (shown as dashed blue lines in panel A with the central binding sites highlighted using the red cross symbols). The 1-D and 2-D maps were directly extracted from the ensemble averaged 3-D fraction of membrane potential map. The location of the phosphate atoms of the outer and inner lipid leaflets along the z axis was highlighted with dashed gray lines.). (**D**) Displacement of charge for Prestin in the Up and Down I conformations at different transmembrane potentials. Data are mean ± SD. The gating charge between the two states is 0.38 e calculated as the offset constant between the linear fits.

### Prestin’s voltage dependence is defined by the interplay between fixed charges and bound anions

A large body of experimental evidence points to the complex nature of the voltage sensor in Prestin (*28, 29, 38, 41*), were intrinsic charges or dipoles would interact with bound anions to generate the NLC, ultimately driving electromotility. Indeed, neutralizing residue R399 fully eliminates the NLC (Fig. 4A) in mammalian Prestin (*17*) while neutralization of a series of fixed charges (K276, K359, K56 and K449) have a partial effect on unitary charge transfer (Bai et al 2009). And yet, Prestin voltage dependence shows an acute dependence to the binding and occupancy of anions (Fig. 2). The present set of Prestin structure provides us with a unique opportunity to interrogate different conformations based on the bound states of the anion binding pocket. The major effect of anions on Prestin’s function is to shift Prestin’s V1/2 along the voltage axis (Fig. 2). This simple experimental manipulation allows for a structure-based estimate of Prestin’s electrostatic properties Fig. 4B, and by extension, a glimpse at the mechanism of voltage dependence. Two important results come to fore. In spite of the presence of a relatively large intracellular water-filled cavity, Fig. 4C shows that the electric field displays fairly limited focussing (in contrast to voltage sensing domains in ion channels and enzymes (*42, 43*)). Within this profile, it is clear that the anion-binding pocket is sensing only ∼30-35% of the field (Fig. S17). As a result, positive fields at the anion-binding pocket are only partially neutralized in the presence of a bound anion, regardless of their valence Figure 4A. In principle, this would explain the apparently paradox posed by the non-correlated effects of anion valence with the elementary charges estimated from NLCs (*28, 38*). This observation would also suggests that while anion binding is a significant component of charge movement other intrinsic contributions are needed to fully account for Prestin’s voltage dependence.

Voltage dependence can be estimated from the movement of the charged residues within a defined electric field (*33*). Gating charge calculations carried out based on the two extreme conformations (Down I to Up states) estimate that ∼0.4 unitary charges move across the electric field in the transition from Prestin’s Inhibited II state to the Up state (Fig. 4D). These estimates are roughly consistent with earlier evaluations of charge movement in Prestin (*30, 44*) and show that the presence of residues R399, K276, K359, K56 and K449 at the mid-plane of the field have the largest contributions to the net positive electric field in this area. Mutation of any of these residues either completely removed the NLC (R399) or diminishes it (K276, K359, K56 and K449) (*45, 46*). It is important to note that while K449 is conserved amongst most SLC proteins, the equivalent charge at R399 is unique to Prestin (Fig. S8). Accordingly, mutating R399 to Q, S or E in our molecular dynamics simulations partially neutralizes the positive field, while only the potential around the helical-dipole remains positive (Fig. S18). Thus, in addition to the TM3-TM10 helical dipole, the positive charges located at the bilayer mid-plane (particularly R399), create an attracting field for anion binding (Down I to Down II) and likely define Prestin’s voltage sensor.

### TM6 helix flexibility is an evolutionary conserved feature of Prestin electromotility

The multiple Prestin structures described above contain two remarkable pieces of evidence that provide a novel framework to understand Prestin-based electromotility. First, Prestin appears to have a strong influence in the shape of its surrounding micelle (Fig. 5A,B). In the Intermediate and Up conformations, the micelle surrounding Prestin is shaped as an elongated oblate which thickens towards the longitudinal ends of the dimer (viewed extracellularly). In contrast, the micelle in the Inhibited II state is not only wider and thinner along the sides, but it shows a “notch” that induces a dramatic thinning of the micelle. Using long-scale (1 µs) molecular dynamics simulations, we show that TM6 bending dramatically thinned the lipid bilayer by up to ∼25% at the lipid-Prestin interface (Fig. 5B, S19).

**Fig. 5:**
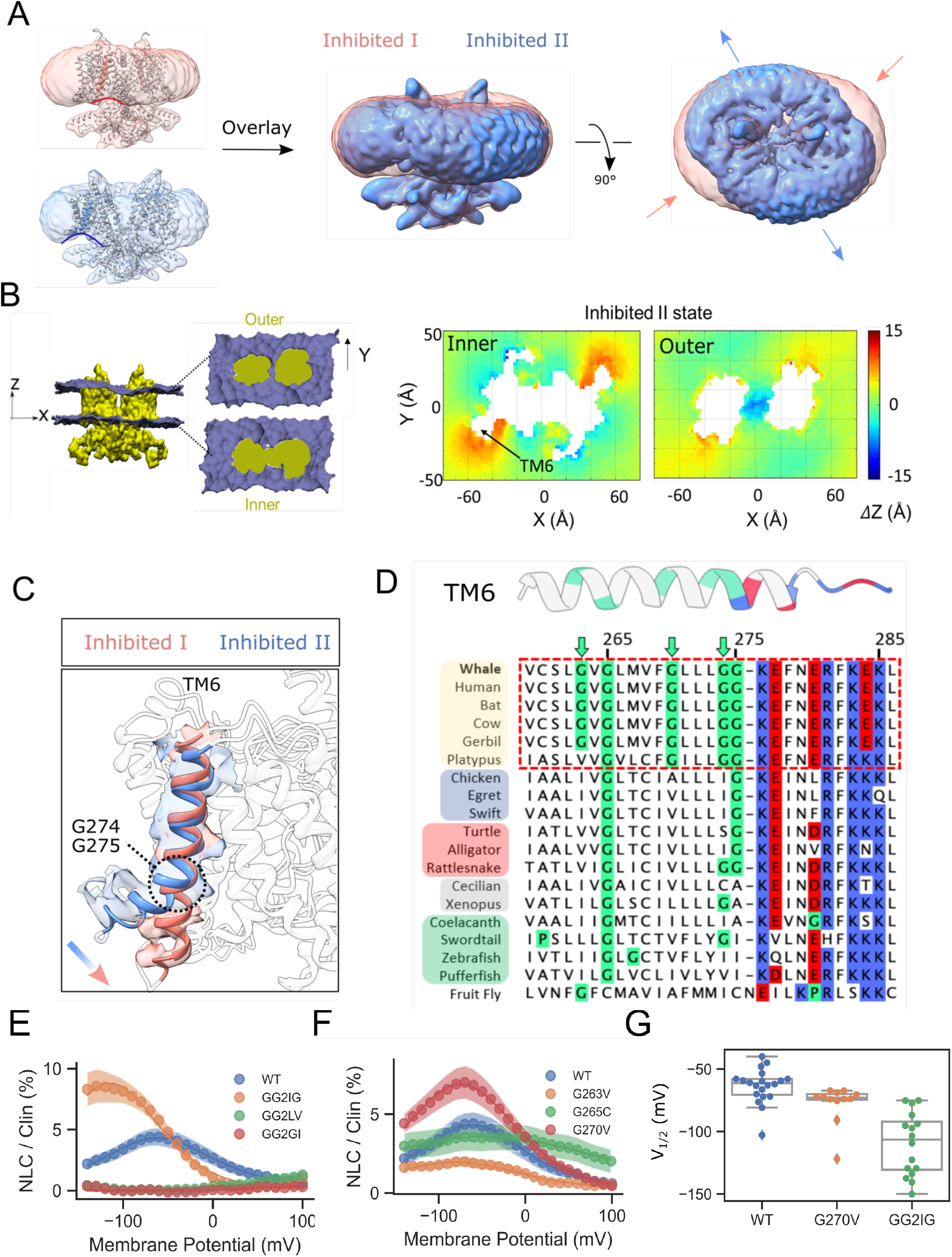
TM6 helix flexibility and the evolutionary transition from transporter to motor protein in Prestin. **(A)** Comparing the change in the micelle morphology between two salicylate-inhibited structures Inhibited I (Cl^−^) and Inhibited II (SO4^2−^+Salicylate) states. The overlay of the two states shows drastic changes in the micelle thickness especially around TM6 region in addition to the overall changes in the micelle in-lane direction, both indicative of major structural rearrangements between the two states. (**B**) MD simulation of Prestin (Inhibited II state) equilibrated in POPC lipid bilayers. The cross-sectional area of outer and inner monolayer with mapped leaflet coordinate in the Z direction (across the membrane thickness) using all-atom molecular dynamics simulations (1µs). The comparison was made between Inhibited II (SO4^2−^) and Up (Cl^−^) states. Points close to TM6, where calculations of local thickness is possible, the membrane is >25 % thinner compared to other regions, indicating severe bilayer distortion in this area. (**C**) A zoomed inset of the structural overlay of the gate region in Up and Inhibited II states. TM6 region has been colored while the rest of the helices are transparent for clarity. TM6 bends around glycine residues at position 274 and 275. To better show the TM6 elbow density the EM density maps used in in this alignment were not postprocessed nor sharpened. (**D**) Evolutionary sequence alignment of TM6 in Prestin across different species. The focus is on the glycine residues that may underlie the flexibility of this helix during its motor function. (**E**) Mutation of evolutionary conserved glycine residues such as G274 and G275 has major consequences on NLC and charge transfer. Double mutation of these two consecutive glycine residues diminishes electromotility. In addition, GG to GI mutation completely removed the NLC, while GG to IG mutation only left-shifted the NLC to −114 ± 3 (n=16; One way Anova, *, P=0.005). (**F, G**) Mutation of other conserved Glycine residues along the TM6 helix shows their effects on the Prestin NLC (B) and V1/2 (n>12; Student’s t-test, *, P=0.01). (**G**) Mutation G270V the residue only conserved in mammals left shifts the NLC to −75 ± 0.5 (n=12), while other G263V and G265A almost completely abolishes the NLC.

Second, these micelle distortions are a consequence of distinct changes in the conformation of TM6 and TM1, and can be stabilized depending on the nature of the bound anion/salicylate. Compared to a relatively straight helical configuration in the Up state, TM6 helix bends ∼65° outwardly most noticeably in Inhibited II (Fig. 5C). We suggest that this helical hinge-bending, the “electromotility elbow”, underlies the putative transition between “compact” and “expanded” Prestin states. In fact, estimation of the cross-sectional area of the dimer and monomer (Fig. S20, Movie S4), between the Down and Up conformations point to a ∼10% increase in the membrane area footprint (at the upper end of lateral tension effects in bilayers). This is fully consistent with the process of OHC electromotility, where cell elongation coincides with an expanded Prestin state (hyperpolarization), while depolarization-driven contractions correlate with a compact Prestin state (*25, 30, 50-54*). This rearrangement also reduces the hydrophobic length of the protein facing the lipid bilayer, essentially narrowing the membrane thickness footprint (Fig. 5A-C, S19).

At the core of this structural rearrangement micelle distortions, we find a series of glycine residues that seem to endow TM6 with its unusual flexibility. While G265 appears to be highly conserved across all vertebrates (*13*), a series of 3 additional Glycines appear only among mammals (G263, G274 and G275) (Fig.5D) and helps define the phylogenetic relations between *bona fide* mammalian electromotive OHCs with and those of other vertebrates (Fig. S21). Two of these residues, G274 and G275, are located precisely at the hinge that allows TM6 bending in the inhibited II state. In contrast, the transition from inward-facing to the intermediate state in SLC26A9 seems to only display rigid body motions (*35*). This emphasizes the idea that TM6 flexibility may be an intrinsic requirement for Prestin-based electromotility in mammals. Notably, the C terminal end of the TM6 is also capped by a cluster of charged residues (Fig. 5C) a fact that likely contributes to the membrane thinning effects of this hinge bending movement.

To assess the importance of the glycine residues in TM6, we systematically mutated them and the individual mutants were subsequently evaluated by patch clamp electrophysiology. Mutating G274 and G275 to Leucine and Valine (GG-LV) abolishes Prestin NLC (Fig. 5E-G). Note that, compared to the wild type Prestin, none of the mutations showed any detectable difference in their level expression in the plasma membrane (Fig. S22). This strongly suggests that these glycines provide TM6 with the necessary flexibility for the outward bending that observed in the Inhibited II structure. Additional evidence derived from normal mode analysis helps us compare the mobility differences between TM6 helices in Prestin and SLC26A9 (Fig. S23). TM6 in SLC26A9 displays the same level of mobility compared to the rest of the helices in the gate domain. Prestin TM6 on the other hand shows considerably higher mobility compared to the rest of the structure, particularly at positions G274-275.

The structural rearrangements between these states are not limited to the anion-binding site. Fig. S24, summarizes the conformational changes all SO4^2−^-stabilized structures when aligned and compared to the Cl^−^ (Up) structure. Following the movement of the binding pocket from Down to Up states, the inter-helical space between the gate and core domains decreases significantly (Movie S4). We find that the largest conformational changes along the periphery helices: TM5b (gate), TM6-TM7 (gate) and TM8 (core) (Fig. 3C). We suggest that this helical hinge-bending, the “electromotility elbow”, together with localized rearrangements on TM5, TM7 and TM8 underlies the putative transition between “compact” and “expanded” Prestin states.

### A putative molecular mechanism for OHC electromotility

The process by which SLC26A5 evolved from an electrogenic anion cotransporter into a voltage dependent piezoelectric motor continues to be one of the outstanding questions in the field. What underlying mechanisms allow Prestin to efficiently transduce membrane voltage and generate cellular-level mechanical forces, when other SLCs cannot? In SLC26A9 (*34, 35*), the electrogenic conformational changes that support transport (the elevator mechanism) do so with minimal changes in cross-sectional area (< 5%). In this comparison, the transition of SLC26A9 from Inward-facing to Intermediate would be equivalent to a Prestin transition from Down to Up states (and yet inaccessible to the extracellular solution). In Prestin, movements in the binding pocket appear to be transmitted allosterically to the helices in the gate domain (TM6-TM7 helices at the protein periphery). During the transitions from Down to Up conformation, the intra-helical spaces of the core domain contracts, leading to the reorientation of the TM6 electromotility elbow and TM1 helix. This generates a ∼10% reduction in its cross-sectional area (Fig. 6A, Fig. S20). This rearrangement also reduces the hydrophobic length of the protein facing the lipid bilayer, essentially narrowing the membrane thickness footprint by ∼25 % (Fig. 5B,S19). This level of membrane deformation is energetically consequential for somatic movement particularly given the high density of Prestin particles at the OHC lateral wall (*11*). Aligned with these observations, electromotility and NLC are strongly influenced by the thickness of the bilayer (*56, 57*) as well as by membrane tension generated from surface area changes (*27, 30, 31*).

**Fig. 6.**
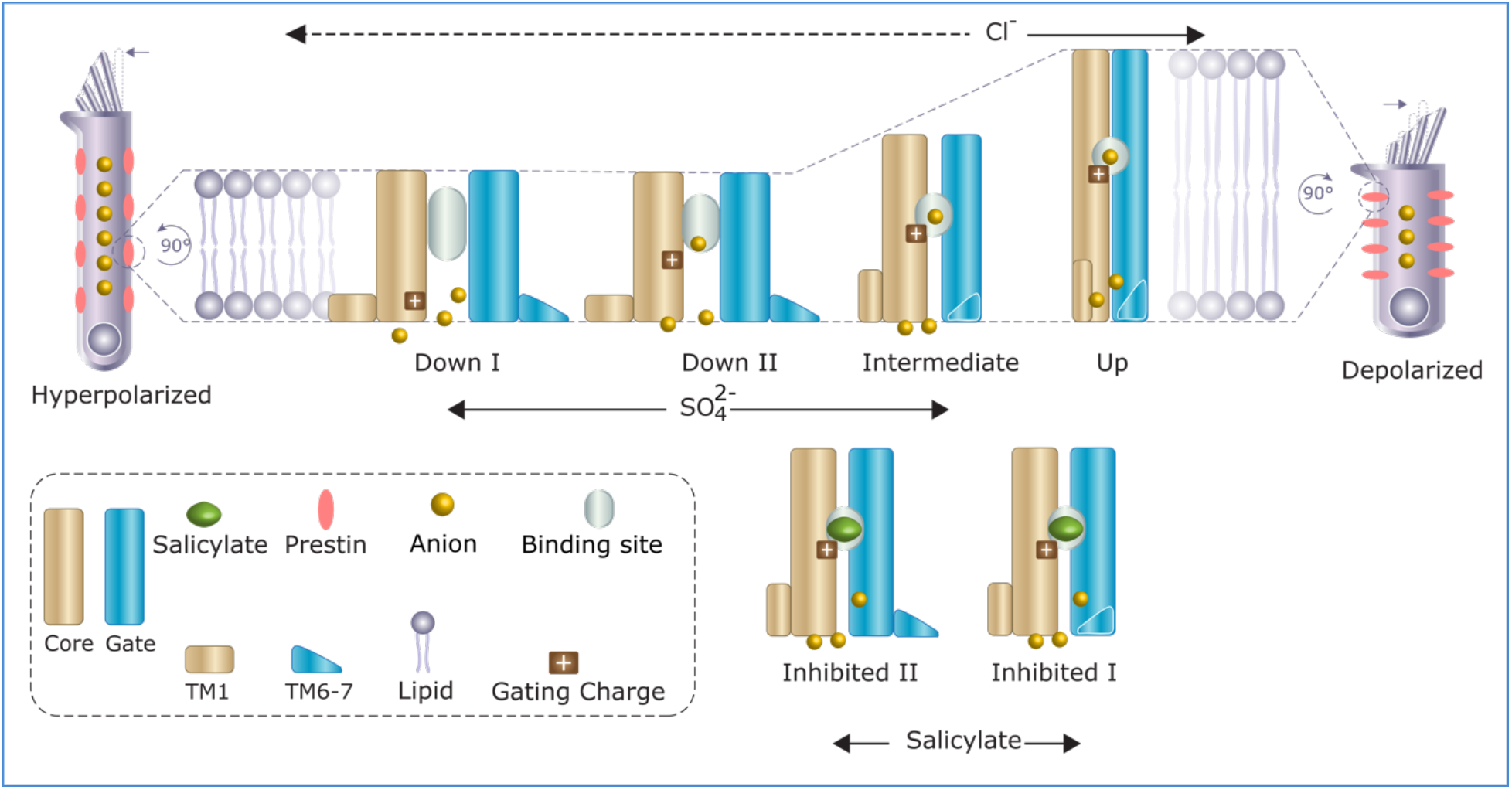
Structural basis of Prestin’s voltage sensitivity and somatic electromotility. Structural transition of the mammalian the Prestin from Down to Up conformation upon membrane depolarization (left to right, top panel). The cartoons shows how Prestin senses the voltage and how it conforms correspondingly. The anions are attracted to the focused positive potential field of the anion-binding pocket, which acts as “voltage sensor”. Upon depolarization, the voltage sensor moves from down to up, which is accompanied by an eccentric contraction of the Prestin’s inter-membrane cross-sectional area (10%) as well as a major increase in the hydrophobic thickness of the protein (exaggerated cartoon). If the plasma membrane follows the footprint of Prestin during the process of Prestin charge transfer, that means that the plasma membrane would thicken and the cell contracts upon membrane depolarization. This is consistent with the OHC movement (contraction) upon applying depolarizing potentials (*22*). Moreover, salicylate physically occludes the pocket and inhibits the voltage sensor movement; therefore, it inhibits Prestin’s charge transfer and electromotility.

Prestin is present in the lateral wall of OHCs at very high densities (∼5,600-7000/µm^2^) (*11*) where they may interact cooperatively, directly, or through yet to be identified adaptor proteins. The efficiency of force transfer to the OHC lateral membrane obviously depends on how individual motors are arranged in the plasma membrane, whether Prestin is polarized (vectorially arranged) or randomly oriented. A quick estimate of the potential mechanism can be based on a model of the OHC lateral membrane with Prestins fully polarized along the longitudinal direction of the area change. Under these conditions, the maximum expected change in length of hair cell would be ∼ 2.6 µm, or 5 % elongation relative to the initial length (See methods; Eq. (3)). This calculation assumes the highest reported copy number of Prestin dimers (7000/ µm^2^), a typical OHC area (∼1500 µm^2^) and the maximum change in the Prestin cross-sectional area (e.g., ∼750 Å^2^ for the Down to Up transition). This estimate is fully consistent with existing experimental determinations where it is estimated to be within ∼4% of length change (*47, 51, 59*), and suggests that electromotility in OHCs is likely intrinsic to the changes in the physical state of the plasma membrane.

Here, we propose that somatic electromotility is derived from novel conformational rearrangements involving Prestin’s transmembrane helices generate vectorial bilayer deformations. Indeed, the conformational trajectories derived from the present set of structures indicate that there are at least four stable energy barriers along Prestin’s electromotility cycle (Fig. 6B). Our data supports the idea that at the molecular level electromotility is more complex than a simple elevator transition in transporters (*24, 60*),. These movements are generated through coupling between the core and gate domains of the individual Prestin monomers, leading to fast and reversible changes in the cross sectional area of thousands of Prestin dimers. The electromotility cycle includes states where the voltage sensitivity is both, intrinsic to Prestin (e.g., Down I state) and where anions are extrinsically involved (e.g., Up state).

A definitive understanding of the molecular and cellular basis of OHC electromotility will necessitate consideration of additional structural rearrangements and potentially other molecular actors. At the dimer level movement and twisting of the Prestin’s cytoplasmic domain is evident when compared to those in the transmembrane domain (Video S4). These movements might have consequences on force transmission through adaptor proteins and other additional factors yet to be identified. Translating any molecular-level forces and movements to the cellular context will have to contend not only with the unusually high density of Prestin in the membrane, but also with its relation to the complex network of sub cisternae membranes and cytoskeletal elements comprising OHC’s basolateral membrane (*25*). Ultimately, new molecular information on Prestin and its potential interaction partners on the cytoplasmic side will need to be combined with macroscopic cell mechanics information to generate a multiscale model of electromotility.

## Materials and Methods

### Cell lines

GnTI^−^ cells used for protein expression and purification were obtained from ATCCC (ATCC CRL-3022). Suspension HEK293S GnTI^−^ cells expressing dolphin Prestin were grown at 37 °C and 7.8% CO_2_ in t FreeStyleTM 293 expression medium (Gibco Fisher Scientific) supplemented with 2% heat-inactivated fetal bovine serum (FBS) and 10 µg/ml penicillin/streptomycin. Adherent HEK293T cells were obtained from ATCTC (ATCC CRL-1573) and were grown in Dulbecco’s Modified Eagle’s Media (DMEM, Gibco Fisher Scientific) supplemented with 10% FBS, at 37 °C and 5% CO_2_. Sf9 cells (Thermo Fisher, 12659017) were cultured in SF-900 II SFM supplemented with 10% FBS and 10 µg/ml Gentamicin at 28 °C.

### Reconstruction of Prestin construct

The primary dolphin (*Tursiops truncates*) Prestin construct was a generous gift from Shi lab (*1*). Dolphin Prestin was subcloned into a modified pEG BacMam vector containing a C-terminal HRV 3C protease, eGFP, and His-8x using restriction sites 5′ NotI and 3′ XbaI. Additionally, the full-length coding sequence of dolphin Prestin was subcloned into the same vector without the protease site, eGFP, and His tags for electrophysiology recordings. Our eGFP alone coding sequence was subcloned into a pcDNA3 vector. P0 Baculovirus was generated via the Bac-to-Bac method (Invitrogen) using Cellfectin II (Thermo Fisher, 10362100). QuikChange site-directed mutagenesis method (Agilent) was used for introducing mutations GG-GI, GG-IG, GG-LV, G265C, G263V and G270V, R399Q and F101Y into dolphin Prestin using KOD DNA polymerase (71085 - EMD Millipore). All mutant and wild type constructs were confirmed by DNA sequencing prior to structural and electrophysiological experiments.

### Purification of dolphin Prestin

P0 virus was amplified once to yield P1 baculovirus, which was used to infect HEK293S GnTI^−^ cells at a 1:10 v/v ratio. After 20-24 h incubation at 37 °C, 10 mM sodium butyrate was added to the cells and the culture was transferred to 30 °C. Cells were collected 50–56 h post infection, washed in phosphate-buffered saline (PBS) pH 7.4, collected by low speed centrifugation, flash frozen and stored at −80 C for later purifications. For purification, all steps were performed at 4 °C. The flash frozen cell pellets were thawed in a room temperature water and were re-suspended and dounce homogenized in Cl^−^ or SO_4_^2−^ based buffer, which for short we refer to as buffer A and buffer B. Buffer A contained 360 mM NaCl, 20 mM Tris-Hcl, 3mM DTT, 1mM EDTA at pH 7.5 and buffer B contained 125 mM Na_2_SO_4_, 5mM Mg(OH)_2_, 20 Tris-OH, the pH was adjusted to 7.5 with 10-15 mM methanesulfonic acid. Protease inhibitor cocktail of 1 μg/ml Leupeptin, 1 μg/ml Aprotinin, 1 μg/ml Pepstatin, 100 μg/ml Soy inhibitor, 1 mM Benzamidine, 0.2 mM PMSF, 0.1 mg/ml AEBSF and 10 μg/ml DNase as well as a cOmplete™ Protease Inhibitor Cocktail tablet (Roche) were added to the solutions. Protein was extracted with a final concentration of 1% *n*-dodecyl-β-D-maltopyranoside (DDM; Anatrace), 0.2% cholesteryl hemisuccinate (CHS, Anatrace), for 90 min. Solubilized supernatant was isolated by ultracentrifugation and the supernatant was incubated for 2 h with 2 mL CNBR-activated Sepharose beads (GE healthcare) coupled with 4 mg high-affinity GFP nanobodies (*2*). Beads were collected by low-speed centrifugation and washed in batch with main-buffer containing 0.05% DDM (Anatrace), 0.01% CHS. Beads were transferred to plastic column and further washed (each wash step was four column volume), exchanging step-wise to buffer containing buffer A (or buffer B) with 0.02% GDN. Additional protease inhibitor were added in each wash step (1 μg/ml Leupeptin, 1 μg/ml Aprotinin, 1 μg/ml Pepstatin, 100 μg/ml Soy inhibitor). Protein was cleaved by HRV 3C protease (*3*) for 2-4 h, concentrated and subjected to size-exclusion chromatography (SEC) on a Superose 6, 10/300 GE column (GE Healthcare) with running buffer including buffer A (or buffer B), 0.02% GDN, 1 μg/ml Aprotinin and 1 μg/ml Pepstatin. Peak fractions were collected and concentrated using 100 kDa molecular weight cut-off centrifugal filter (Millipore concentrator unit) to 2–3 mg/ml. The concentrated protein was immediately used for cryo-EM grid freezing step. For samples with Salicylate, 50 mM Na-Salicylate was added to the purified protein prior to freezing the cryo-EM grids.

### Cryo-EM sample preparation and imaging

Quantifoil 200-mesh 1.2/1.3 grids (Quantifoil) were plasma cleaned for 30 s in an air mixture in a Solarus Plasma Cleaner (Gatan). Purified Prestin sample was applied onto the grids and were frozen in liquid nitrogen-cooled liquid ethane in a Vitrobot Mark IV (FEI) using the following parameters: 3.5 μl sample volume, 2.5 s, 3.5 s, 5 s blot times (blot time varied from sample to sample), blot force 3, 100% humidity, at a temperature of 22 °C and double filter papers on each side of the vitrobot. Grids were screened on a 200 kV Talos side entry microscope (FEI) equipped with K2 summit direct detector (Gatan) using a Gatan 626 single-tilt holder. Replicate grids from the same preparation were either imaged at our own facility (University of Chicago) or shipped to the National Cryo-Electron Microscopy Facility at the National Cancer Institute (NCI) and Case Western Reserve University (CWRU). Grids were imaged on a Titan Krios with K2 detector (super-resolution mode) and GIF energy filter (set to 20 eV) at a nominal magnification of 130,000 corresponding to a super-resolution pixel size 0.5315 Å, 0.55 Å, 0.56 Å per pixel depending on the default set up at above mentioned EM facilities, respectively. Movies were acquired at 1e^−^/A^2^ per frame for 50 frames.

### Single particle Cryo-EM analysis

All the structure determination steps were done in Relion (*4*). All the movies were binned by 2 and motion corrected by Motioncor2 (*5*). CTF estimation was done using CTFFIND4.1 (*6*). A total of 2,000 particles were manually picked and classified in 2D to generate autopicking templates. We used either SPHIRE-crYOLO package (*7*) or Relion’s built-in reference based auto picker for particle picking and the coordinates were fed into Relion for particle extraction. For each datasets, we picked between 1,000,000-5,000,000 initial particles, which were subjected to 2D classification. Some ∼ 150,000-400,000 particles were selected from good classes depending on the dataset. Between 130,000 to 250,000 of these particles were used to generate an initial model with C1 (only for the very first obtained density for prestin) and in other cases C2 symmetry imposed. All particles were then subjected to 3D refinement with C2 symmetry, yielding a 4.8 Å nominal resolution map for Up (Cl^−^) state. Classification of the particles in C1-symmetry closely resembled the overall architecture of the C2-symmetry-imposed map, albeit with lower resolution. Postprocessing of the focused transmembrane map was performed using the star file of the K2 detector at 300 kV and a masked nominal resolution of 3.3 Å by 0.143 Fourier shell correlation (FSC) criterion was calculated for Up (Cl^−^) state (*8, 9*). The nominal resolution for other states were 3.8, 3.7, 4.2, 6.7 and 4.6 Å for Inhibited I, Inhibited II, Down I, Down II and Intermediate states, respectively. After a subset of particles (between 110,000 to 180,000 depending on the state) were identified for the final refinement, the particles underwent per particle CTF refinement followed by Bayesian polishing. A final 3D refinement followed by a posprocessing step using a tighter mask and by imposing C2 symmetry. Local resolution was calculated by ResMap (*10*).

### Model building and molecular visualization

For the very first dolphin Prestin model, Swiss-Model (*11*) was used to generate a homology model based on murine and human SLC26A9 template structures (*12, 13*). The homology model was then mutated to poly alanine using Chainsaw (*14*), and all loops were subsequently deleted. Then the secondary structural elements were rigid body fit to the density. We used the 3.3 Å density map (Up) for the initial model building. We pursued the rest of model building manually and in Coot (*15–17*), registered secondary structural elements using bulky residues (e.g., Phe, Arg) and built loops where appropriate. The density was of sufficient quality to assign rotamers for key residues. Side chains of residues that could not be assigned even tentative rotamers were stubbed at the Cβ. Models were refined in real space without secondary structure restraints using phenix.real_space_refine (*18, 19*). Strong non-crystallographic symmetry constraints in phenix.real_space_refine were used to immobilize the domain that was not currently being refined (that is, the cytosolic domain during the transmembrane domain-focused map refinement). Several iterations of manual refinement and global refinement using Phenix and Coot were performed after visual screening. The dimer model was generated by applying C2 symmetry operations to the monomer in UCSF Chimera and ChimeraX (*20–22*). The initially built model was used as template for building the models for our other five EM densities. A primary fit was made using Cryofit2 tool in Phenix. Then the resulting structures were subjected to several round of refinement in Phenix and Coot. The EM density map used in modeling TM6-TM7 kink in Inhibited II state were not postprocessed or sharpened, as the kink is more defined before any postprocessing step. The sidechains and other parts of this model were fit to the postprocessed map. For models containing Salicylate, a PDB of salicylate was imported into Coot and fit to the density as a ligand. Molecular visualization and analyses performed with UCSF Chimera, ChimeraX and VMD (*23*).

### Patch-clamp electrophysiology

Adherent HEK 293 cells were used for all heterologous expression experiments. HEK 293 cells were plated for 24 hours before transient transfection. Then, 3-3.5 µg prestin plasmids and 0.4 µg eGFP plasmid were transfected to HEK 293 cells using 10 µl of Lipofectamine ® 3000 (reagent 2:1 ratio; ThermoFisher Scientific) in 500-ml Opti-MEM (Life Technologies). Cells were transferred from 37 °C to 30 °C after 20-24 h incubation to boost the expression. After 24 to 48 hours of incubation, successfully transfected cells were used for NLC measurements.

The membrane capacitance was measured in whole cell configuration, using sine wave stimulus of frequency 1 kHz and 10 mV of amplitude. Applied during voltage steps 10 ms after the transient response. Voltage steps varying from −140 to 140 mV with holding potential of −70 mV. The admittance (Y(*ω*)) of the system was calculated by spectral analysis and the DC conductance (b) was obtained from the steady state current before the sine wave stimulus. The circuit components: the capacitance (C), membrane resistance (Rm), and series resistance (Rs), were calculated is it follow (*1*):

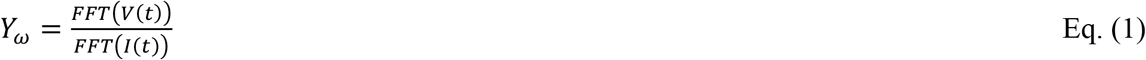

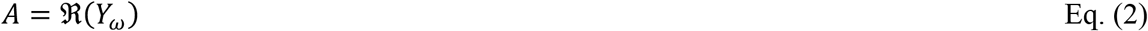

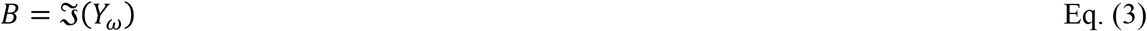

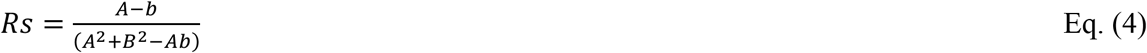

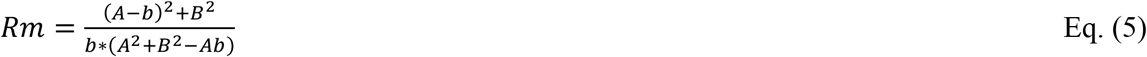

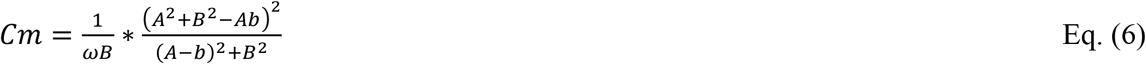

The membrane capacitance was fitted to the derivative of a Boltzmann function plus a lineal component:

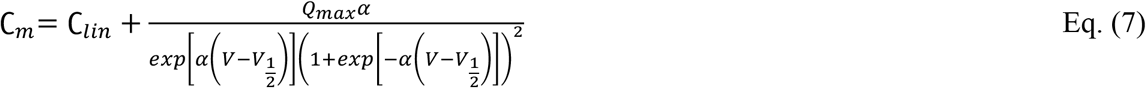

where, *Q_max_* is maximum charge transfer in response to voltage stimulation; *V_1/2_* is the voltage at which the maximum charge is equally distributed across the membrane, or in other words the peak of the voltage-dependent capacitance; *C_lin_* is the linear capacitance which is proportional to the surface area of the cell. α is the slope factor of the voltage dependence of the charge transfer,

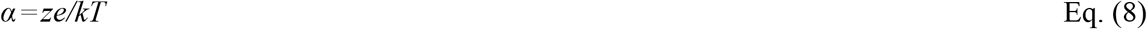

where *k* is Boltzmann’s constant, *T* is absolute temperature, *z* is valence of charge movement, and *e* is electron charge.

the NLC was obtained subtracting the linear component (*NLC= C_m_-C_lin_*). the NLC by C_lin_ (*NLC/C_lin_*) to compare the magnitude of NLC obtained from different cells with different levels of Prestin expression. The charge movement was also normalized to C_lin_, hence has the units of femtocoulomb per picofarad (*Q_max_/C_lin_*, fC/pF). We used our custom written MATLAB codes for data processing and fitting. For salicylate inhibition recordings, first, the NLC was measured then 10 mM Na-salicylate were added to the bath; after 3 minutes the NLC measurement was repeated.

Borosilicate glass pipettes were pulled using Sutter micropipette puller (P-1000, Flaming/Brown). The resistance of the capillary pipettes was from 1.6 to 2.4 megohms. The current was amplified with an Axopatch 200B amplifier (Molecular Devices, Sunnyvale, CA, USA), the data acquired at 5 kHz with a Digidata 1440A (Axon instruments) interface using pCLAMP 10 acquisition software (Molecular Devices). The Internal and external (bath) solutions are chosen such that the endogenous ionic current from HEK293 cells are minimal. Almost identical solutions are used here as also used in other Prestin studies (*1, 24*).

For the Cl^−^-based experiments, the internal solution contains 140 mM CsCl, 2 mM MgCl2, 10 mM EGTA, and 10 mM Hepes. The external solution contains 115 mM NaCl, 20 mM TEA-Cl, 5mM CsCl, 2 mM CoCl, 2 mM MgCl2, 10 mM Hepes, and 5 mM glucose. The osmolarities of the internal and external solutions were adjusted with glucose to 310 and 320 mOsm/liter, respectively and both were adjusted to pH 7.4. For sulfate based measurements, the internal solution contains 10 mM CsCl,130 mM Cs_2_SO_4_, 2 mM MgCl2, 10 mM EGTA, and 10 mM Hepes; the external solution, we used 10 Mm CsCl, 115 mM Na_2_SO_4_, 5mM Mg(OH)_2_, 20 Tris-OH, the pH was adjusted to 7.5 using 10-15 mM methanesulfonic acid. All data were acquired from at least three independent transfected cells.

### Electromotility measurements

For testing dolphin Prestin electromotility, HEK293 transiently expressing dolphin Prestin were first tested for their level of NLC, while bathed in the Cl^−^-based extracellular solution. The electromotility was followed only when the NLC was more than 0.8 pF (indicative of high Prestin expression). To better track the cell movement, cells were first lifted off the substrate using a uMp piezoelectric manipulator (Sensapex). To evoke Prestin-mediated electromotility, either voltage steps of 0.4 s, going from −140 to +150 mV in 10-mV increasing steps (e.g., Fig. 1B) or in some cases constant voltage steps of −120 to +120 mV were applied; Simultaneously the video of the cell movement were recorded using our CS2100M-USB Quantalux sCMOS camera (Thorlabs) and using 60x magnification. The videos were analyzed using a custom written Python code to track the cell movement. The maximum cellular displacement normalized by the largest diameter of the corresponding cell, *d0*, as shown in Figure 1B. These normalized values in Figure Fig. S1A are the corresponding values at change of membrane potential from +120 mv to −120 mV. The holding potential was set at −70 mV, and all the recordings were done at whole-cell configuration at room temperature. All data were acquired from at least three independent transfected cells.

### Phylogeny analyses and sequence alignments

Selected metazoan Prestin protein sequences were extracted from the complete proteomes in the NCBI Assembly database. From each proteome, one protein showing the highest BLAST bit score (*25*) to the human Prestin protein query was extracted. Sequences were aligned using MUSCLE (v.3.5) (*26*), and the ML phylogeny was inferred using RAxML (v.8.2.11) (*27*) (best-fit model of evolution: LG+G+X). The schematic representation of the phylogeny was generated using iTOL (*28*). Clustal Omega was used for the sequence alignments (*29*).

### Molecular dynamics calculations

All-atom systems were constructed using the six atomic cryo-EM structures of Prestin determined in different conditions. Orientation and position of the Prestin structures in membranes were calculated online using the web server of “Orientations of Proteins in Membranes (OPM) database” (*30*). The protein was inserted into a POPC lipid bilayer and solvated in 100 mM KCl using the VMD program (*23*). Titratable residues were assigned their default protonation state at pH 7. One additional chloride ion, sulphate ion or salicylate molecule was added into one of the two ion binding sites of Prestin. The two ion binding sites were further hydrated by water molecules according to prediction using the Dowser++ program (*31*). The resulting systems contain ∼300,000 atoms with orthorhombic periodic box dimensions of ∼180 x 120 x 150 Å^3^ and were electronically neutral.

All the simulation systems were initially energy minimized for 5,000 steps, and then equilibrated for 20 ns with gradually decreasing positional restraints being applied to the protein heavy atoms. Each system was further simulated for 150 ns under the NPT ensemble to investigate the morphology of lipid molecules surrounding the protein, the backbone atoms of which were harmonically restrained with a force constant of 1 kcal/mol/Å^2^. Two representative systems (Occluded (Cl^−^) and Inhibited II (SO_4_^2−^ and salicylate)) were simulated for another 1 μs to increase statistical significance using the special-purpose supercomputer ANTON2 (*32*). To calculate the electrostatic potential of the central ion binding sites, we performed an extra 50 ns simulation under the NVT ensemble for each system, after the 150 ns NPT run, with the same positional restraint being applied to the backbone atoms of the protein. To calculate the fraction of membrane potential, the same simulations were performed under an additional transmembrane voltage of 500 mV or −500 mV. To estimate the gating charge corresponding to the conformational change of apo-Prestin, we constructed five new systems based on the Occluded (Cl^−^) system by changing the protein from Up (Cl^−^) to the other five conformations using 20 ns targeted MD simulations. Thus, the final systems have exactly the same compositions, and without any ions in the central binding sites. Each system was then simulated for 50 ns under the NVT ensemble at −500 mV, −100 mV, 100 mV and 500 mV, with the backbone atoms of the protein being harmonically restrained with a force constant of 1 kcal/mol/Å^2^ and the ions in the solution being prevented from entering the cavity of the protein using the tcl force plugin.

The MD simulations other than the ANTON2 simulation were carried out using the NAMD program (*33*) with a time step of 2 fs. The CHARMM36 force field with torsional backbone corrections (*34, 35*) was used for protein, lipids and ions and the TIP3P model (*36*) for water in all the simulations. The GAMMP program was used to parameterize the all-atom force field for salicylate and sulphate anions (*37*). In NAMD, the temperature and pressure were controlled at 300 K and 1 atm, respectively, using the Langevin dynamics and the Nose−Hoover Langevin piston method (*38, 39*). The van der Waals interactions were smoothly switched off at 10−12 Å. The long-range electrostatic interactions were calculated using the particle mesh Ewald (PME) method (*40*). In ANTON2, the Nose–Hoover thermostat and the semi-isotropic MTK barostat (*38, 41*) were used to control the temperature and pressure, respectively. The *k*-space Gaussian split Ewald method (*42*) was used to calculate the long-range electrostatic interactions.

### Electrostatic potential and fraction of membrane potential calculations

The electrostatic potential maps were calculated using the PMEPOT plugin (*43*) of VMD. Snapshots (*n* = 4,000) from the last 40 ns trajectory of each system run under the NVT ensemble were used for the calculation. The time-averaged three-dimensional electrostatic potential maps were then used to compute the two-dimensional potential in the *x*-*z* plane (*44, 45*) crossing the two central ion binding sites. Snapshots (*n* = 4,000) from the last 40 ns trajectories of the systems run under a transmembrane voltage of +500 mV and −500 mV were used for the calculation of the fraction of membrane potential drop. The time-averaged three-dimensional electrostatic potential maps were then used to compute the fraction of membrane potential in the *x*-*z* plane crossing the two central binding sites.

### Lipid bilayer morphology and cross-section areas of the protein

For area calculations, PDBs were aligned such that their *Z* axis aligns with the symmetry axis using PPM server. Based on local resolution maps residue 460-505 always are sub 3A resolution and least mobile. Hence, we align all the structures based on this region, unless otherwise specified. We kept the same criteria for SLC26A9 for comparison. CHARMM-GUI (*46*) membrane builder tool was used for Prestin and SLC26A9 area calculation across the membrane thickness.

### Charge displacement calculations

The gating charges of protein conformational change were estimated by calculating the average displacement charge using 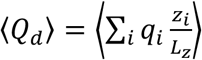 where *q*_i_ and *z*_i_ are the partial charge and the unwrapped *z* coordinate of atom *i*, respectively, and *L*_z_ is the length of the simulation box along the *z* direction (*3, 44, 47*). Snapshots (*n* = 4,000) from the last 40 ns trajectories at different trans-membrane voltages of each system were used to calculate the average displacement charges. The calculated average displacement charges of each system were then linearly fitted together, and the offset constants correspond to the gating charge changes between different conformations.

### Calculation of cell motility based on Prestin’s cross-sectional expansion

We assumed that all the Prestin dimer is aligned along the lateral line of the OHCs and they are cooperative; also, that the OHC membrane fully follows the Prestin’s footprint and only the lateral wall of OHC extends due to Prestin cross-sectional expansion. For ease, it is also assumed that OHCs have a perfect cylindrical shape.

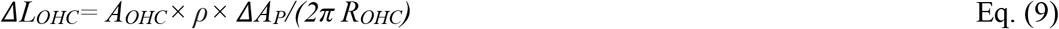

Where:

*ΔL_OHC_*, Maximum possible somatic motility of the OHC based solely based on collective cross-sectional expansion of Prestin molecules in the lateral wall.

*A_OHC,_* Average lateral area of an OHC.

*ρ*, Prestin density (prestin counts per µm^2^)

*ΔA_Prestin_*, Average cross-sectional area expansion of a Prestin dimer from Down to Up (Cl^−^).

*R_OHC_*, Average circumferential radius of OHC.

*L_OHC_, Average length of OHC*

Typical values used for above variables were, *A_OHC_= (Average length of ρ* =7000 prestin/ µm^2^, *ΔA_Prestin_*=750 Å^2^, *R_OHC_* =4 µm and *L_OHC_=50* µm (*48–50*).

Note that the OHC dimensions (length and dimeter) vary along tonotopically-defined frequency segments of the cochlea and are also species dependent. Hence, the calculations provided represent rough estimates of whether Prestin in-plane area expansion at the molecular scale could underlie somatic motility in the OHCs.

### Statistical analysis

Statistical significance (criteria: *P-value<0.005) was determined using an un-paired Student’s t Test. In electrophysiology data, this was determined by comparison to data obtained from wild-type dolphin Prestin in Cl^−^ with Wilde type Prestin in SO_4_^2−^. In order to compare electrophysiology data from the mutants with the Wild type Prestin, we used One-way Anova with a Tukey’s post-hoc test.

---

**Fig. S1.**
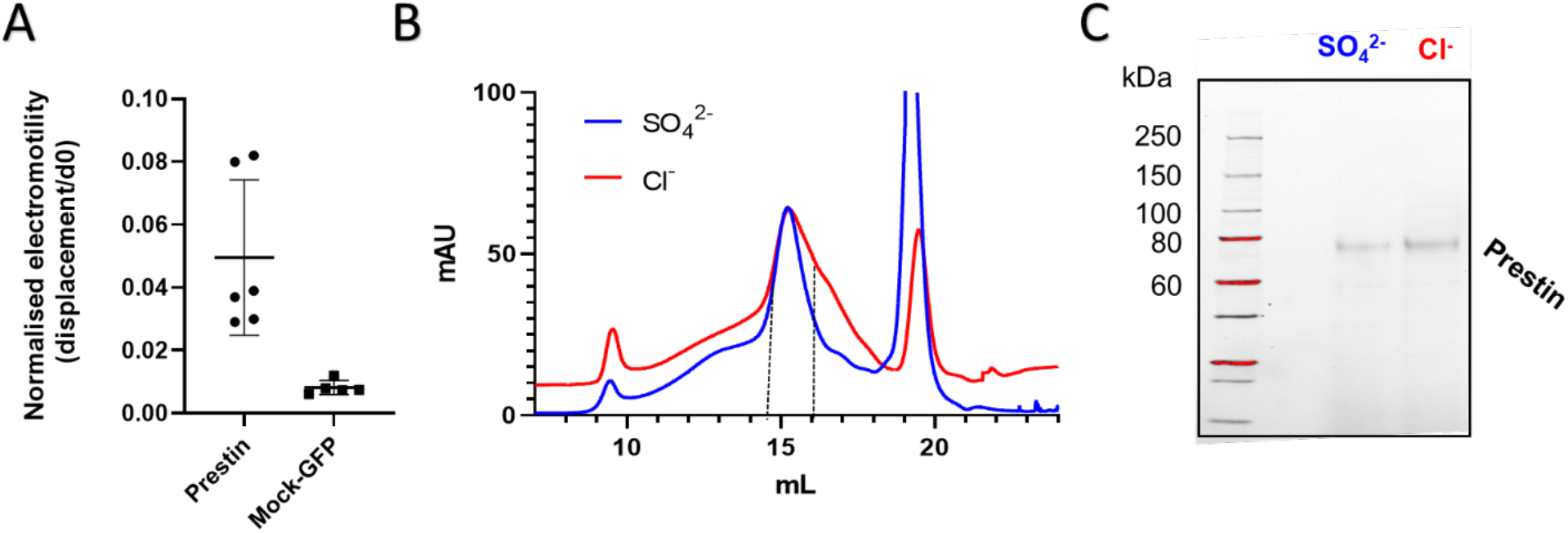
(A) Electromotility analysis of HEK 293 cells transfected with wild type dolphin Prestin compared to GFP-only transfected cell (Mock-GFP). The cellular displacement has been normalized based on the cell largest diameter, *d_0_* (Fig. 1B). The normalized electromotility was 0.05±02 versus 0.008±0.002 for wild type Prestin and Mock-GFP, respectively. These values were measured at the depolarizing voltage step changing from +120 mV to −120 mV (mean ± SD; Nonparametric Student t-test, unpaired, *P*=0.005). (B) Size exclusion chromatography (SEC) curves of the full-length dolphin Prestin purified in GDN, run on a Superose 6 column, in high Cl^−^ (red) and SO_4_^2−^ (blue) based solution. The fractions indicated by black dotted lines in both represent purified proteins that were used for cryo-EM imaging. (B) Purified dolphin Prestin cryo-EM samples, run on a Stain-free SDS-PAGE gel, indicating size of ∼ 75 kDa for the full-length Prestin monomer.

**Fig. S2.**
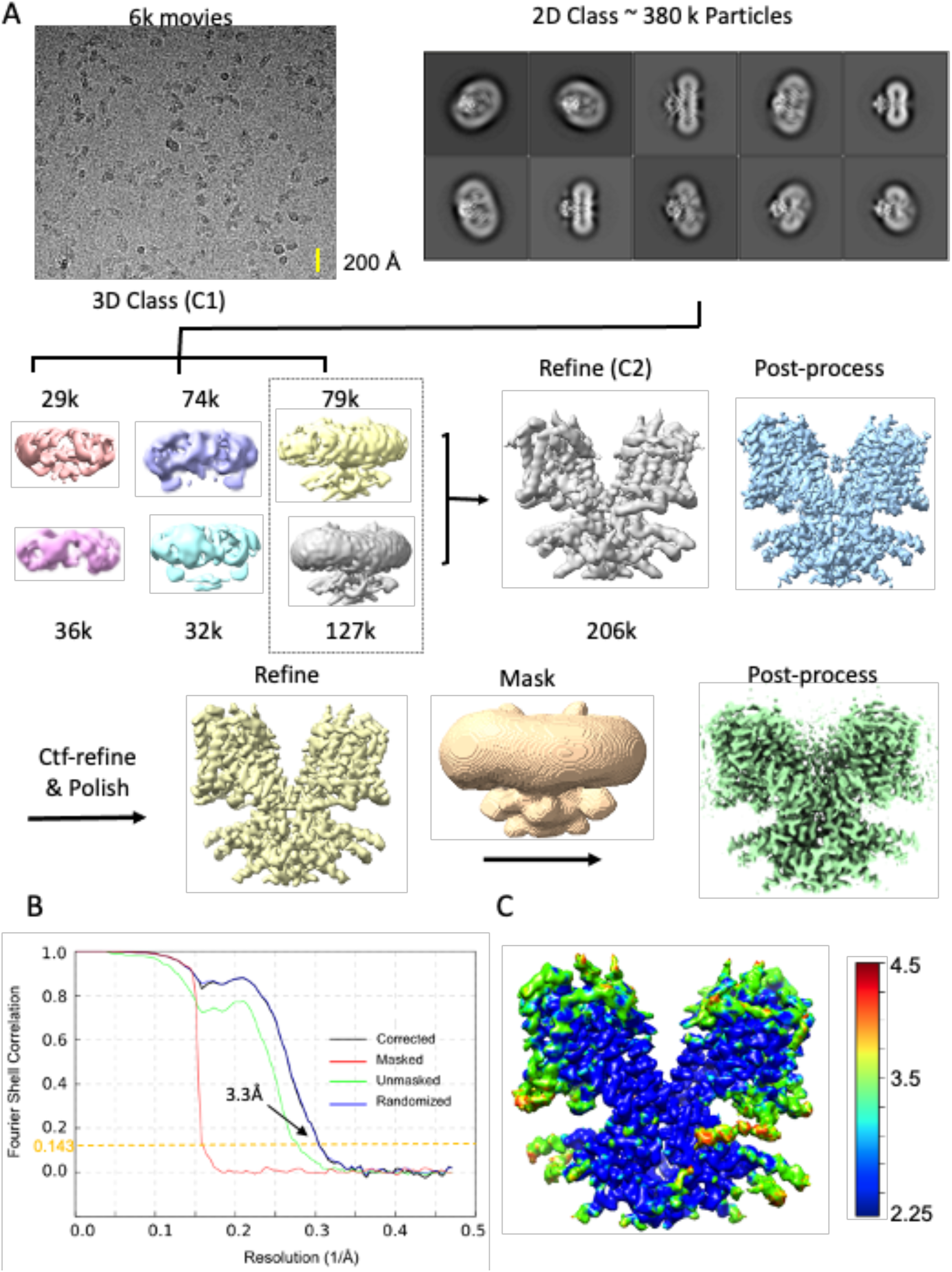
Flow chart for the cryo-EM data processing and structure determination of the dolphin Prestin in high Cl^−^ condition (See Methods for details). The final reconstruction has a normal resolution of 3.3 Å (at Fsc=0.143). All the images in this figure were created in UCSF ChimeraX.

**Fig. S3.**
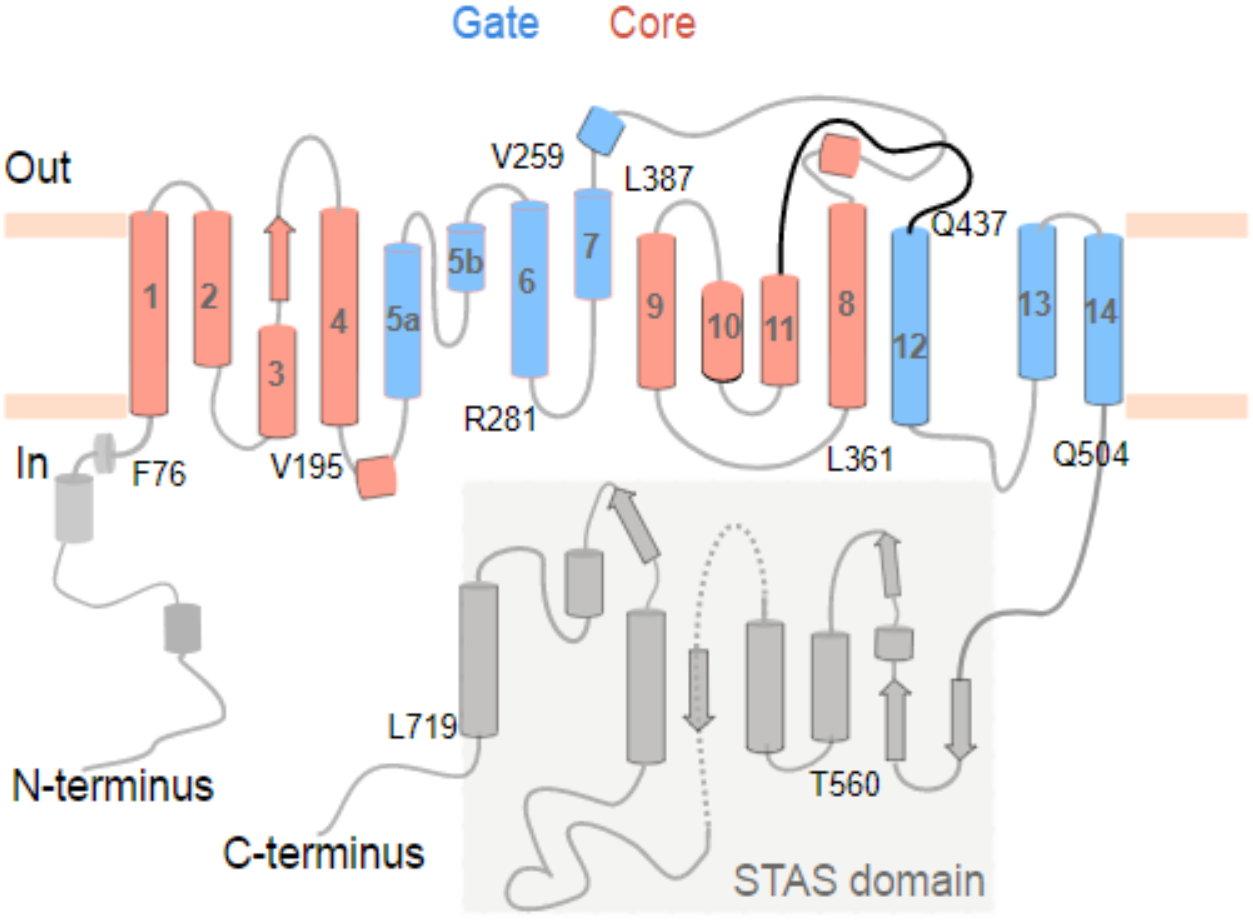
Topology of dolphin Prestin. Different domains are indicated by color; the gate domain is colored in blue, the core domain in red and the C- and N-termini as well as the STAS domain in grey. The transmembrane helices are numbered from 1 to 14. The N- and C-termini as well as the STAS domain are oriented towards the cytoplasm.

**Fig. S4.**
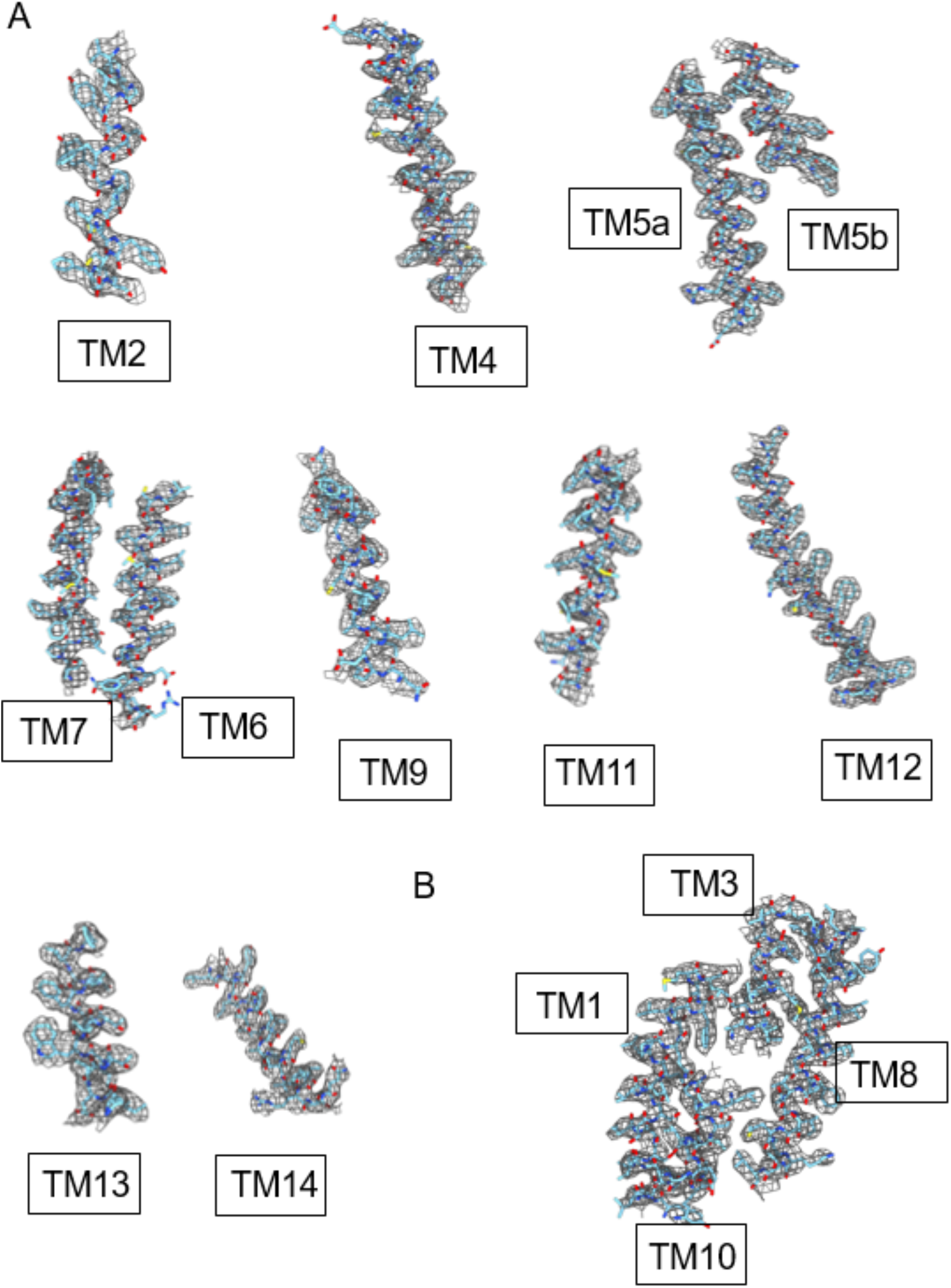
Details of the atomic model fitted to the postprocessed cryo-EM density map are illustrated for (A) the transmembrane helices including those forming (B) the anion-binding pocket.

**Fig. S5.**
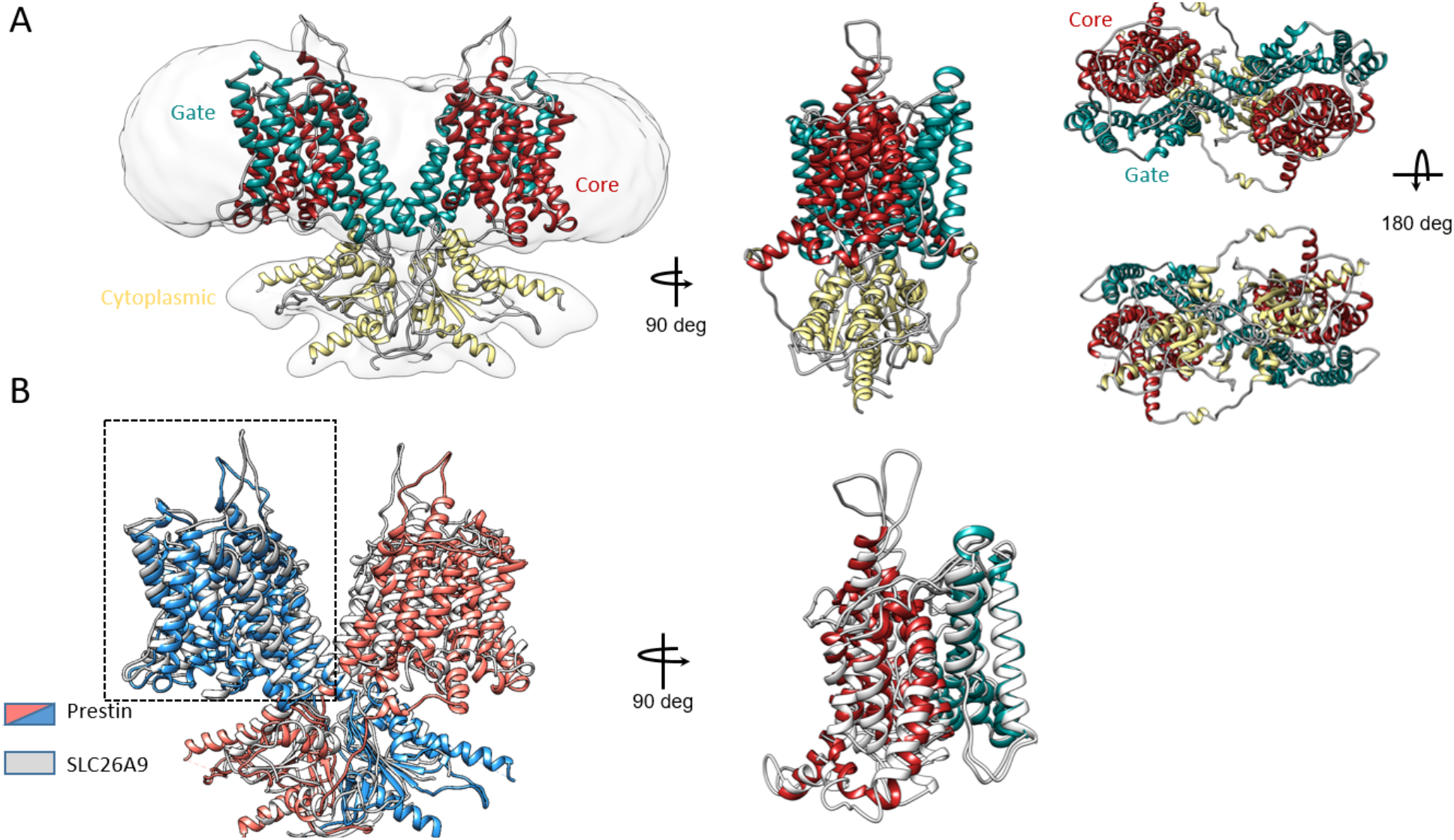
Structure of Prestin in high Cl^−^ and comparison with the Intermediate conformation of SLC26A9 (PDB:6RTF). (A) Overall architecture of dolphin Prestin at (high Cl^−^) fit to the cryo-EM density, with different domains highlighted by color. (B) Comparison between Prestin (high Cl^−^) (blue and Red) and SLC26A9 Intermediate state (grey). Subunits A (dotted box) from both proteins were aligned. The structures are aligned based on residue 460 to 505 (TM13-TM14). ChimeraX was used for illustration.

**Fig. S6.**
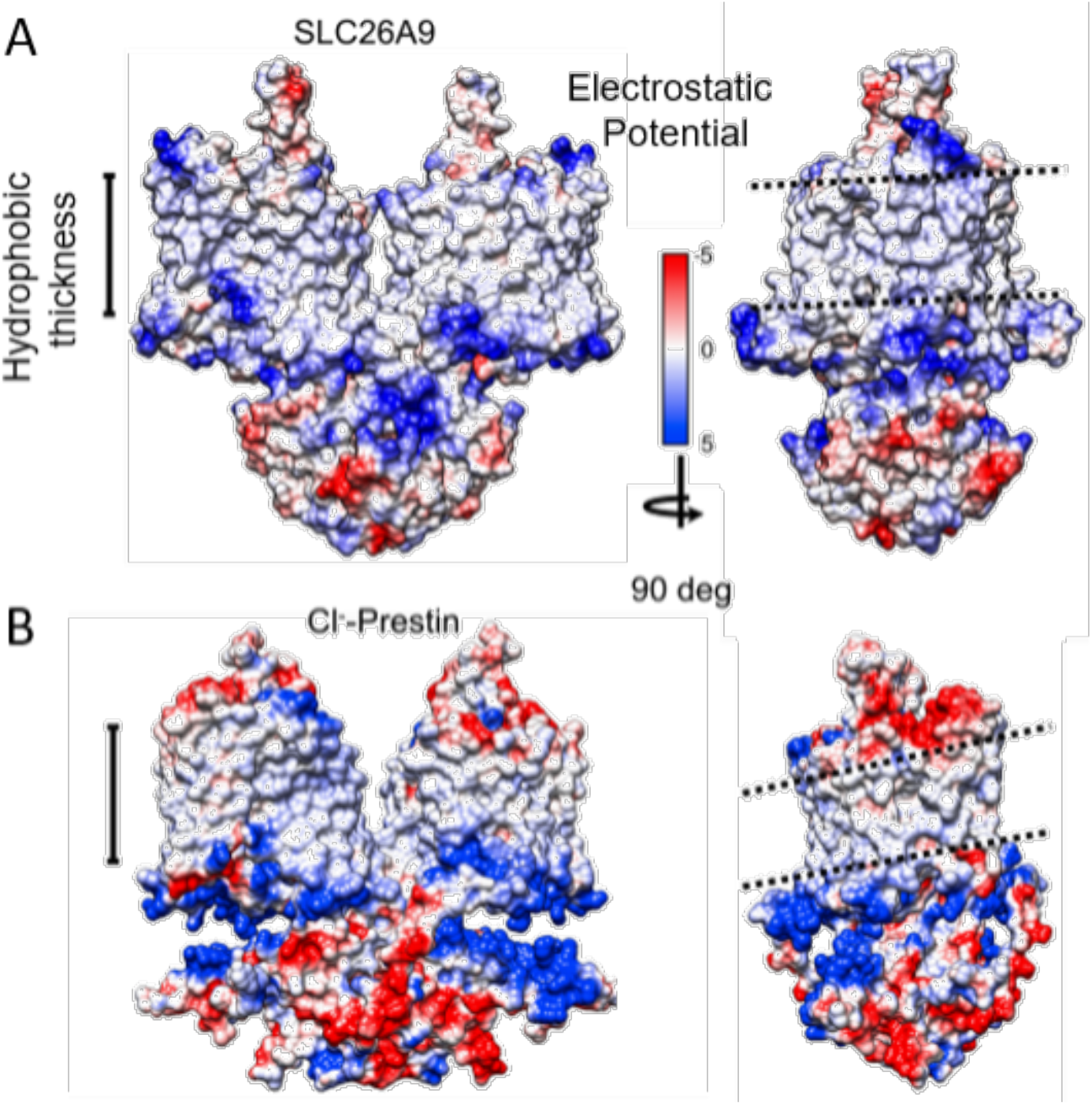
Electrostatic potential and surface charge distribution of SLC26A9 intermediate state (6RTC, panel (A)) compared with that of Prestin in high Cl^−^ panel (B). The electrostatic charge distribution ranges from −5 to 5 kT from negative to positive charge. ChimeraX was used for illustration.

**Fig. S7.**
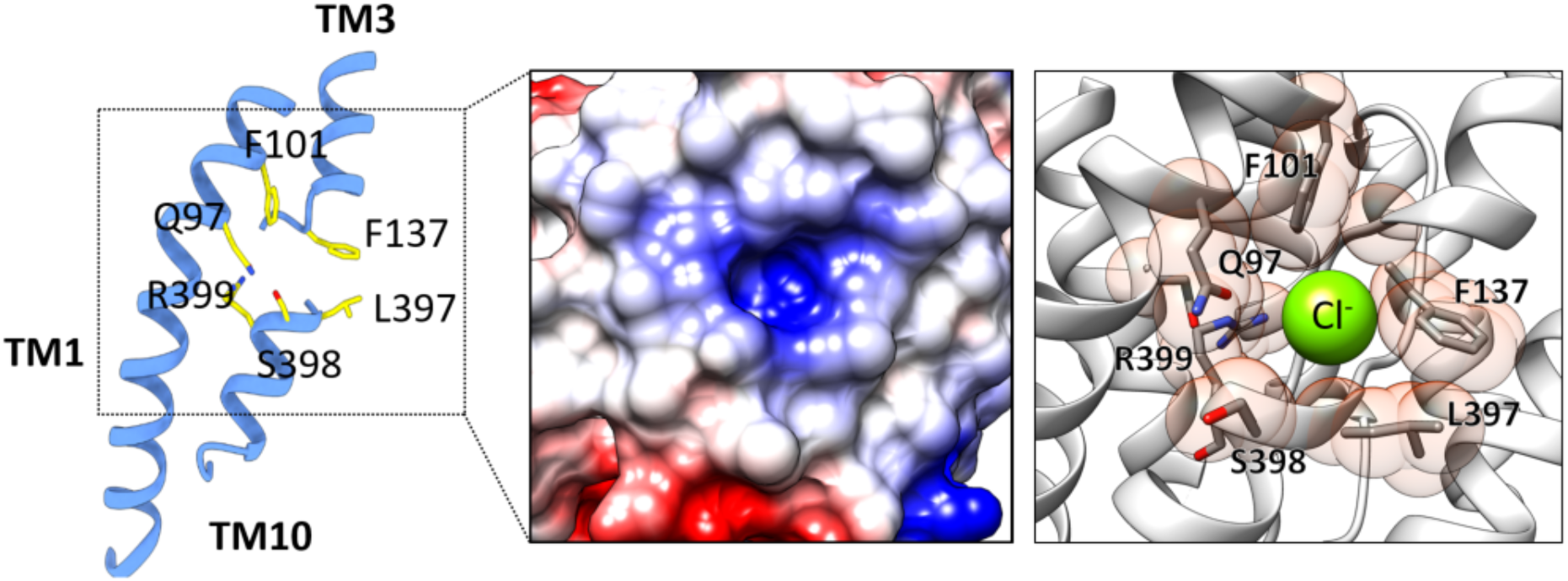
Detailed Structure of Prestin’s anion-binding site. Left panel: Key anion binding site residues Q97, F101, F137, L397, S398 and R399 are shown in yellow color and by their heteroatom. Middle panel: the electrostatic potential at this site shows a positive field (middle panel); we manually placed a Cl^−^ anion into the positive cavity (right panel). Only TM10-TM3 dipole and TM1 are shown for clarity (right panel). UCSF ChimeraX was used for illustration.

**Fig. S8.**
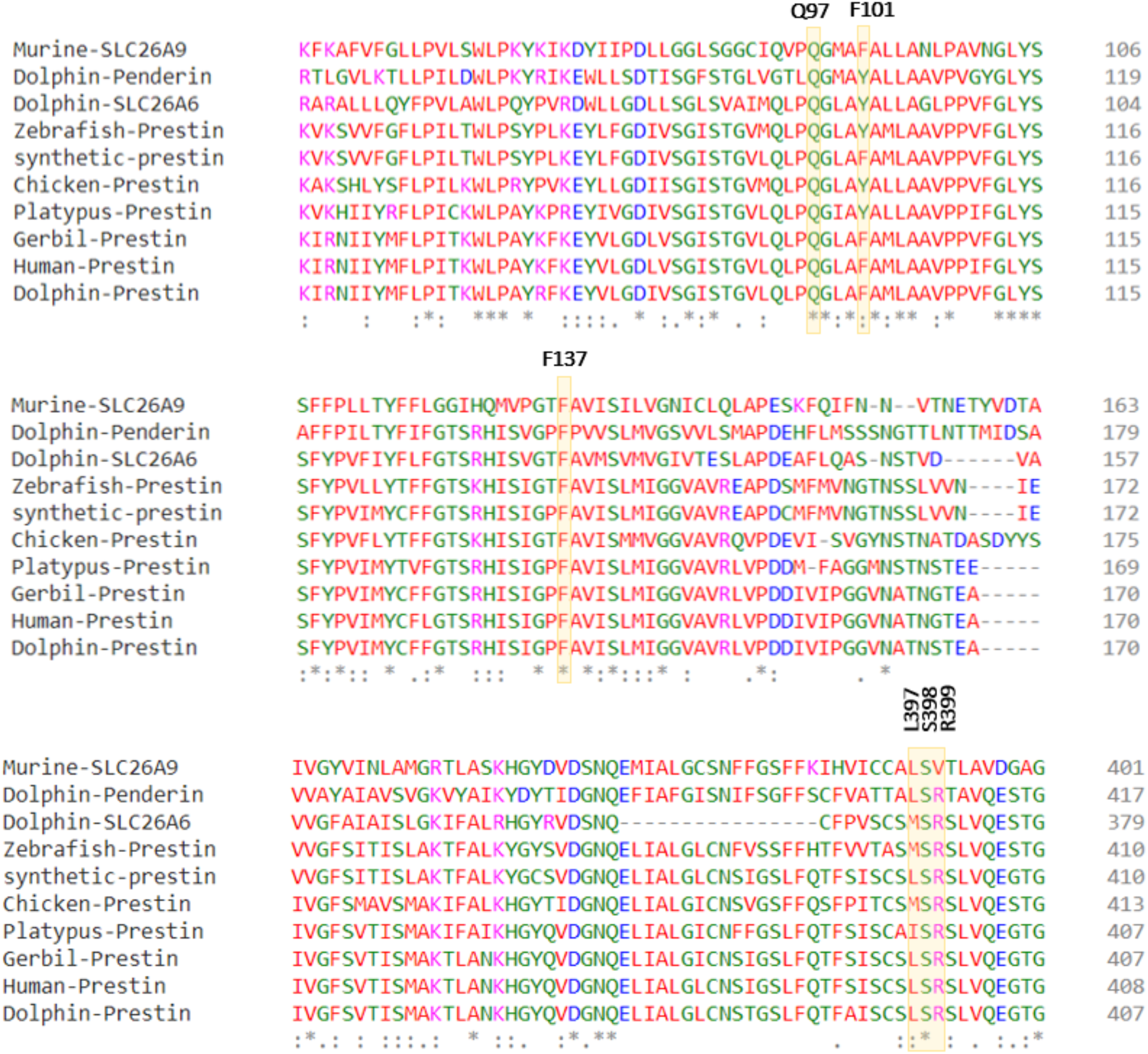
Sequence alignment of Prestin and close SLC transporters across different species. Residues forming the anion-binding site are largely conserved (e.g. Q97, F101, F137). Putative voltage-sensing residue R399 in dolphin Prestin is replaced by a valine in murine SLC26A9. Clustal Omega was used for the sequence alignments.

**Fig. S9.**
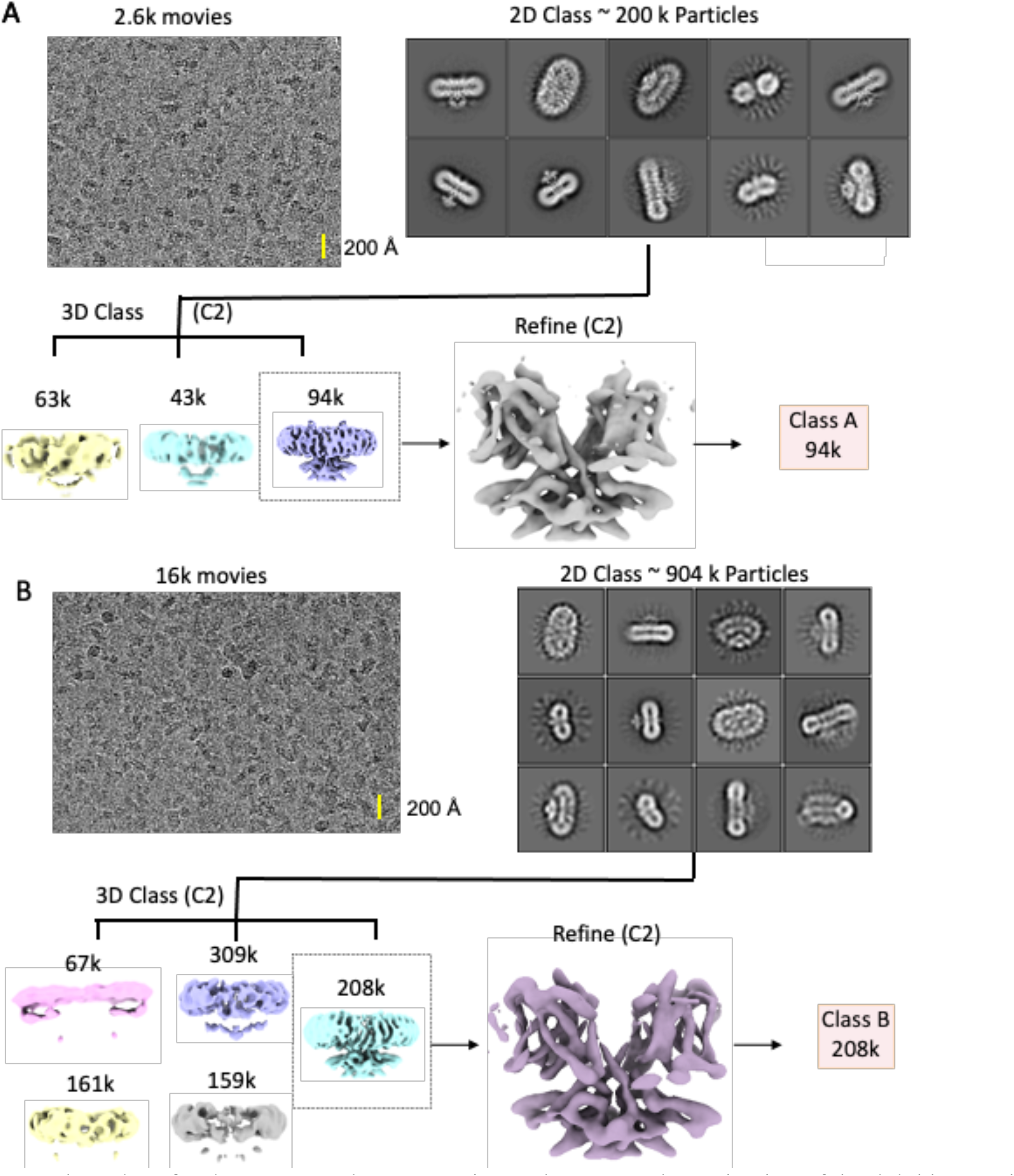
Flow chart for the cryo-EM data processing and structure determination of the dolphin Prestin in SO_4_^2−^ (see methods). UCSF ChimeraX was used for illustration.

**Fig. S10.**
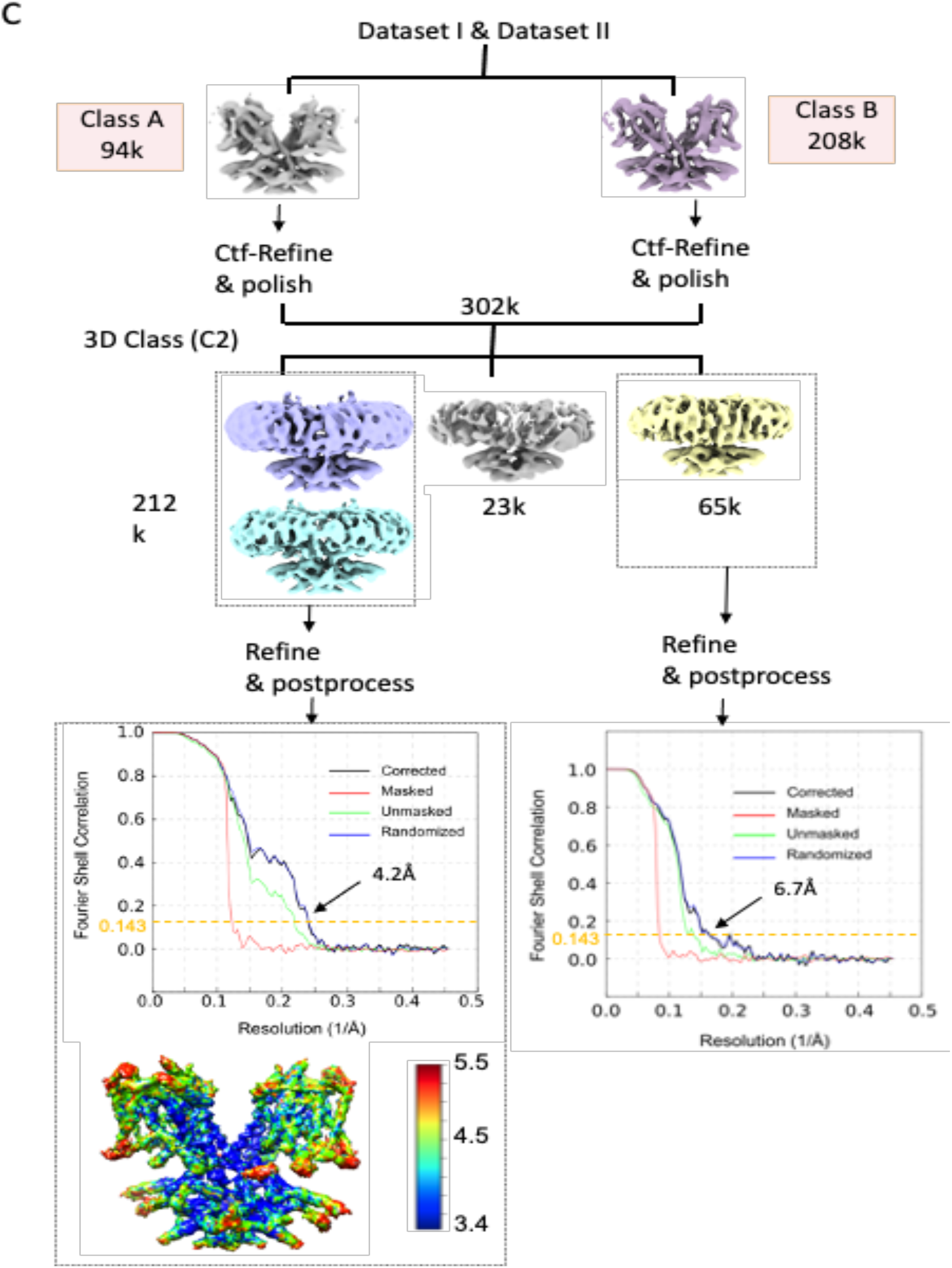
Flow chart for the cryo-EM data processing and structure determination of the dolphin Prestin in Down I (SO_4_^2−^) and Down II (SO_4_^2−^) states (See Methods for details). Class A was obtained from Dataset I, which was combined with (B) Class B from Dataset II. (C) The final reconstruction yielded two structures, Down I (SO_4_^2−^) and Down II (SO_4_^2−^), which have nominal resolutions of 4.2 and 6.7 Å, respectively (at FSC=0.143). Evidence of both states was found in dataset II, however merging of datasets was required to improve resolution of states. All the images in this figure were created in UCSF ChimeraX.

**Fig. S11.**
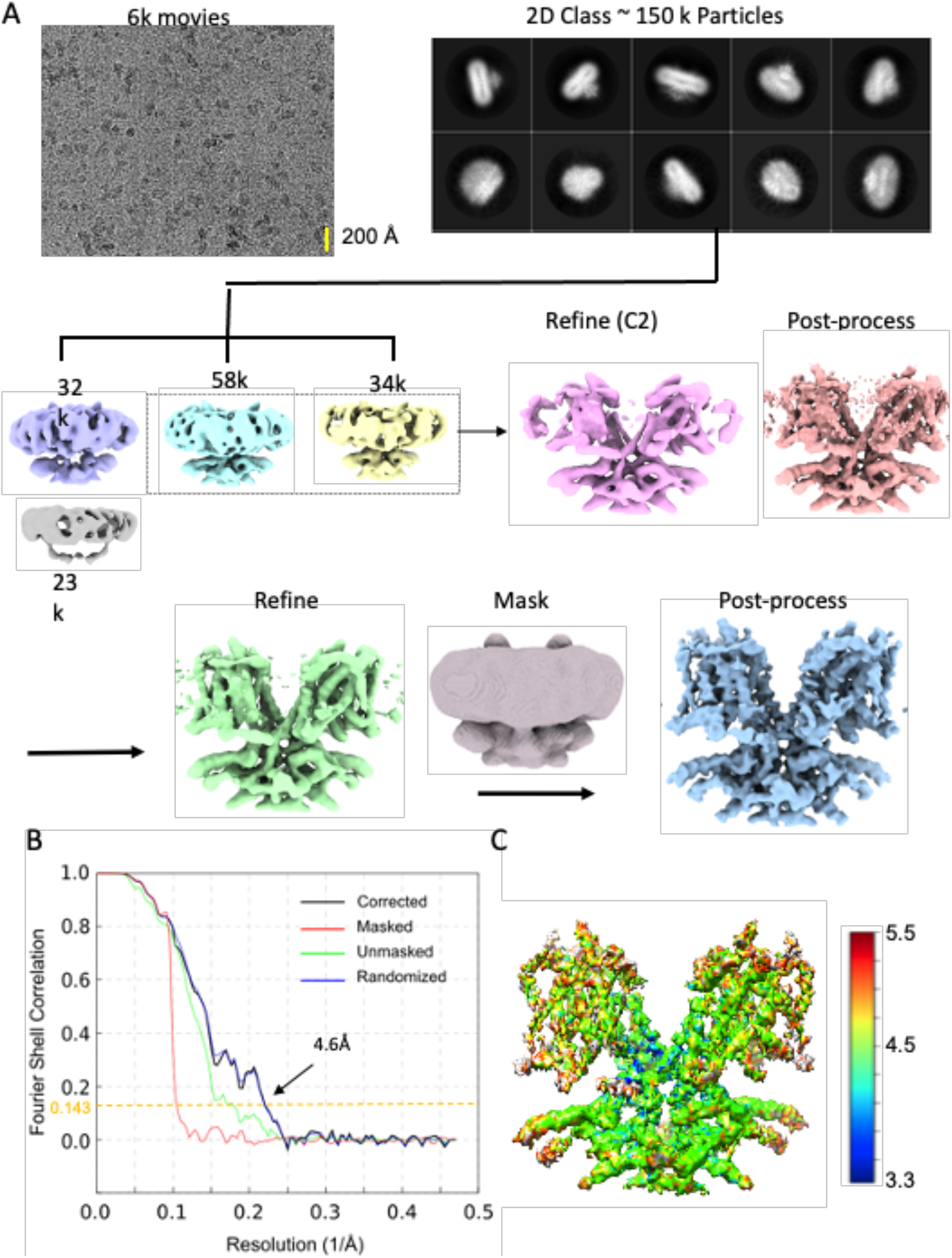
Flow chart for the cryo-EM data processing and structure determination of the dolphin Prestin in the Intermediate state (SO_4_^2−^) (See Methods for details). The final reconstruction has a nominal resolution of 4.6 Å (at FSC=0.143). All the structures were illustrated in UCSF ChimeraX.

**Fig. S12.**
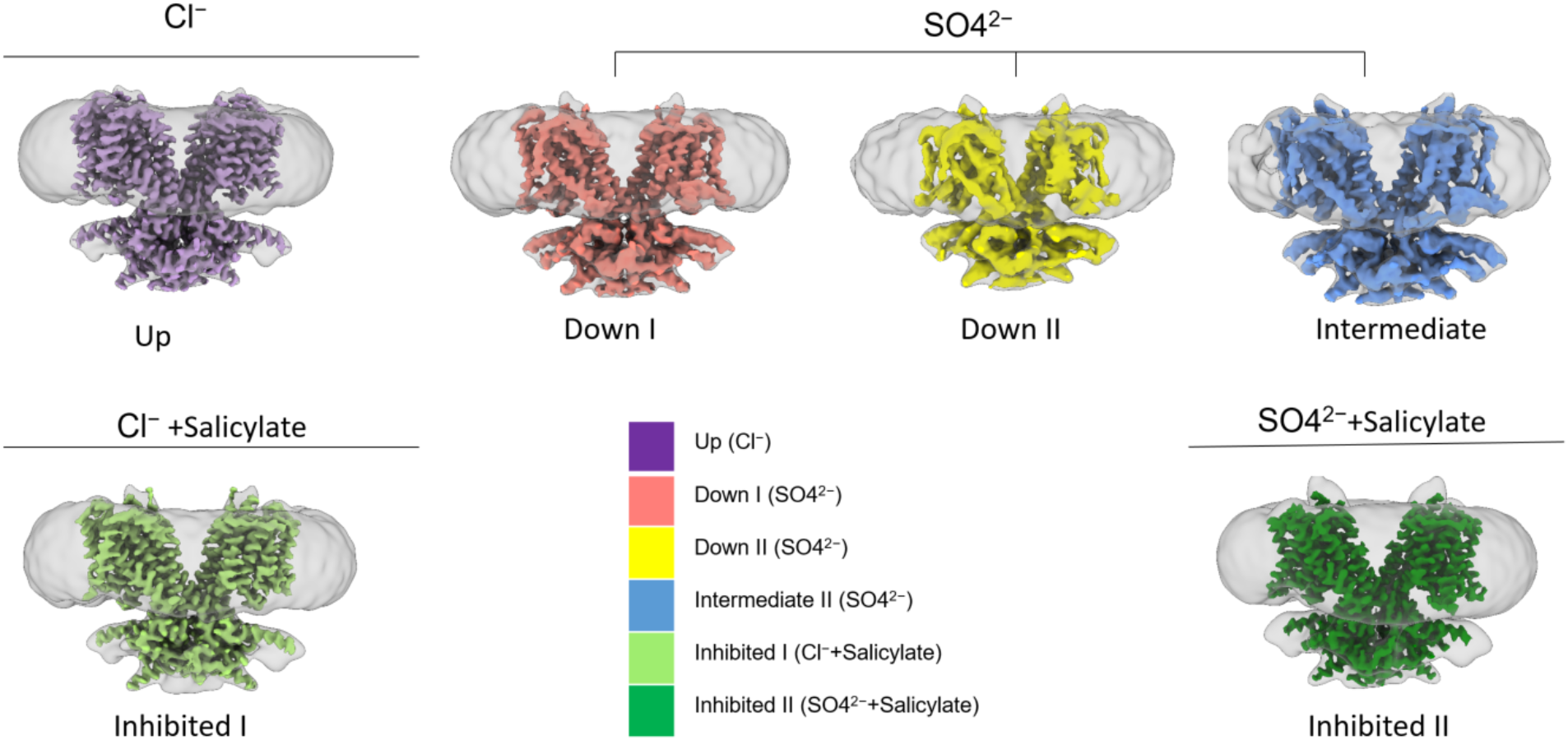
Overview of determined structures. Dolphin Prestin’s structure has been captured in six distinct states, namely: Up (Cl^−^), Down I (SO_4_^2−^), Down II (SO_4_^2−^), Intermediate (SO_4_^2−^), Inhibited I (Cl^−^ + salicylate) and inhibited II (SO_4_^2−^+salicylate). All the structures were illustrated in UCSF ChimeraX.

**Fig. S13.**
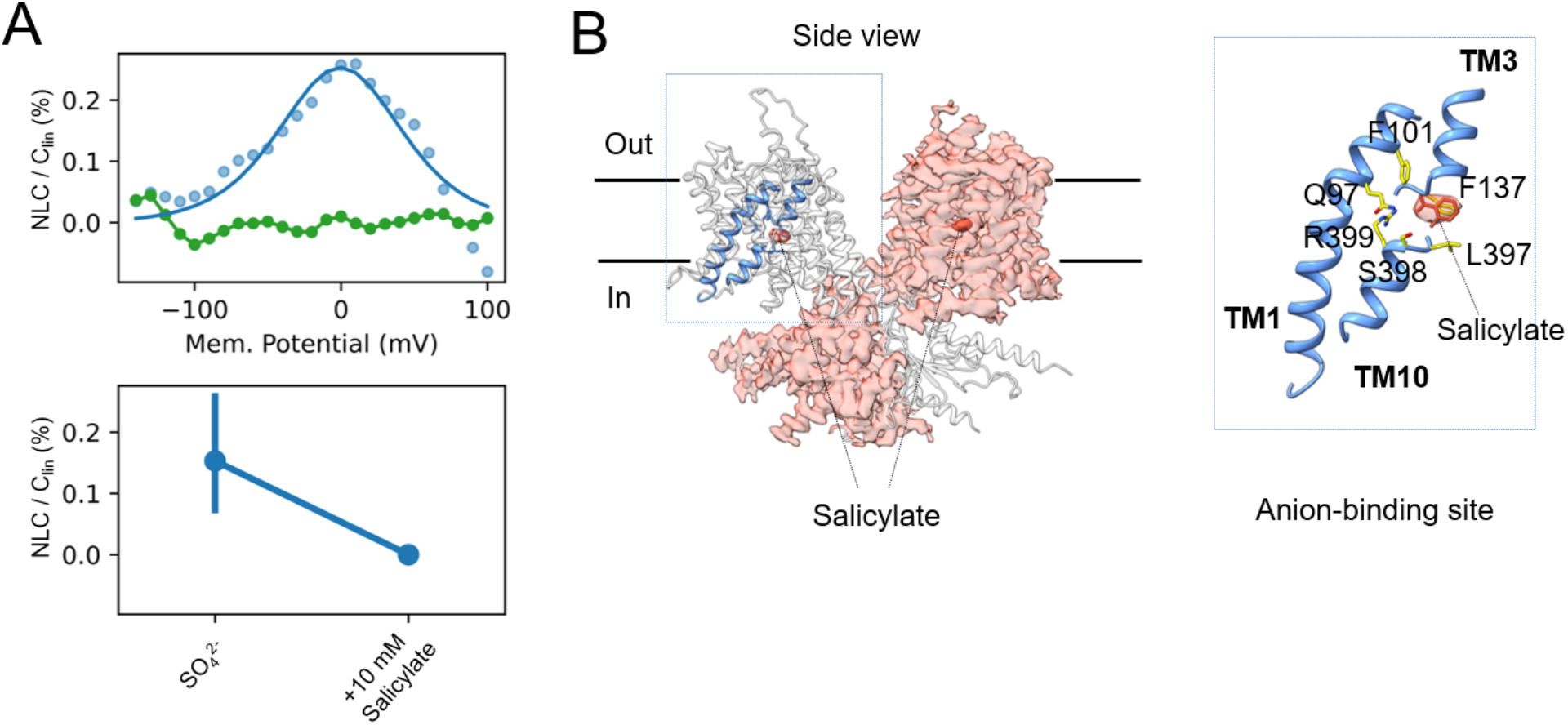
Salicylate outcompetes SO_4_^2−^ in binding to anion-binding pocket. (A) Patch clamp electrophysiology of HEK293 cells transfected with Dolphin Prestin. The NLC is abrogated by 10 mM Na-Salicylate, when SO_4_^2−^ is the main anion of the bath and pipette solutions (See Methods). (B) Density of Salicylate (orange) in the anion-binding site (blue) was resolved in the Inhibited II (SO_4_^2−^) state of dolphin Prestin.

**Fig. S14.**
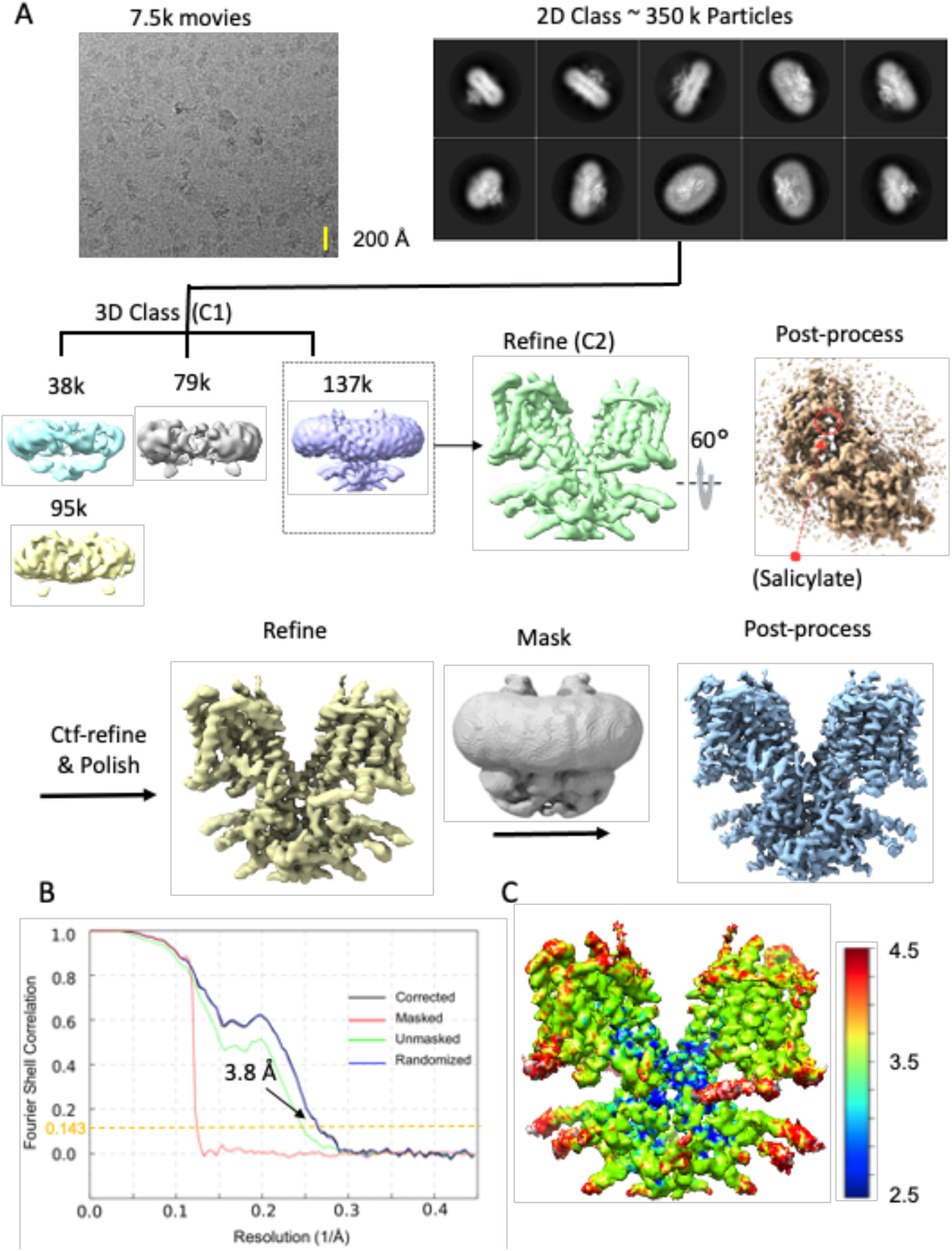
Flow chart for the cryo-EM data processing and structure determination of the dolphin Prestin in the Inhibited I state (Cl^−^ + Salicylate) (See Methods for details). The final reconstruction has a nominal resolution of 3.8 Å (at FSC=0.143). All the images in this figure were created in UCSF ChimeraX.

**Fig. S15.**
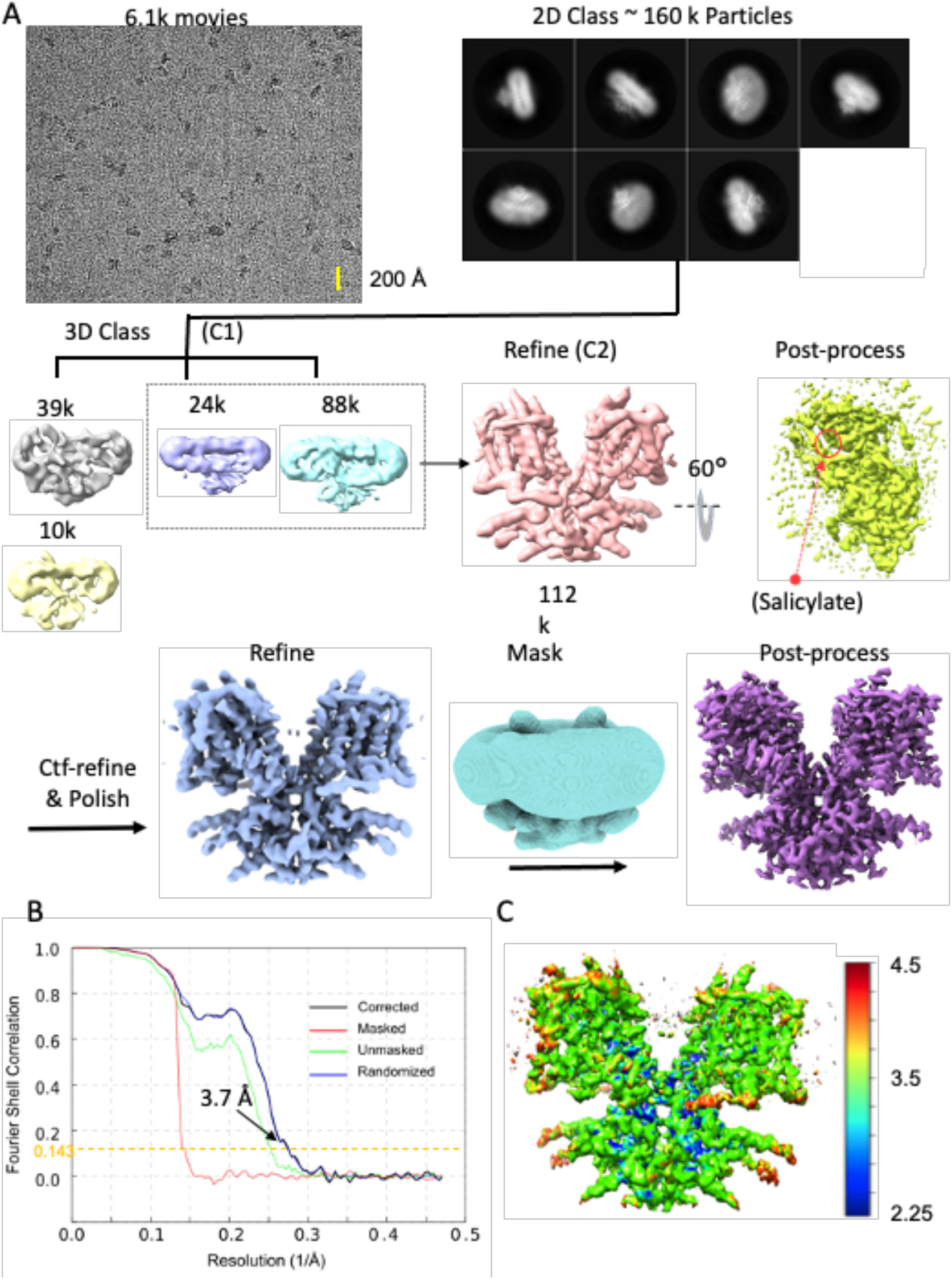
Flow chart for the cryo-EM data processing and structure determination of the dolphin Prestin in the Inhibited II state (SO_4_^2−^ + Salicylate) (See Methods for details). The final reconstruction has a nominal resolution of 3.7 Å (at FSC=0.143). All the images in this figure were created in UCSF ChimeraX.

**Fig. S16.**
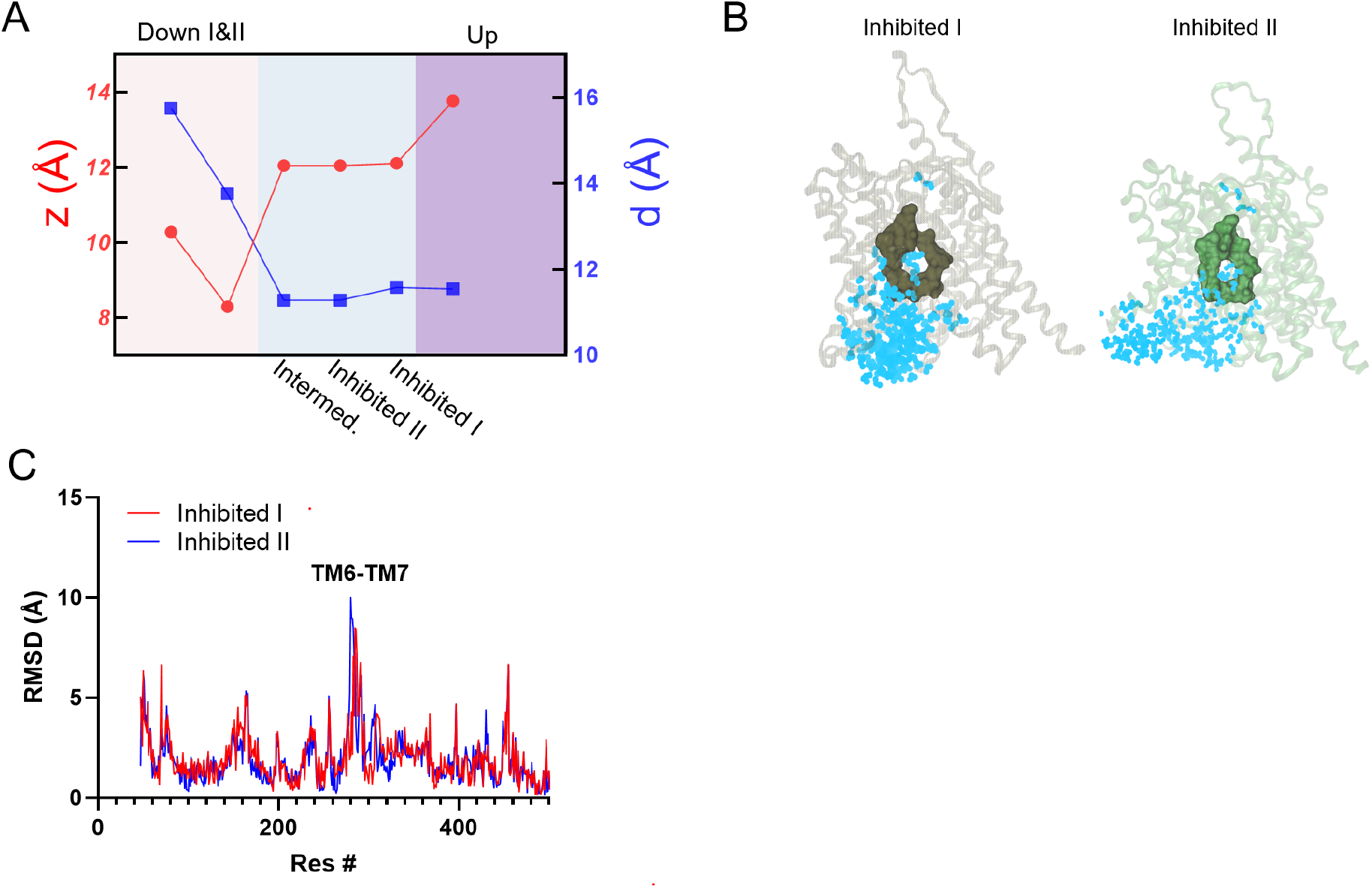
Inhibited I & II (+ salicylate) states are closest to the Intermediate (SO_4_^2−^) state. (A) The vertical distance between R399 and V499 (as a measure for the movement of the voltage sensor), *z*, and the distance between F137 and R399, *d*, are calculated across states. When comparing the Inhibited states and the Intermediate state, differences between *z* and *d* are minimal, indicating a similar conformation of the voltage sensing and anion binding sites in the Inhibited and Intermediate states. (B) MD simulations shows that the anion-binding pocket (green) in inhibited I & II is not accessible by water (cyan). The water molecules within 8 of the residues Q97, F101, F137 V397, S398, R399, E280 and E404 have been screened and illustrated using VMD. (C) RMSD calculation of Inhibited I & II (salicylate) states compared with the Intermediate (SO_4_^2−^) state. The major difference between them is in the TM6-TM7 helical dipole region as well as in the TM8 helix. The structures were aligned based on residue 460 to 505 (TM13-TM14).

**Fig. S17.**
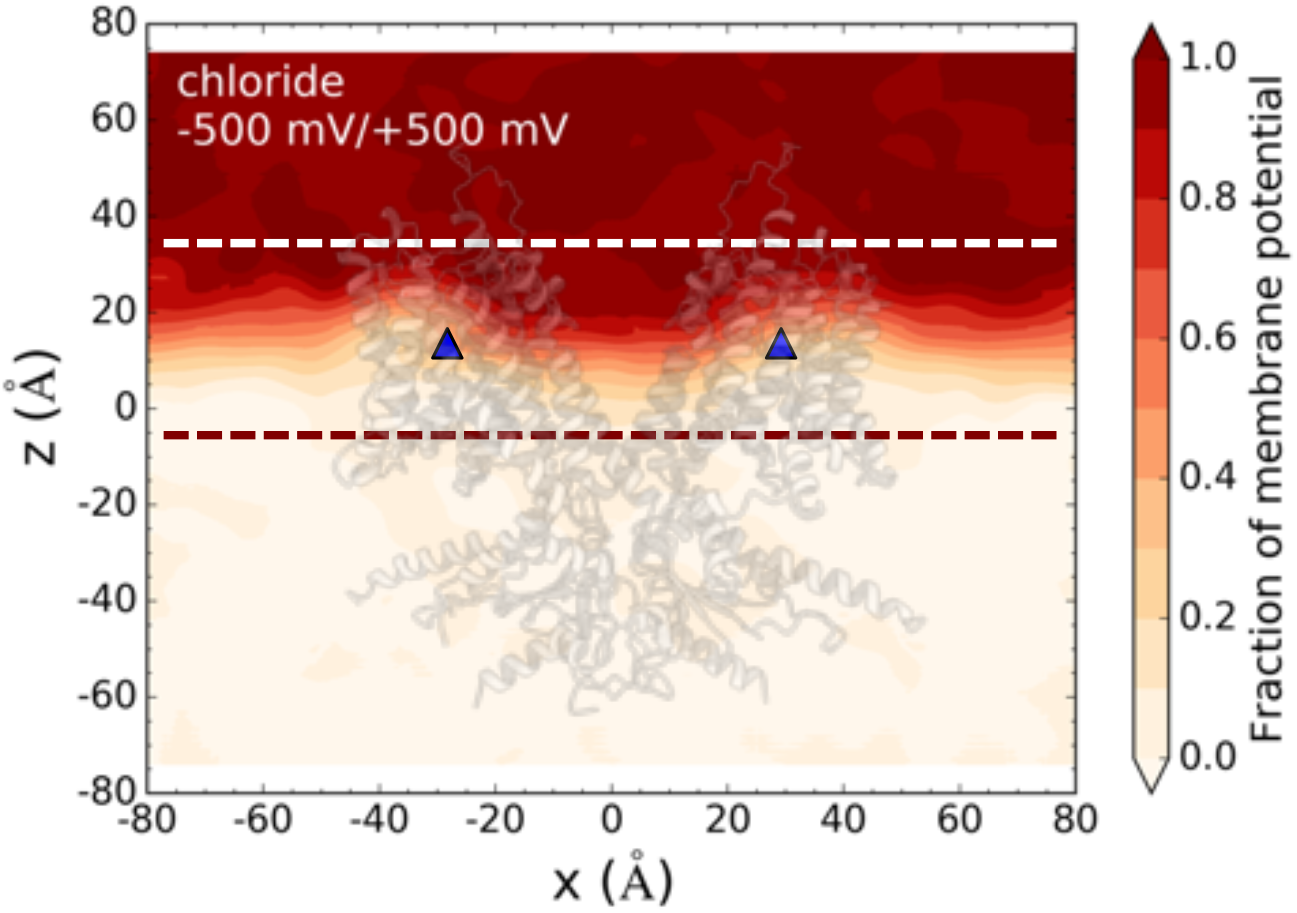
The fraction of membrane potential around the binding sites of Up (Cl^−^) state. The MD simulation box on the left and the corresponding 2-D fraction of membrane potential in the *x*-*z* plane crossing the central binding sites.

**Fig. S18.**
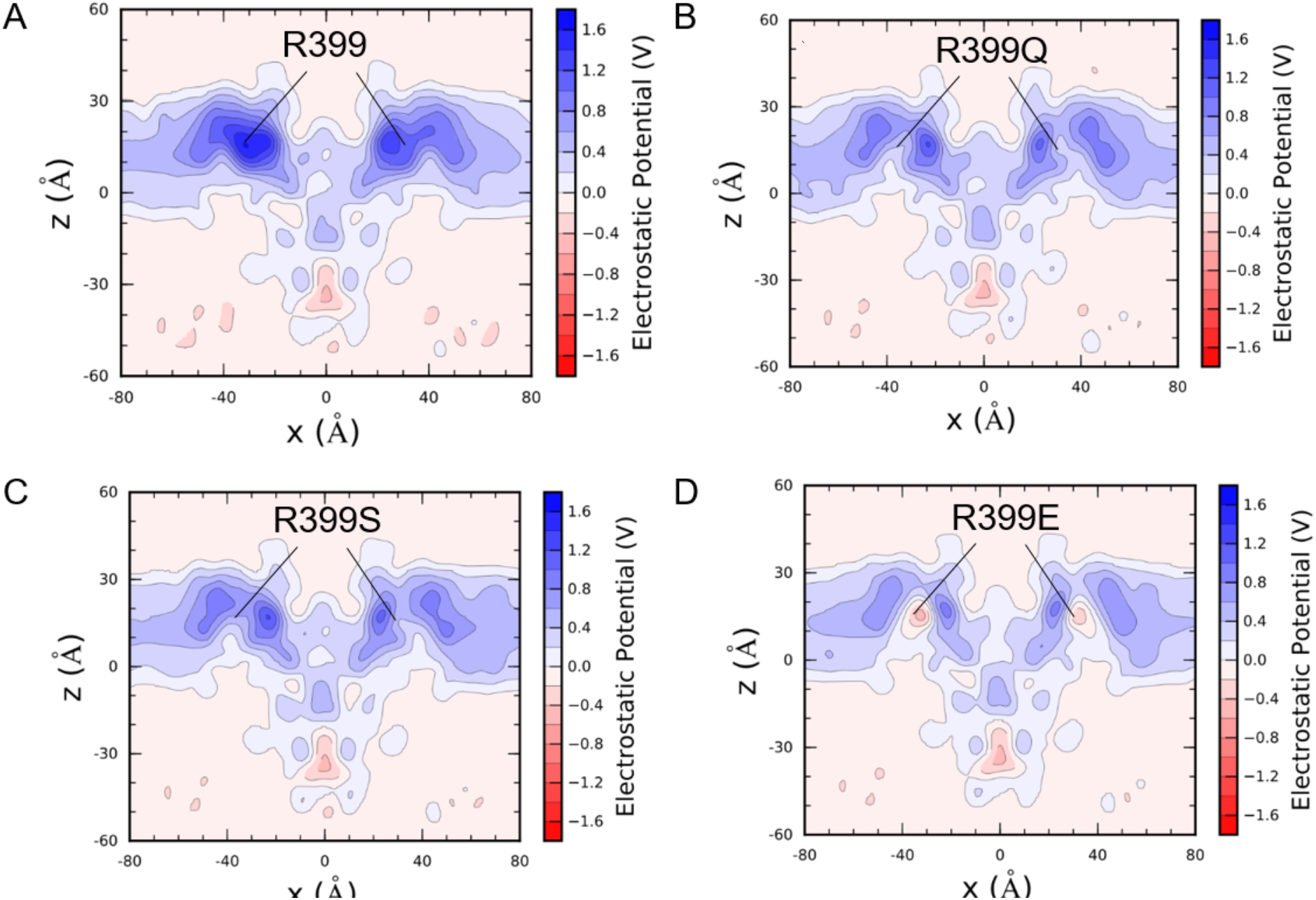
2D electrostatic calculations of Up state, without Cl^−^ bound to either of the two binding sites, based on all-atom molecular dynamics simulation. (A-D) R399 in both monomers have been mutated to Q and S and E in different systems to see the contribution of R399 residue to the positive charge at the bilayer mid-plane. R399 mutation to polar residues shows that R399 has almost ∼40% contribution the positive charge of the field at the bilayer mid-plane. The remainder likely comes from the TM3-TM10 helical dipole and other positive charges in this area.

**Fig. 19.**
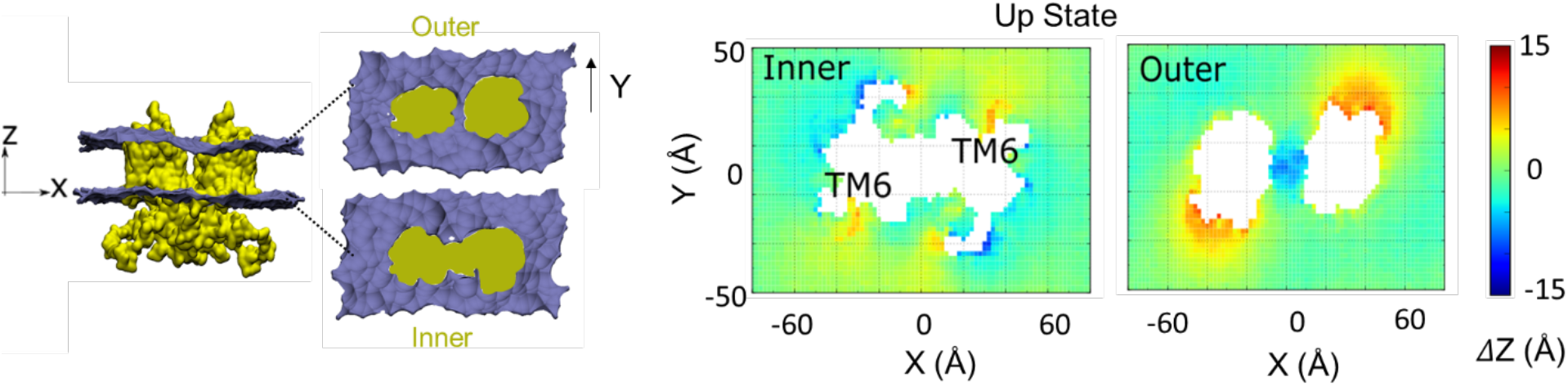
MD simulation of Prestin (Up states) equilibrated in POPC lipid bilayers. The cross-sectional area of outer and inner monolayer with mapped leaflet coordinate in the Z direction (across the membrane thickness) using all-atom molecular dynamics simulations (1µs). The comparison was made between Up (Cl^−^) and Inhibited II (SO_4_^2−^) states. The largest difference was observed at the location of the TM6 helix.

**Fig. S20.**
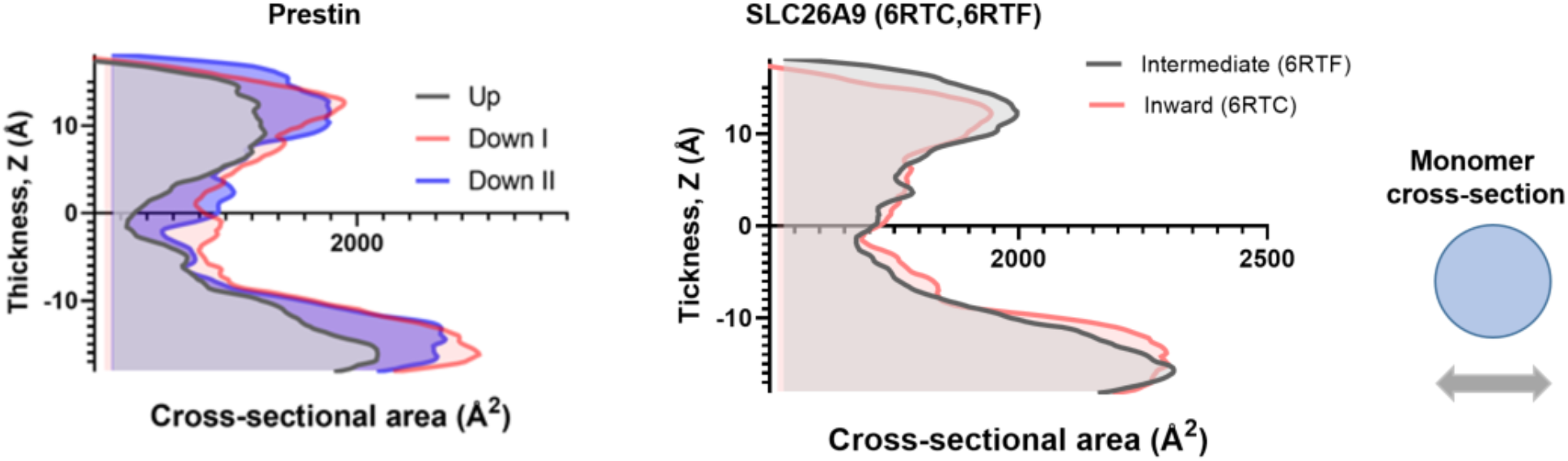
Cross-sectional area calculations of the transmembrane domain of different dolphin Prestin structures compared to their corresponding structures of SLC26A9 along the hydrophobic thickness using CHARMM-membrane builder. Cross-sectional area change from Down to Occluded in Prestin and that of SLC26A9 from Inward-facing to Intermediate states (6RTC and 6RTF) per monomer. Note that prior to area calculation, the spatial arrangements of all the structures with respect to the hydrocarbon core of the lipid bilayer were first adjusted using the PPM server. The structures were aligned based on residues 460 to 550 (TM13-TM14).

**Fig. S21.**
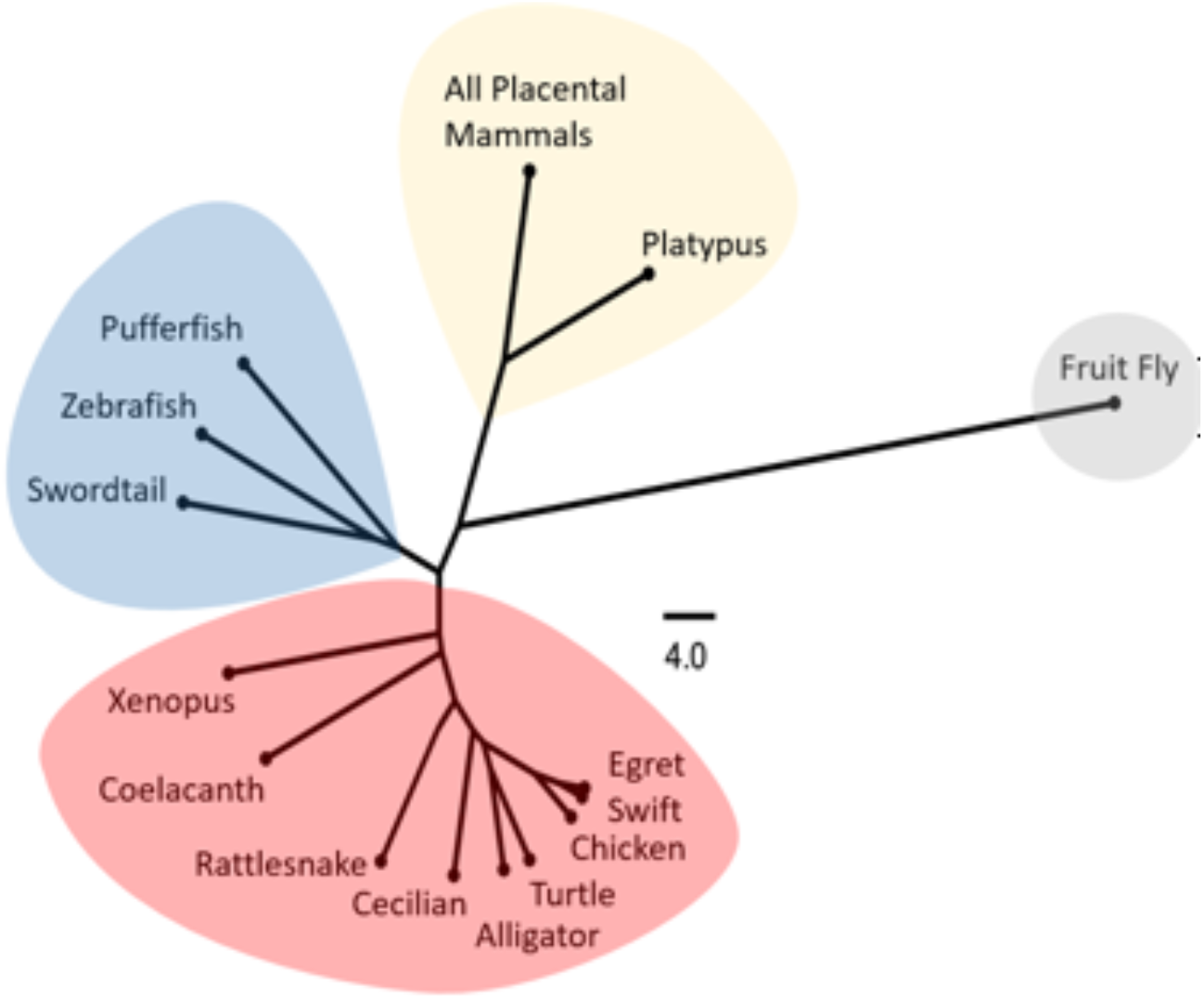
Schematic phylogenetic tree of Prestin TM6 helix sequences, showing the nodes where ancestral Prestin genes have diverged.

**Fig. S22.**
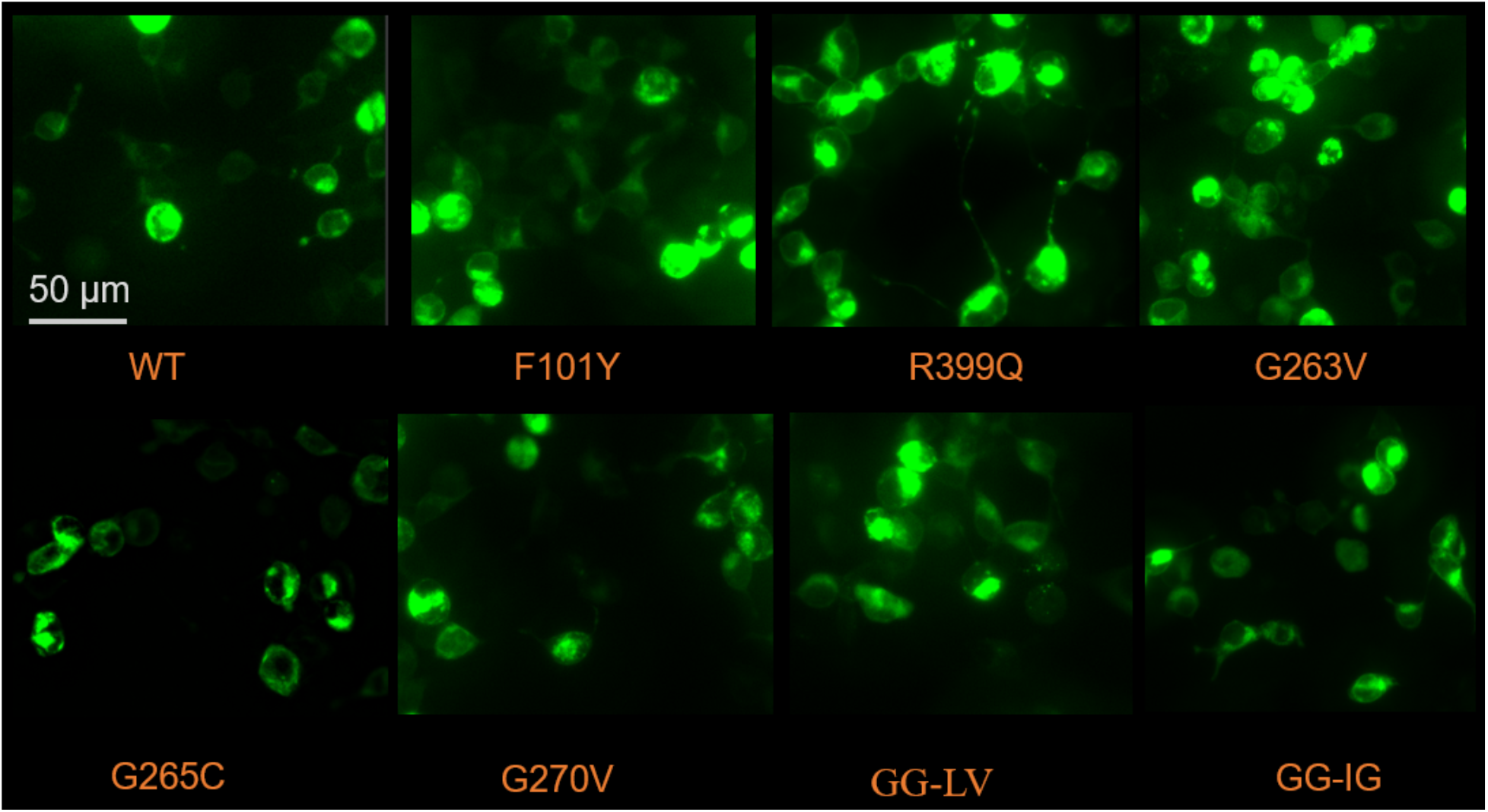
Transient expression and membrane localization of wild type and mutant dolphin Prestin in HEK 293 cells. The grey solid line shows a scale bar of 50 µm. Prestin was over expressed in HEK 293 cells using the Prestin-C-EGFP pEG BacMam construct (see methods). Cellular localization of Prestin was detected by epifluorescence of prestin-EGFP (green) and images were captured by THUNDER microscopy (40X lens; Leica Microsystems). The cells were imaged between 32-36 hours post transfection (repeated 3 times). Both wild type and mutant Prestin are detected in the plasma membrane including in the phillopodia regions (which is most likely devoid of other cellular organelles). No visible difference was detected in terms of plasma membrane trafficking between the mutants and the wild type Prestin. Except that mutation G265C showed a relatively slower rate of expression (expresses post 30 h) compared to wild Prestin and other mutant (expresses post 24 h). Nevertheless, after this period, they are indistinguishable. Each transfection and imaging has been repeated at least three times.

**Fig. S23.**
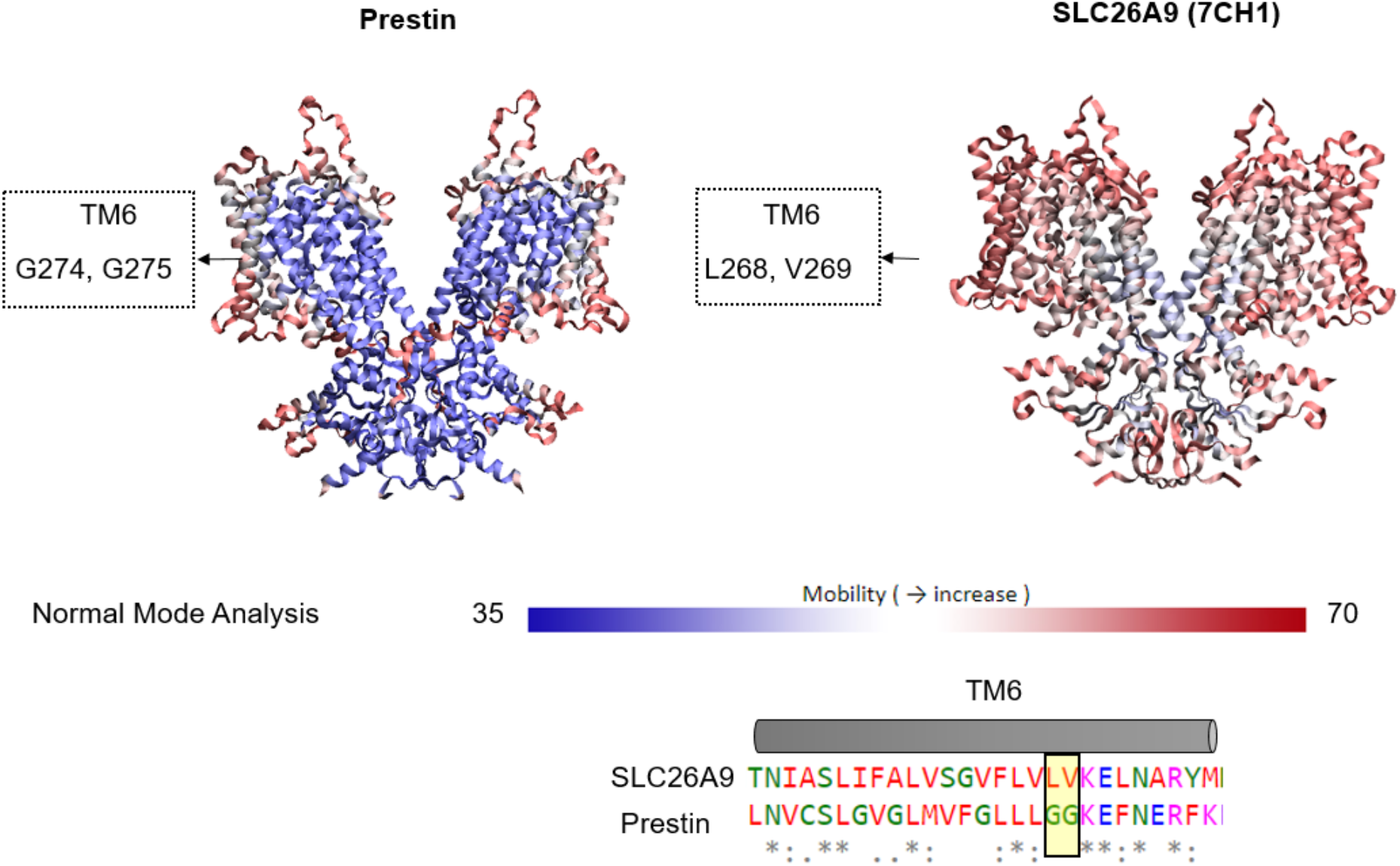
Mobility and flexibility of Prestin versus SLC26A9. Degree of mobility of dolphin Prestin and human SLC26A9 (7CH1), mapped onto their corresponding structure using the DynOmics ENM server and normal mode analysis (*52*). This map shows that while TM6-TM7 region in Prestin has distinct higher mobility from the rest of Prestin structure, the corresponding helices in SLC26A9 have the same level of mobility as several other TM helices in the core and gate domain (indicative of a rigid body motion of the whole region).

**Fig. S24.**
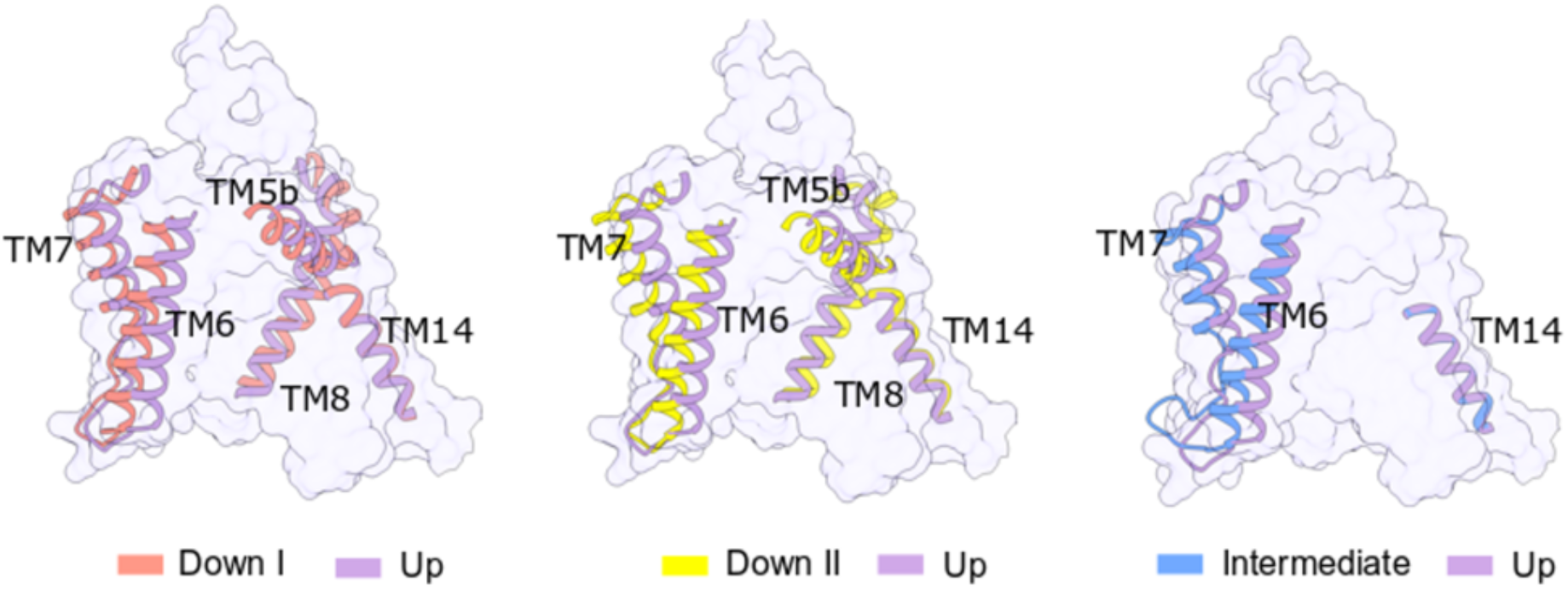
Upon the transition from Down I to Up state and the movement of the anion-binding site, the most obvious changes are seen in the Periphery helices TM5b, TM6-TM7, and TM8.

**Movie S1**

The electromotility measurements of HEK 293 cells transfected with dolphin Prestin using whole-cell patch clamp electrophysiology. To evoke Prestin-mediated electromotility, The membrane potential was held at −70 mV; a 10-mV increase-in-amplitude voltage steps were applied up to the final steps which was from +150 mV to −140 mV (Fig. 1B). The pink square indicates the area that was chosen in our custom-written code to track the cellular displacements.

**Movie S2**

Movement of the anion-binding site (i.e., the voltage sensor) from Down I to Down II conformation. For clarity, only one monomer has been shown. The anion-binding site has been highlighted in red and R399 has been shown in Stick representation and in yellow. The movies are made in UCSF ChimeraX.

**Movie S3**

Structural changes from Down I to, Down II, Intermediate and Up conformation in series. For clarity, only one monomer has been shown. The anion-binding site has been highlighted in red and R399 has been shown in Stick representation and in yellow. The movies are made in UCSF ChimeraX.

**Movie S4**

Structural changes from Down I to, Down II, Intermediate and Up conformation in series. The side front and top views of the dimer have been shown in one single frame. The anion-binding site has been highlighted in red and R399 has been shown in Stick representation and in yellow. The movies are made in UCSF ChimeraX.

## References

1. B. Masterton, H. Heffner, R. Ravizza, The evolution of human hearing. The Journal of the Acoustical Society of America 45, 966–985 (1969).

2. H. Heffner, B. Masterton, Hearing in glires: domestic rabbit, cotton rat, feral house mouse, and kangaroo rat. The Journal of the Acoustical Society of America 68, 1584–1599 (1980).

3. R. Fettiplace, Diverse mechanisms of sound frequency discrimination in the vertebrate cochlea. Trends in neurosciences 43, 88–102 (2020).

4. G. Neuweiler, Auditory adaptations for prey capture in echolocating bats. Physiological reviews 70, 615–641 (1990).

5. Y. Li, Z. Liu, P. Shi, J. Zhang, The hearing gene Prestin unites echolocating bats and whales. Current Biology 20, R55–R56 (2010).

6. K. W. Schultz, D. H. Cato, P. J. Corkeron, M. Bryden, Low frequency narrow-band sounds produced by bottlenose dolphins. Marine Mammal Science 11, 503–509 (1995).

7. W. W. Au, K. J. Snyder, Long-range target detection in open waters by an echolocating Atlantic Bottlenose dolphin (T ursiopstruncatus). The Journal of the Acoustical Society of America 68, 1077–1084 (1980).

8. J. Ashmore, A fast motile response in guinea-pig outer hair cells: the cellular basis of the cochlear amplifier. The Journal of physiology 388, 323–347 (1987).

9. W. E. Brownell, C. R. Bader, D. Bertrand, Y. De Ribaupierre, Evoked mechanical responses of isolated cochlear outer hair cells. Science 227, 194–196 (1985).

10. B. Kachar, W. E. Brownell, R. Altschuler, J. Fex, Electrokinetic shape changes of cochlear outer hair cells. Nature 322, 365–368 (1986).

11. D. Z. He et al., Changes in plasma membrane structure and electromotile properties in prestin deficient outer hair cells. Cytoskeleton 67, 43–55 (2010).

12. J. Zheng et al., Prestin is the motor protein of cochlear outer hair cells. Nature 405, 149–155 (2000).

13. M. Beurg, X. Tan, R. Fettiplace, A prestin motor in chicken auditory hair cells: active force generation in a nonmammalian species. Neuron 79, 69–81 (2013).

14. M. C. Liberman et al., Prestin is required for electromotility of the outer hair cell and for the cochlear amplifier. Nature 419, 300–304 (2002).

15. X. Z. Liu et al., Prestin, a cochlear motor protein, is defective in non-syndromic hearing loss. Human molecular genetics 12, 1155–1162 (2003).

16. P. Dallos et al., Prestin-based outer hair cell motility is necessary for mammalian cochlear amplification. Neuron 58, 333–339 (2008).

17. D. Gorbunov et al., Molecular architecture and the structural basis for anion interaction in prestin and SLC26 transporters. Nature communications 5, 1–13 (2014).

18. R. Hallworth, M. G. Nichols, Prestin in HEK cells is an obligate tetramer. Journal of neurophysiology 107, 5–11 (2012).

19. J. Zheng et al., Analysis of the oligomeric structure of the motor protein prestin. Journal of Biological Chemistry 281, 19916–19924 (2006).

20. D. Navaratnam, J.-P. Bai, H. Samaranayake, J. Santos-Sacchi, N-terminal-mediated homomultimerization of prestin, the outer hair cell motor protein. Biophysical journal 89, 3345–3352 (2005).

21. D. Z. He, S. Lovas, Y. Ai, Y. Li, K. W. Beisel, Prestin at year 14: progress and prospect. Hearing research 311, 25–35 (2014).

22. M. Holley, J. F. Ashmore, On the mechanism of a high-frequency force generator in outer hair cells isolated from the guinea pig cochlea. Proceedings of the Royal society of London. Series B. Biological sciences 232, 413–429 (1988).

23. H. P. Zenner, U. Zimmermann, A. H. Gitter, Fast motility of isolated mammalian auditory sensory cells. Biochemical and biophysical research communications 149, 304–308 (1987).

24. K. Homma, P. Dallos, Evidence that prestin has at least two voltage-dependent steps. Journal of Biological Chemistry 286, 2297–2307 (2011).

25. K. Iwasa, A two-state piezoelectric model for outer hair cell motility. Biophysical journal 81, 2495–2506 (2001).

26. J. Ludwig et al., Reciprocal electromechanical properties of rat prestin: the motor molecule from rat outer hair cells. Proceedings of the National Academy of Sciences 98, 4178–4183 (2001).

27. J. Santos-Sacchi, W. Shen, J. Zheng, P. Dallos, Effects of membrane potential and tension on prestin, the outer hair cell lateral membrane motor protein. The Journal of physiology 531, 661–666 (2001).

28. V. Rybalchenko, J. Santos-Sacchi, Anion control of voltage sensing by the motor protein prestin in outer hair cells. Biophysical journal 95, 4439–4447 (2008).

29. D. Oliver et al., Intracellular anions as the voltage sensor of prestin, the outer hair cell motor protein. Science 292, 2340–2343 (2001).

30. X.-X. Dong, K. Iwasa, Tension sensitivity of prestin: comparison with the membrane motor in outer hair cells. Biophysical journal 86, 1201–1208 (2004).

31. S. Kakehata, J. Santos-Sacchi, Membrane tension directly shifts voltage dependence of outer hair cell motility and associated gating charge. Biophysical journal 68, 2190–2197 (1995).

32. J. Santos-Sacchi, Reversible inhibition of voltage-dependent outer hair cell motility and capacitance. Journal of Neuroscience 11, 3096–3110 (1991).

33. F. Bezanilla, How membrane proteins sense voltage. Nature reviews Molecular cell biology 9, 323–332 (2008).

34. X. Chi et al., Structural insights into the gating mechanism of human SLC26A9 mediated by its C-terminal sequence. Cell discovery 6, 1–10 (2020).

35. J. D. Walter, M. Sawicka, R. Dutzler, Cryo-EM structures and functional characterization of murine Slc26a9 reveal mechanism of uncoupled chloride transport. eLife 8, e46986 (2019).

36. Z. Liu, F.-Y. Qi, D.-M. Xu, X. Zhou, P. Shi, Genomic and functional evidence reveals molecular insights into the origin of echolocation in whales. Science advances 4, eaat8821 (2018).

37. X. Tan et al., From zebrafish to mammal: functional evolution of prestin, the motor protein of cochlear outer hair cells. Journal of neurophysiology 105, 36–44 (2011).

38. J. Santos-Sacchi, L. Song, J. Zheng, A. L. Nuttall, Control of mammalian cochlear amplification by chloride anions. Journal of Neuroscience 26, 3992–3998 (2006).

39. V. Rybalchenko, J. Santos-Sacchi, Cl− flux through a non-selective, stretch-sensitive conductance influences the outer hair cell motor of the guinea-pig. The Journal of physiology 547, 873–891 (2003).

40. K. Homma, C. Duan, J. Zheng, M. A. Cheatham, P. Dallos, The V499G/Y501H mutation impairs fast motor kinetics of prestin and has significance for defining functional independence of individual prestin subunits. Journal of Biological Chemistry 288, 2452–2463 (2013).

41. L. Song, J. Santos-Sacchi, Conformational state-dependent anion binding in prestin: evidence for allosteric modulation. Biophysical journal 98, 371–376 (2010).

42. V. Jogini, B. Roux, Dynamics of the Kv1. 2 voltage-gated K+ channel in a membrane environment. Biophysical journal 93, 3070–3082 (2007).

43. D. M. Starace, F. Bezanilla, A proton pore in a potassium channel voltage sensor reveals a focused electric field. Nature 427, 548–553 (2004).

44. X.-x. Dong, D. Ehrenstein, K. Iwasa, Fluctuation of motor charge in the lateral membrane of the cochlear outer hair cell. Biophysical journal 79, 1876–1882 (2000).

45. J.-P. Bai et al., Current carried by the Slc26 family member prestin does not flow through the transporter pathway. Scientific reports 7, 1–13 (2017).

46. J.-P. Bai et al., Prestin’s anion transport and voltage-sensing capabilities are independent. Biophysical journal 96, 3179–3186 (2009).

47. J. Ashmore, Cochlear outer hair cell motility. Physiological reviews 88, 173–210 (2008).

48. D. Z. He, B. N. Evans, P. Dallos, First appearance and development of electromotility in neonatal gerbil outer hair cells. Hearing research 78, 77–90 (1994).

49. P. Dallos, The active cochlea. Journal of Neuroscience 12, 4575–4585 (1992).

50. J. Santos-Sacchi, Asymmetry in voltage-dependent movements of isolated outer hair cells from the organ of Corti. Journal of Neuroscience 9, 2954–2962 (1989).

51. J. Santos-Sacchi, J. Dilger, Whole cell currents and mechanical responses of isolated outer hair cells. Hearing research 35, 143–150 (1988).

52. Y. Cazals, Auditory sensori-neural alterations induced by salicylate. Progress in neurobiology 62, 583–631 (2000).

53. G.-D. Chen et al., Salicylate-induced cochlear impairments, cortical hyperactivity and re-tuning, and tinnitus. Hearing research 295, 100–113 (2013).

54. S. Kakehata, J. Santos-Sacchi, Effects of salicylate and lanthanides on outer hair cell motility and associated gating charge. Journal of Neuroscience 16, 4881–4889 (1996).

55. T. J. Schaechinger, D. Oliver, Nonmammalian orthologs of prestin (SLC26A5) are electrogenic divalent/chloride anion exchangers. Proceedings of the National Academy of Sciences 104, 7693–7698 (2007).

56. C. Izumi, J. E. Bird, K. H. Iwasa, Membrane thickness sensitivity of prestin orthologs: the evolution of a piezoelectric protein. Biophysical journal 100, 2614–2622 (2011).

57. J. Fang, C. Izumi, K. H. Iwasa, Sensitivity of prestin-based membrane motor to membrane thickness. Biophysical journal 98, 2831–2838 (2010).

58. N. Bavi, C. D. Cox, E. Perozo, B. Martinac, Toward a structural blueprint for bilayer-mediated channel mechanosensitivity. Channels (Austin) 11, 91–93 (2017).

59. J. Santos-Sacchi, On the frequency limit and phase of outer hair cell motility: effects of the membrane filter. Journal of Neuroscience 12, 1906–1916 (1992).

60. J. Santos-Sacchi, E. Navarrete, L. Song, Fast electromechanical amplification in the lateral membrane of the outer hair cell. Biophysical journal 96, 739–747 (2009).

## References

1. Z. Liu, F.-Y. Qi, D.-M. Xu, X. Zhou, P. Shi, Genomic and functional evidence reveals molecular insights into the origin of echolocation in whales. Science advances 4, eaat8821 (2018).

2. A. Kirchhofer et al., Modulation of protein properties in living cells using nanobodies. Nature structural & molecular biology 17, 133–138 (2010).

3. M. D. Clark, G. F. Contreras, R. Shen, E. Perozo, Electromechanical coupling in the hyperpolarization-activated K+ channel KAT1. Nature 583, 145–149 (2020).

4. S. H. Scheres, RELION: implementation of a Bayesian approach to cryo-EM structure determination. Journal of structural biology 180, 519–530 (2012).

5. S. Q. Zheng et al., MotionCor2: anisotropic correction of beam-induced motion for improved cryo-electron microscopy. Nature methods 14, 331–332 (2017).

6. A. Rohou, N. Grigorieff, CTFFIND4: Fast and accurate defocus estimation from electron micrographs. Journal of structural biology 192, 216–221 (2015).

7. T. Wagner et al., SPHIRE-crYOLO is a fast and accurate fully automated particle picker for cryo-EM. Communications biology 2, 1–13 (2019).

8. S. H. Scheres, S. Chen, Prevention of overfitting in cryo-EM structure determination. Nature methods 9, 853–854 (2012).

9. P. B. Rosenthal, R. Henderson, Optimal determination of particle orientation, absolute hand, and contrast loss in single-particle electron cryomicroscopy. Journal of molecular biology 333, 721–745 (2003).

10. A. Kucukelbir, F. J. Sigworth, H. D. Tagare, Quantifying the local resolution of cryo-EM density maps. Nature methods 11, 63–65 (2014).

11. M. Biasini et al., SWISS-MODEL: modelling protein tertiary and quaternary structure using evolutionary information. Nucleic acids research 42, W252–W258 (2014).

12. J. D. Walter, M. Sawicka, R. Dutzler, Cryo-EM structures and functional characterization of murine Slc26a9 reveal mechanism of uncoupled chloride transport. eLife 8, e46986 (2019).

13. X. Chi et al., Structural insights into the gating mechanism of human SLC26A9 mediated by its C-terminal sequence. Cell discovery 6, 1–10 (2020).

14. N. Stein, CHAINSAW: a program for mutating pdb files used as templates in molecular replacement. Journal of applied crystallography 41, 641–643 (2008).

15. P. Emsley, K. Cowtan, Coot: model-building tools for molecular graphics. Acta crystallographica section D: biological crystallography 60, 2126–2132 (2004).

16. P. Emsley, B. Lohkamp, W. G. Scott, K. Cowtan, Features and development of Coot. Acta Crystallographica Section D: Biological Crystallography 66, 486–501 (2010).

17. A. Brown et al., Tools for macromolecular model building and refinement into electron cryo-microscopy reconstructions. Acta Crystallographica Section D: Biological Crystallography 71, 136–153 (2015).

18. P. D. Adams et al., PHENIX: a comprehensive Python-based system for macromolecular structure solution. Acta Crystallographica Section D: Biological Crystallography 66, 213–221 (2010).

19. P. V. Afonine et al., Real-space refinement in PHENIX for cryo-EM and crystallography. Acta Crystallographica Section D: Structural Biology 74, 531–544 (2018).

20. E. F. Pettersen et al., UCSF Chimera—a visualization system for exploratory research and analysis. Journal of computational chemistry 25, 1605–1612 (2004).

21. E. F. Pettersen et al., UCSF ChimeraX: Structure visualization for researchers, educators, and developers. Protein Science 30, 70–82 (2021).

22. T. D. Goddard et al., UCSF ChimeraX: Meeting modern challenges in visualization and analysis. Protein Science 27, 14–25 (2018).

23. W. Humphrey, A. Dalke, K. Schulten, VMD: visual molecular dynamics. Journal of molecular graphics 14, 33–38 (1996).

24. J. Ashmore, A fast motile response in guinea-pig outer hair cells: the cellular basis of the cochlear amplifier. The Journal of physiology 388, 323–347 (1987).

25. J. Santos-Sacchi, Reversible inhibition of voltage-dependent outer hair cell motility and capacitance. Journal of Neuroscience 11, 3096–3110 (1991).

26. V. Rybalchenko, J. Santos-Sacchi, Anion control of voltage sensing by the motor protein prestin in outer hair cells. Biophysical journal 95, 4439–4447 (2008).

27. C. Camacho et al., BLAST+: architecture and applications. BMC bioinformatics 10, 1–9 (2009).

28. R. C. Edgar, MUSCLE: multiple sequence alignment with high accuracy and high throughput. Nucleic acids research 32, 1792–1797 (2004).

29. A. Stamatakis, RAxML version 8: a tool for phylogenetic analysis and post-analysis of large phylogenies. Bioinformatics 30, 1312–1313 (2014).

30. I. Letunic, P. Bork, Interactive Tree Of Life (iTOL) v4: recent updates and new developments. Nucleic acids research 47, W256–W259 (2019).

31. M. A. Lomize, I. D. Pogozheva, H. Joo, H. I. Mosberg, A. L. Lomize, OPM database and PPM web server: resources for positioning of proteins in membranes. Nucleic acids research 40, D370–D376 (2012).

32. A. Morozenko, A. Stuchebrukhov, Dowser++, a new method of hydrating protein structures. Proteins: Structure, Function, and Bioinformatics 84, 1347–1357 (2016).

33. D. E. Shaw et al., Anton, a special-purpose machine for molecular dynamics simulation. Communications of the ACM 51, 91–97 (2008).

34. J. C. Phillips et al., Scalable molecular dynamics with NAMD. Journal of computational chemistry 26, 1781–1802 (2005).

35. A. D. MacKerell Jr, M. Feig, C. L. Brooks, Improved treatment of the protein backbone in empirical force fields. Journal of the American Chemical Society 126, 698–699 (2004).

36. A. D. MacKerell Jr et al., All-atom empirical potential for molecular modeling and dynamics studies of proteins. The journal of physical chemistry B 102, 3586–3616 (1998).

37. W. L. Jorgensen, J. Chandrasekhar, J. D. Madura, R. W. Impey, M. L. Klein, Comparison of simple potential functions for simulating liquid water. The Journal of chemical physics 79, 926–935 (1983).

38. L. Huang, B. Roux, Automated force field parameterization for nonpolarizable and polarizable atomic models based on ab initio target data. Journal of chemical theory and computation 9, 3543–3556 (2013).

39. G. J. Martyna, D. J. Tobias, M. L. Klein, Constant pressure molecular dynamics algorithms. The Journal of chemical physics 101, 4177–4189 (1994).

40. S. E. Feller, Y. Zhang, R. W. Pastor, B. R. Brooks, Constant pressure molecular dynamics simulation: the Langevin piston method. The Journal of chemical physics 103, 4613–4621 (1995).

41. U. Essmann et al., A smooth particle mesh Ewald method. The Journal of chemical physics 103, 8577–8593 (1995).

42. G. J. Martyna, M. L. Klein, M. Tuckerman, Nosé–Hoover chains: The canonical ensemble via continuous dynamics. The Journal of chemical physics 97, 2635–2643 (1992).

43. Y. Shan, J. L. Klepeis, M. P. Eastwood, R. O. Dror, D. E. Shaw, Gaussian split Ewald: A fast Ewald mesh method for molecular simulation. The Journal of chemical physics 122, 054101 (2005).

44. A. Aksimentiev, K. Schulten, Imaging α-hemolysin with molecular dynamics: ionic conductance, osmotic permeability, and the electrostatic potential map. Biophysical journal 88, 3745–3761 (2005).

45. B. Roux, The membrane potential and its representation by a constant electric field in computer simulations. Biophysical journal 95, 4205–4216 (2008).

46. J. P. Castillo et al., Mechanism of potassium ion uptake by the Na+/K+-ATPase. Nature communications 6, 1–8 (2015).

47. S. Jo, T. Kim, V. G. Iyer, W. Im, CHARMM-GUI: a web-based graphical user interface for CHARMM. Journal of computational chemistry 29, 1859–1865 (2008).

48. F. Khalili-Araghi et al., Calculation of the gating charge for the Kv1. 2 voltage-activated potassium channel. Biophysical journal 98, 2189–2198 (2010).

49. J. Santos-Sacchi, J. Dilger, Whole cell currents and mechanical responses of isolated outer hair cells. Hearing research 35, 143–150 (1988).

50. J. Santos-Sacchi, On the frequency limit and phase of outer hair cell motility: effects of the membrane filter. Journal of Neuroscience 12, 1906–1916 (1992).

51. J. Ashmore, Cochlear outer hair cell motility. Physiological reviews 88, 173–210 (2008).

