## Supplemental Methods and Figures for "Cryo-EM Structures of Prestin and the Molecular Basis of Outer Hair Cell Electromotility"

$$Y_\omega = \frac{FFT(V(t))}{FFT(I(t))} \quad \text{Eq. (1)}$$

$$A = \Re(Y_\omega) \quad \text{Eq. (2)}$$

$$B = \Im(Y_\omega) \quad \text{Eq. (3)}$$

$$Rs = \frac{A-b}{(A^2+B^2-Ab)} \quad \text{Eq. (4)}$$

$$Rm = \frac{(A-b)^2+B^2}{b*(A^2+B^2-Ab)} \quad \text{Eq. (5)}$$

$$Cm = \frac{1}{\omega B} * \frac{\left(A^2+B^2-Ab\right)^2}{(A-b)^2+B^2} \qquad \text{Eq. (6)}$$

The membrane capacitance was fitted to the derivative of a Boltzmann function plus a lineal component:

$$C_m = C_{lin} + \frac{Q_{max}\alpha}{exp\left[\alpha\left(V-V_{\frac{1}{2}}\right)\right]\left(1+exp\left[-\alpha\left(V-V_{\frac{1}{2}}\right)\right]\right)^2} \qquad \text{Eq. (7)}$$

$$\Delta L_{OHC} = A_{OHC} \times \rho \times \Delta A_P / (2\pi\, R_{OHC}) \qquad \text{Eq. (9)}$$

Where:

$\Delta L_{OHC}$, Maximum possible somatic motility of the OHC based solely based on collective cross-sectional expansion of Prestin molecules in the lateral wall.

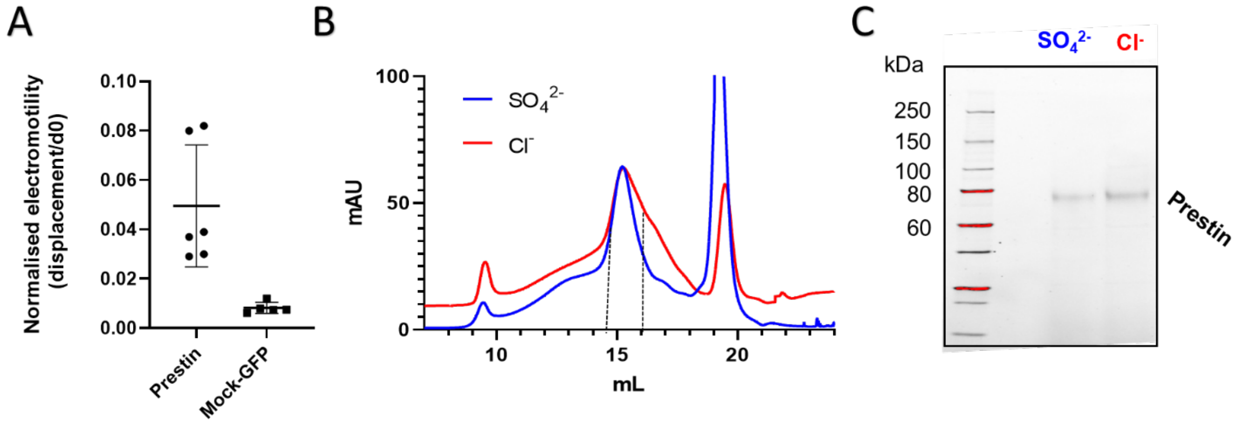


**Fig. S1.** (A) Electromotility analysis of HEK 293 cells transfected with wild type dolphin Prestin compared to GFP-only transfected cell (Mock-GFP). The cellular displacement has been normalized based on the cell largest diameter, $d_0$ (Fig. 1B). The normalized electromotility was 0.05±02 versus 0.008±0.002 for wild type Prestin and Mock-GFP, respectively. These values were measured at the depolarizing voltage step changing from +120 mV to -120 mV (mean ± SD; Nonparametric Student t-test, unpaired, $P$=0.005). (B) Size exclusion chromatography (SEC) curves of the full-length dolphin Prestin purified in GDN, run on a Superose 6 column, in high $Cl^-$ (red) and $SO_4^{2-}$ (blue) based solution. The fractions indicated by black dotted lines in both represent purified proteins that were used for cryo-EM imaging. (B) Purified dolphin Prestin cryo-EM samples, run on a Stain-free SDS-PAGE gel, indicating size of ~ 75 kDa for the full-length Prestin monomer.

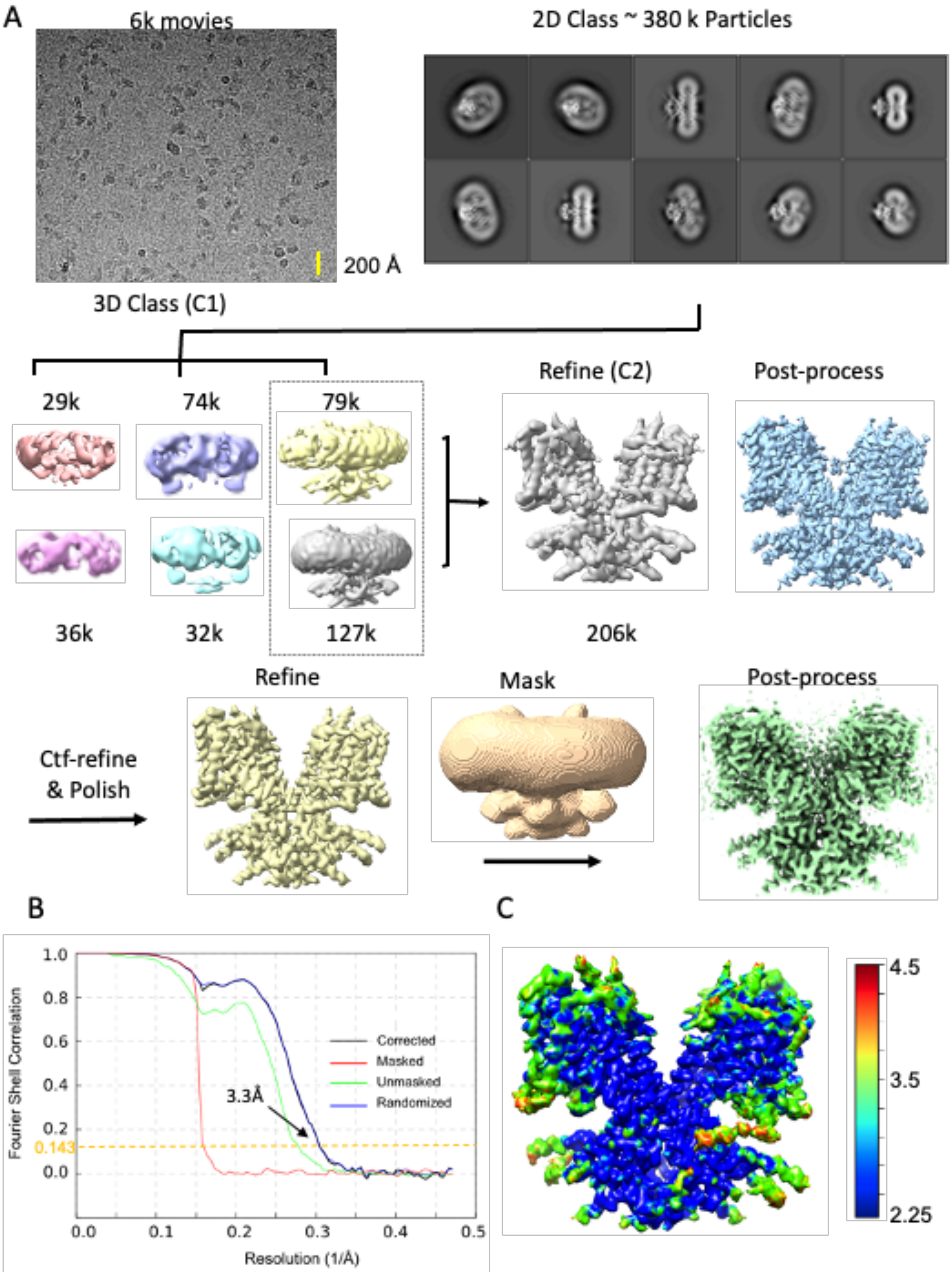


**Fig. S2.** Flow chart for the cryo-EM data processing and structure determination of the dolphin Prestin in high $Cl^-$ condition (See Methods for details). The final reconstruction has a normal resolution of 3.3 Å (at Fsc=0.143). All the images in this figure were created in UCSF ChimeraX.

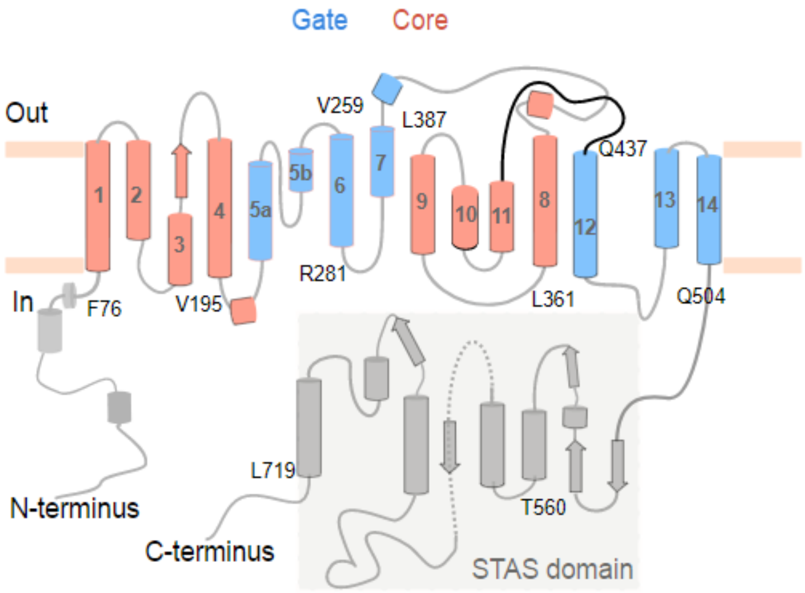


**Fig. S3.** Topology of dolphin Prestin. Different domains are indicated by color; the gate domain is colored in blue, the core domain in red and the C- and N-termini as well as the STAS domain in grey. The transmembrane helices are numbered from 1 to 14. The N- and C-termini as well as the STAS domain are oriented towards the cytoplasm.

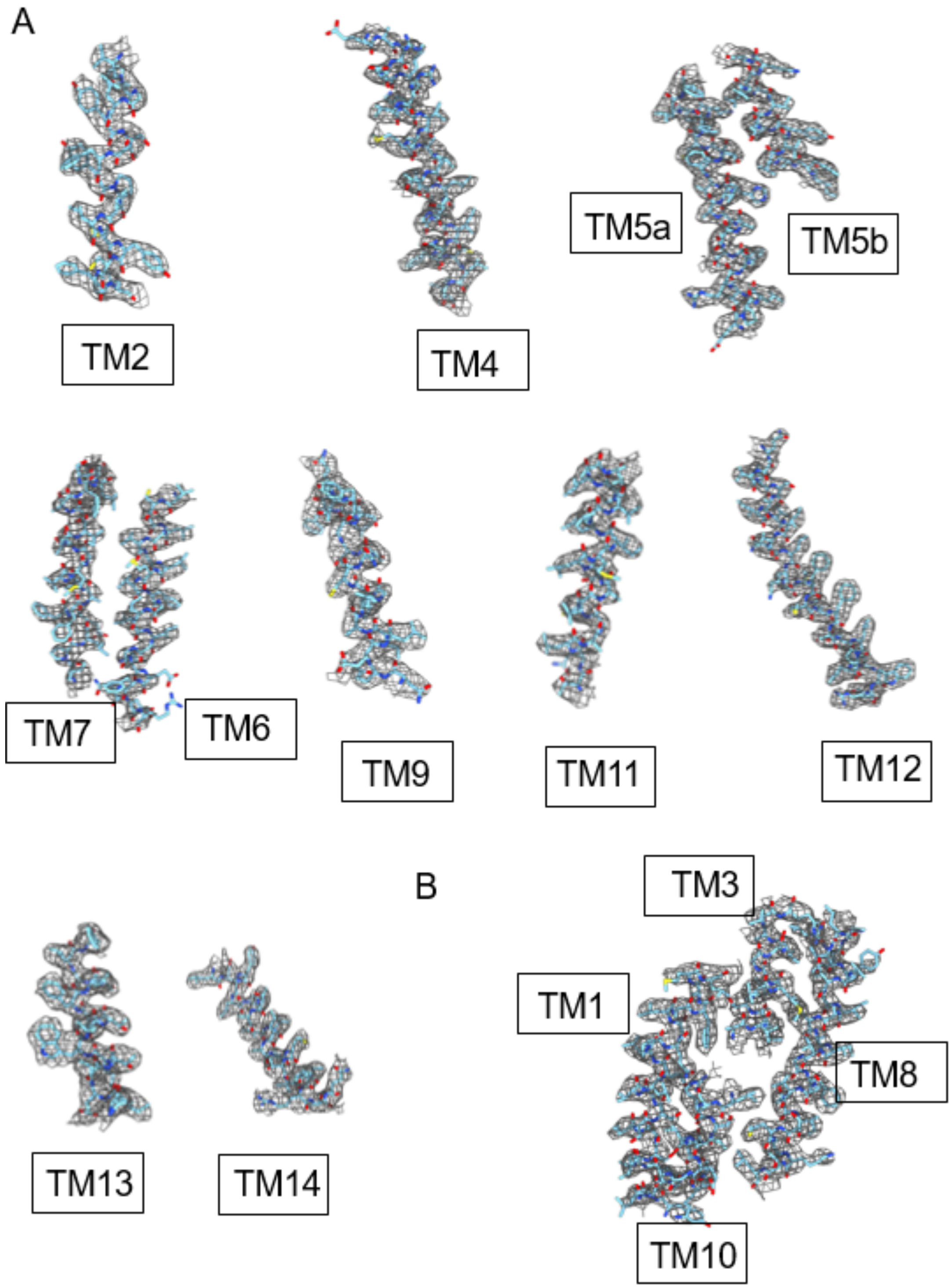


**Fig. S4.** Details of the atomic model fitted to the postprocessed cryo-EM density map are illustrated for (A) the transmembrane helices including those forming (B) the anion-binding pocket.

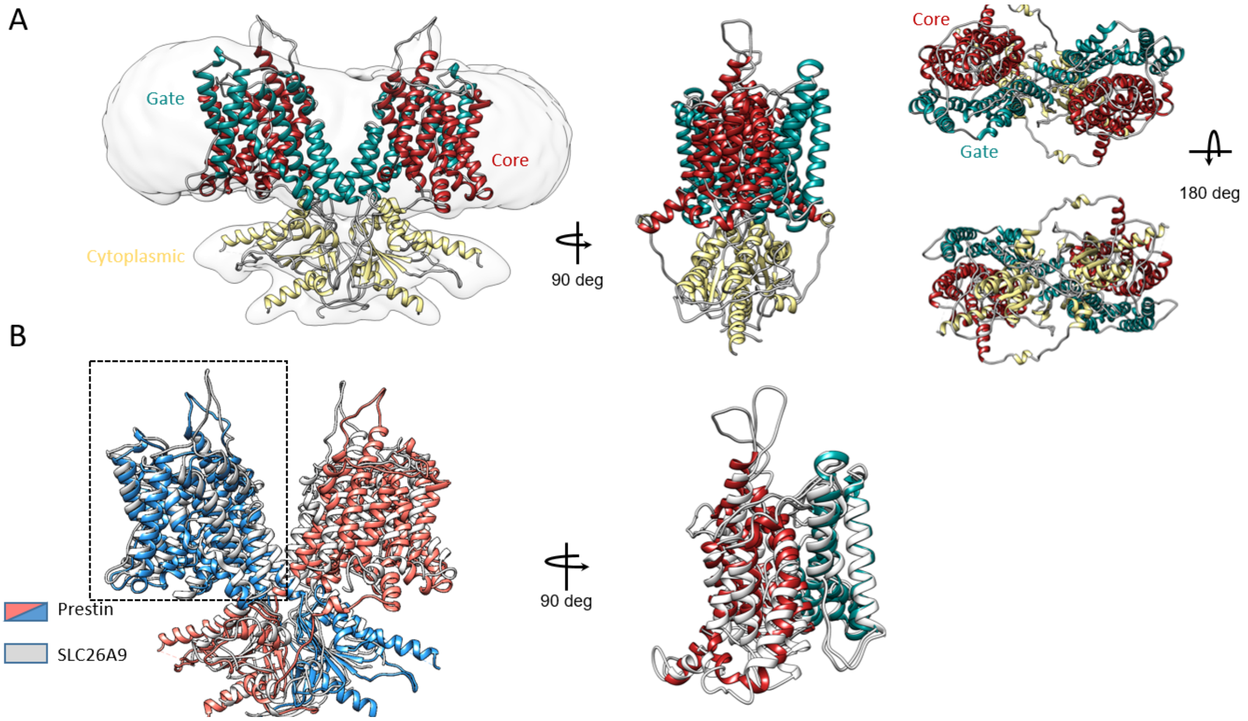


**Fig. S5.** Structure of Prestin in high Cl$^-$ and comparison with the Intermediate conformation of SLC26A9 (PDB:6RTF). (A) Overall architecture of dolphin Prestin at (high Cl$^-$) fit to the cryo-EM density, with different domains highlighted by color. (B) Comparison between Prestin (high Cl$^-$) (blue and Red) and SLC26A9 Intermediate state (grey). Subunits A (dotted box) from both proteins were aligned. The structures are aligned based on residue 460 to 505 (TM13-TM14). ChimeraX was used for illustration.

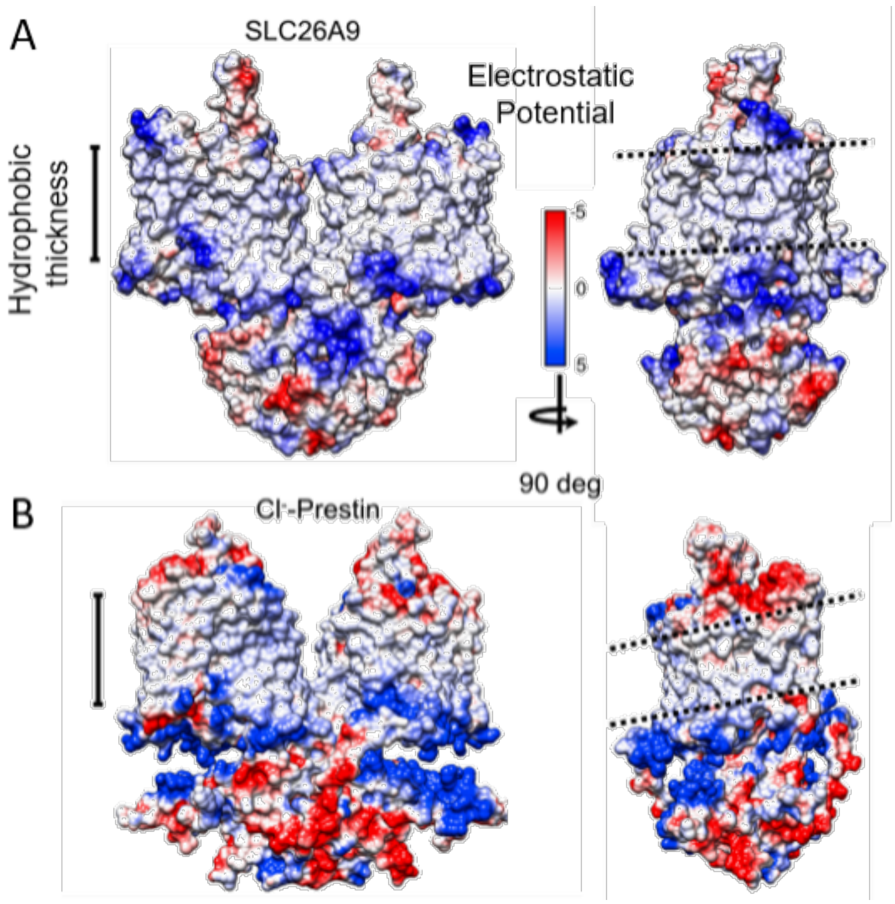


**Fig. S6.** Electrostatic potential and surface charge distribution of SLC26A9 intermediate state (6RTC, panel (A)) compared with that of Prestin in high $Cl^-$ panel (B). The electrostatic charge distribution ranges from -5 to 5 kT from negative to positive charge. ChimeraX was used for illustration.

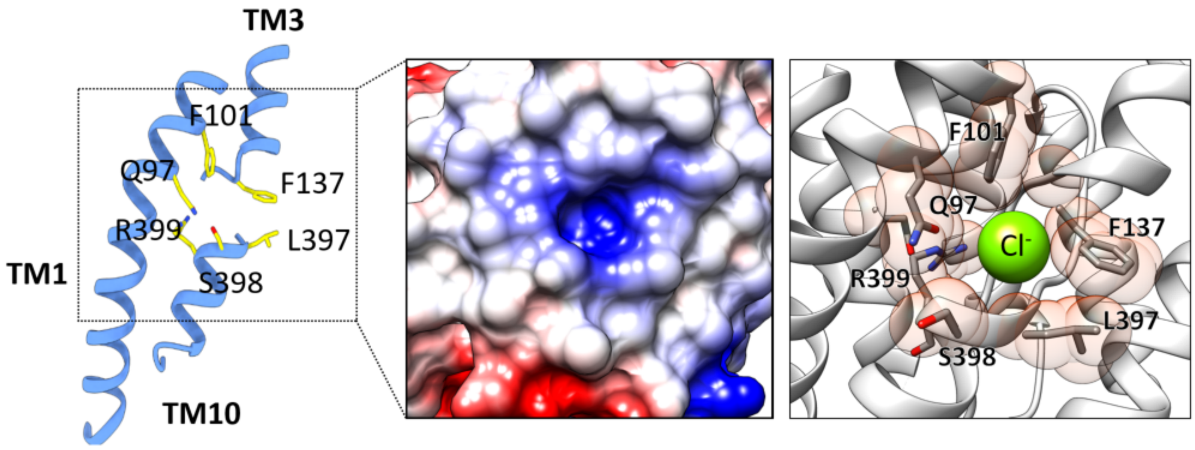


**Fig. S7.** Detailed Structure of Prestin's anion-binding site. Left panel: Key anion binding site residues Q97, F101, F137, L397, S398 and R399 are shown in yellow color and by their heteroatom. Middle panel: the electrostatic potential at this site shows a positive field (middle panel); we manually placed a $Cl^-$ anion into the positive cavity (right panel). Only TM10-TM3 dipole and TM1 are shown for clarity (right panel). UCSF ChimeraX was used for illustration.

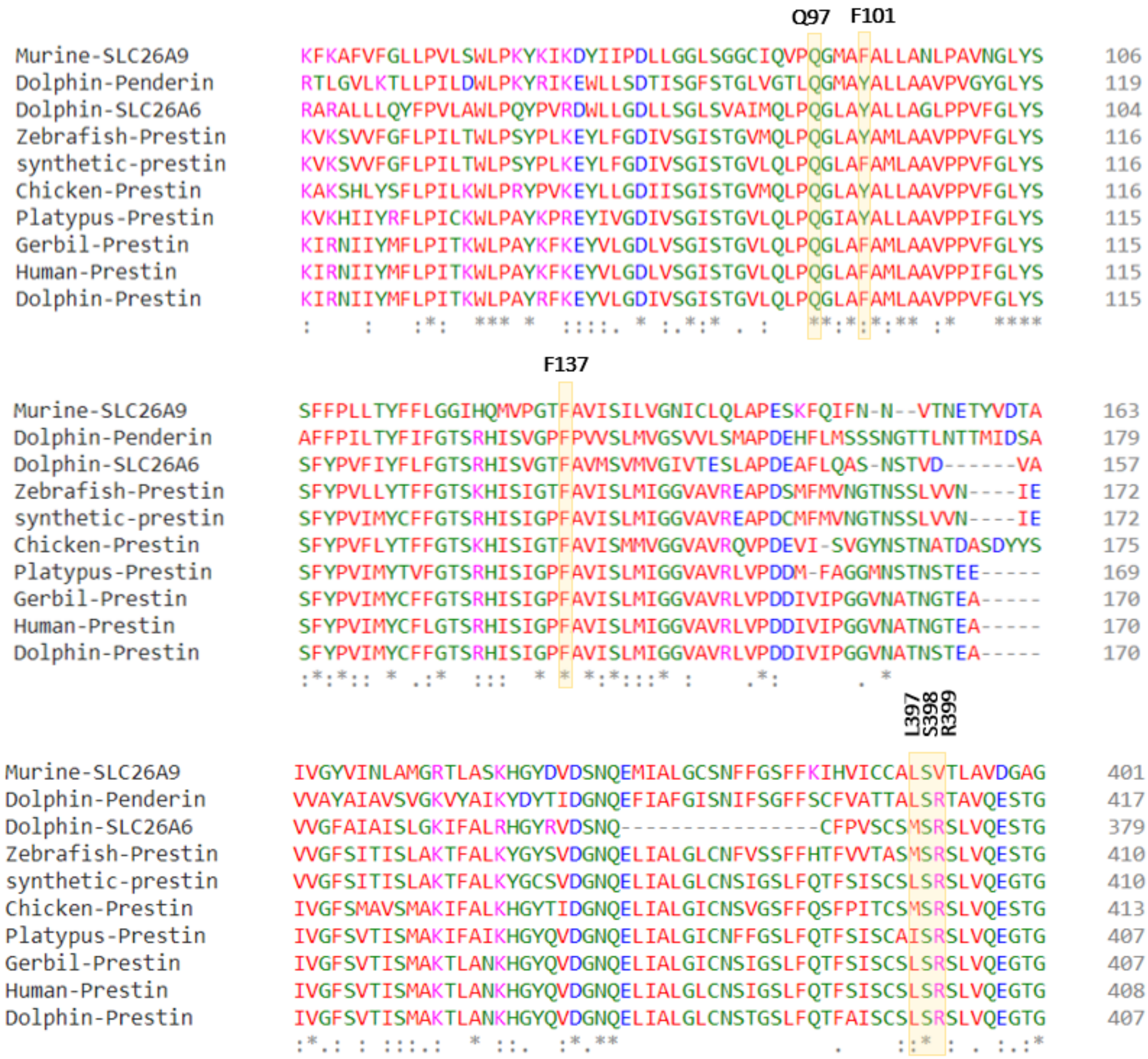


**Fig. S8.** Sequence alignment of Prestin and close SLC transporters across different species. Residues forming the anion-binding site are largely conserved (e.g. Q97, F101, F137). Putative voltage-sensing residue R399 in dolphin Prestin is replaced by a valine in murine SLC26A9. Clustal Omega was used for the sequence alignments.

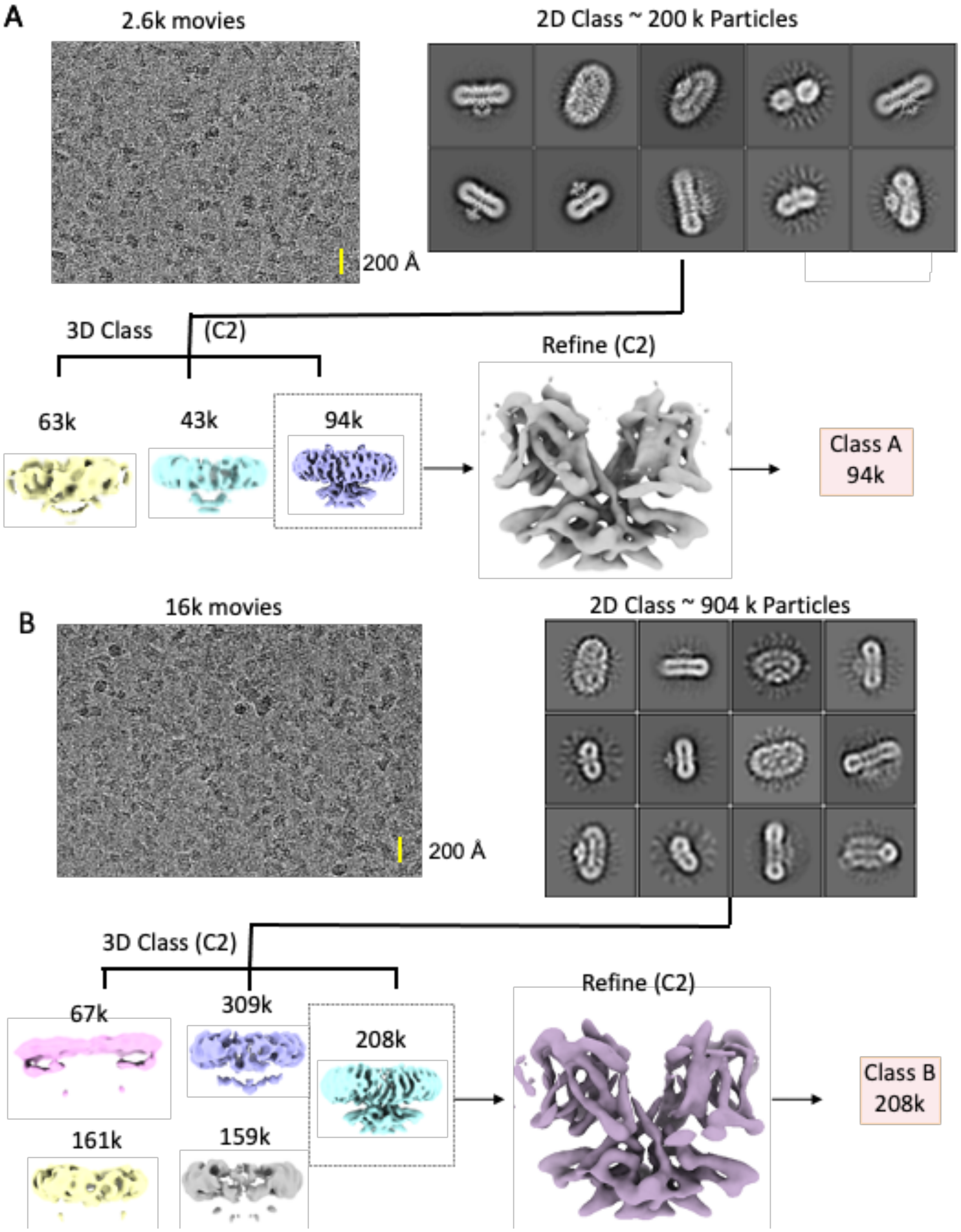


**Fig. S9.** Flow chart for the cryo-EM data processing and structure determination of the dolphin Prestin in $SO_4^{2-}$ (see methods). UCSF ChimeraX was used for illustration.

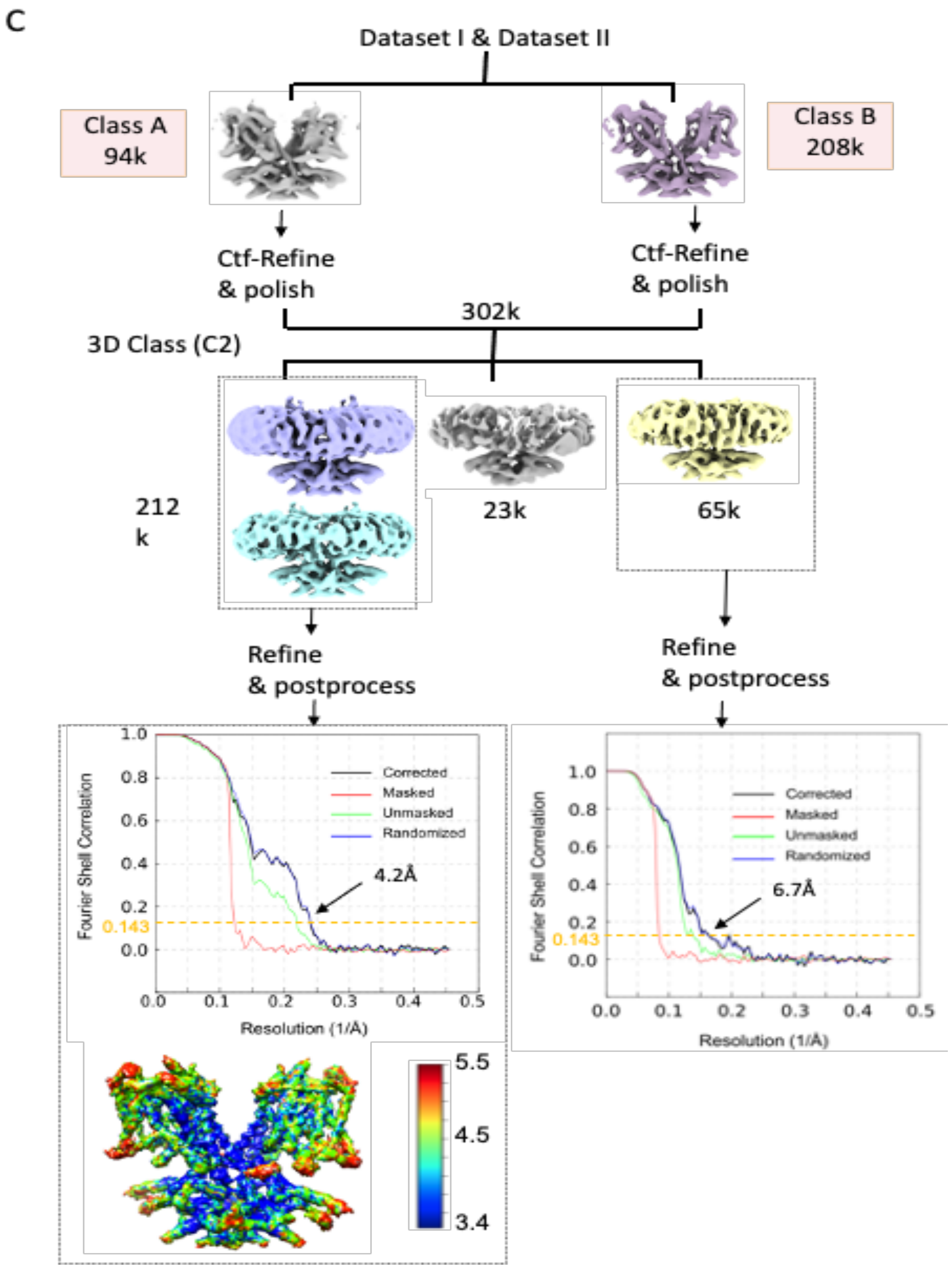


**Fig. S10.** Flow chart for the cryo-EM data processing and structure determination of the dolphin Prestin in Down I ($SO_4^{2-}$) and Down II ($SO_4^{2-}$) states (See Methods for details). Class A was obtained from Dataset I, which was combined with (B) Class B from Dataset II. (C) The final reconstruction yielded two structures, Down I ($SO_4^{2-}$) and Down II ($SO_4^{2-}$), which have nominal resolutions of 4.2 and 6.7 Å, respectively (at FSC=0.143). Evidence of both states was found in dataset II, however merging of datasets was required to improve resolution of states. All the images in this figure were created in UCSF ChimeraX.

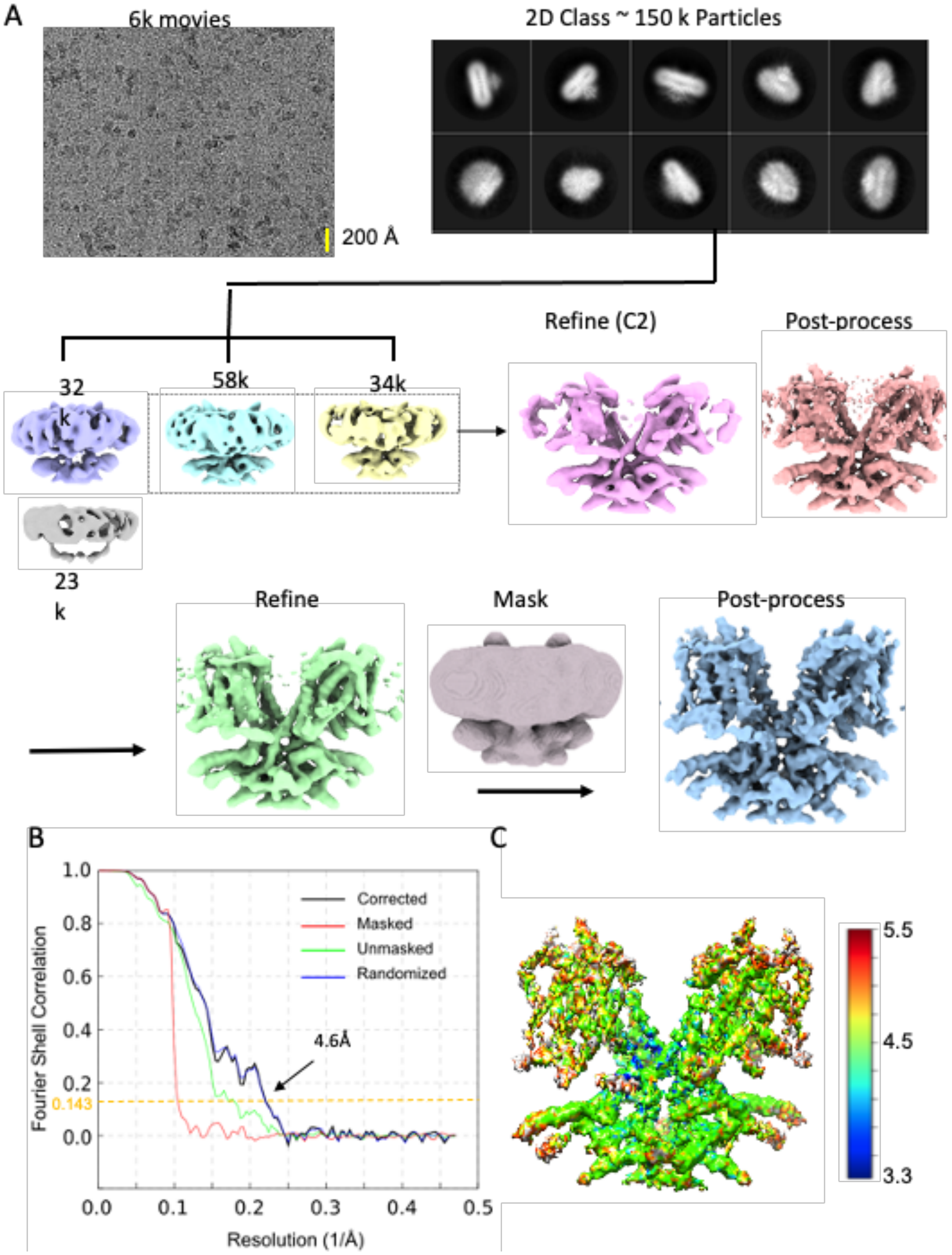


**Fig. S11.** Flow chart for the cryo-EM data processing and structure determination of the dolphin Prestin in the Intermediate state ($SO_4^{2-}$) (See Methods for details). The final reconstruction has a nominal resolution of 4.6 Å (at FSC=0.143). All the structures were illustrated in UCSF ChimeraX.

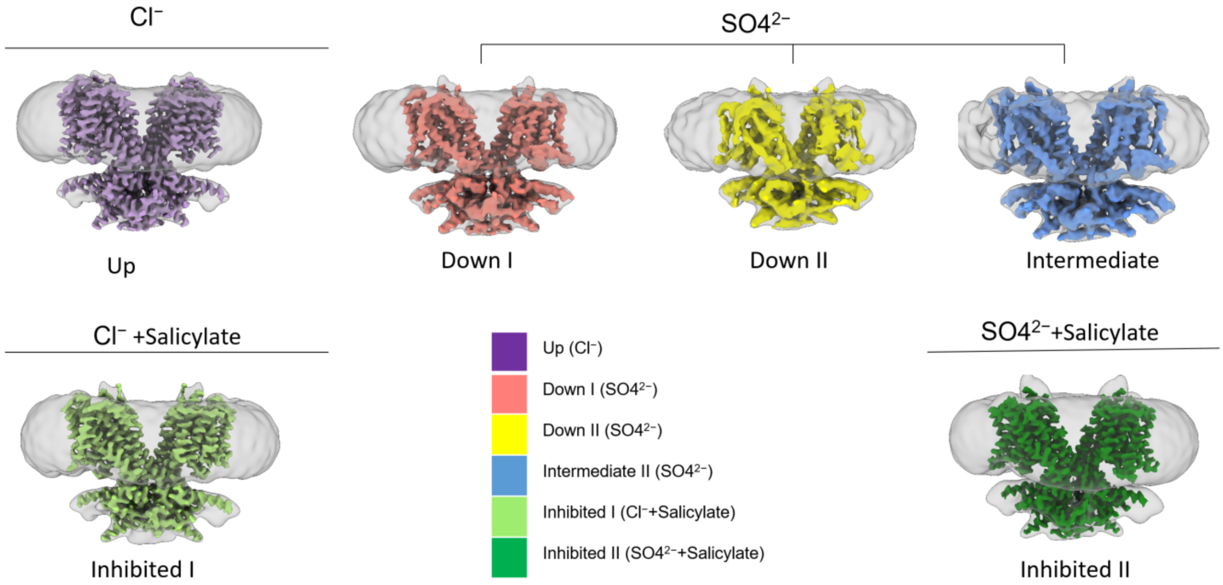


**Fig. S12.** Overview of determined structures**.** Dolphin Prestin's structure has been captured in six distinct states, namely: Up ($Cl^-$), Down I ($SO_4^{2-}$), Down II ($SO_4^{2-}$), Intermediate ($SO_4^{2-}$), Inhibited I ($Cl^-$ + salicylate) and inhibited II ($SO_4^{2-}$+salicylate). All the structures were illustrated in UCSF ChimeraX.

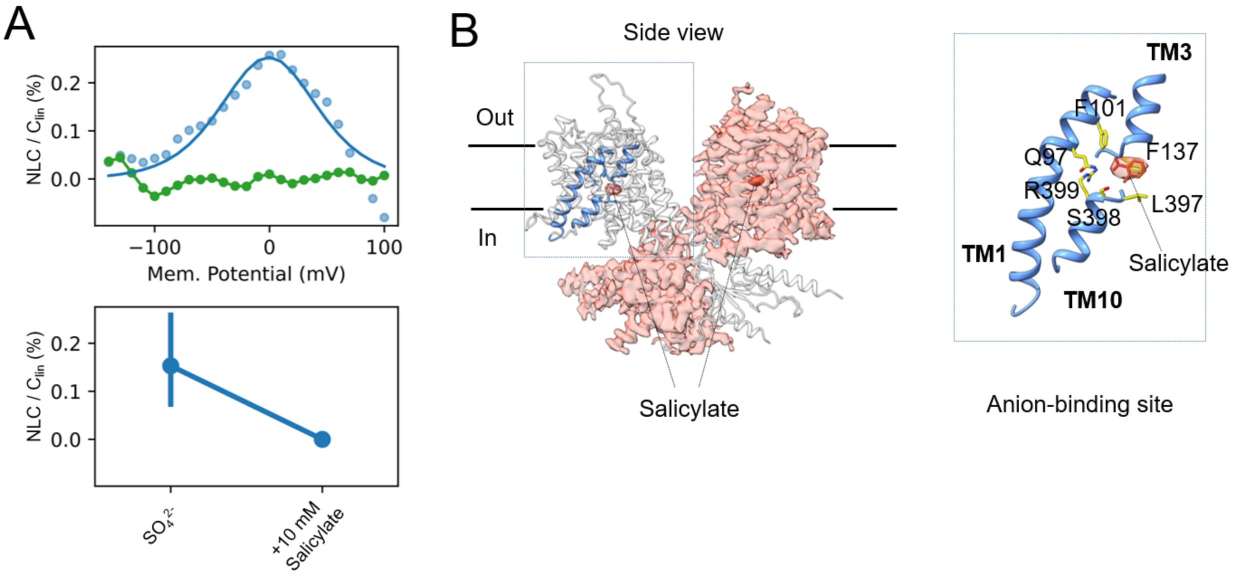


**Fig. S13.** Salicylate outcompetes $SO_4^{2-}$ in binding to anion-binding pocket. (A) Patch clamp electrophysiology of HEK293 cells transfected with Dolphin Prestin. The NLC is abrogated by 10 mM Na-Salicylate, when $SO_4^{2-}$ is the main anion of the bath and pipette solutions (See Methods). (B) Density of Salicylate (orange) in the anion-binding site (blue) was resolved in the Inhibited II ($SO_4^{2-}$) state of dolphin Prestin.

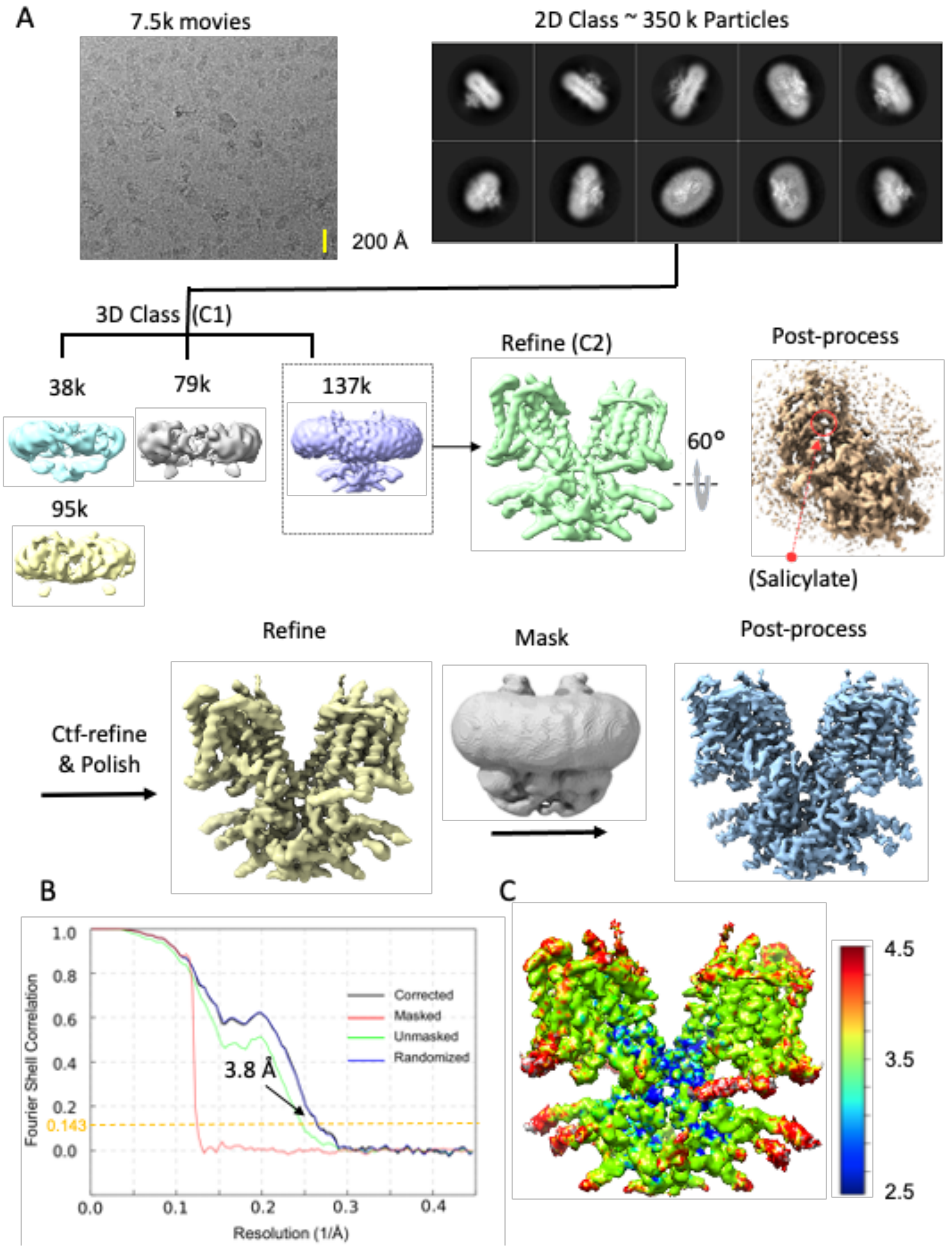


**Fig. S14.** Flow chart for the cryo-EM data processing and structure determination of the dolphin Prestin in the Inhibited I state ($Cl^-$ + Salicylate) (See Methods for details). The final reconstruction has a nominal resolution of 3.8 Å (at FSC=0.143). All the images in this figure were created in UCSF ChimeraX.

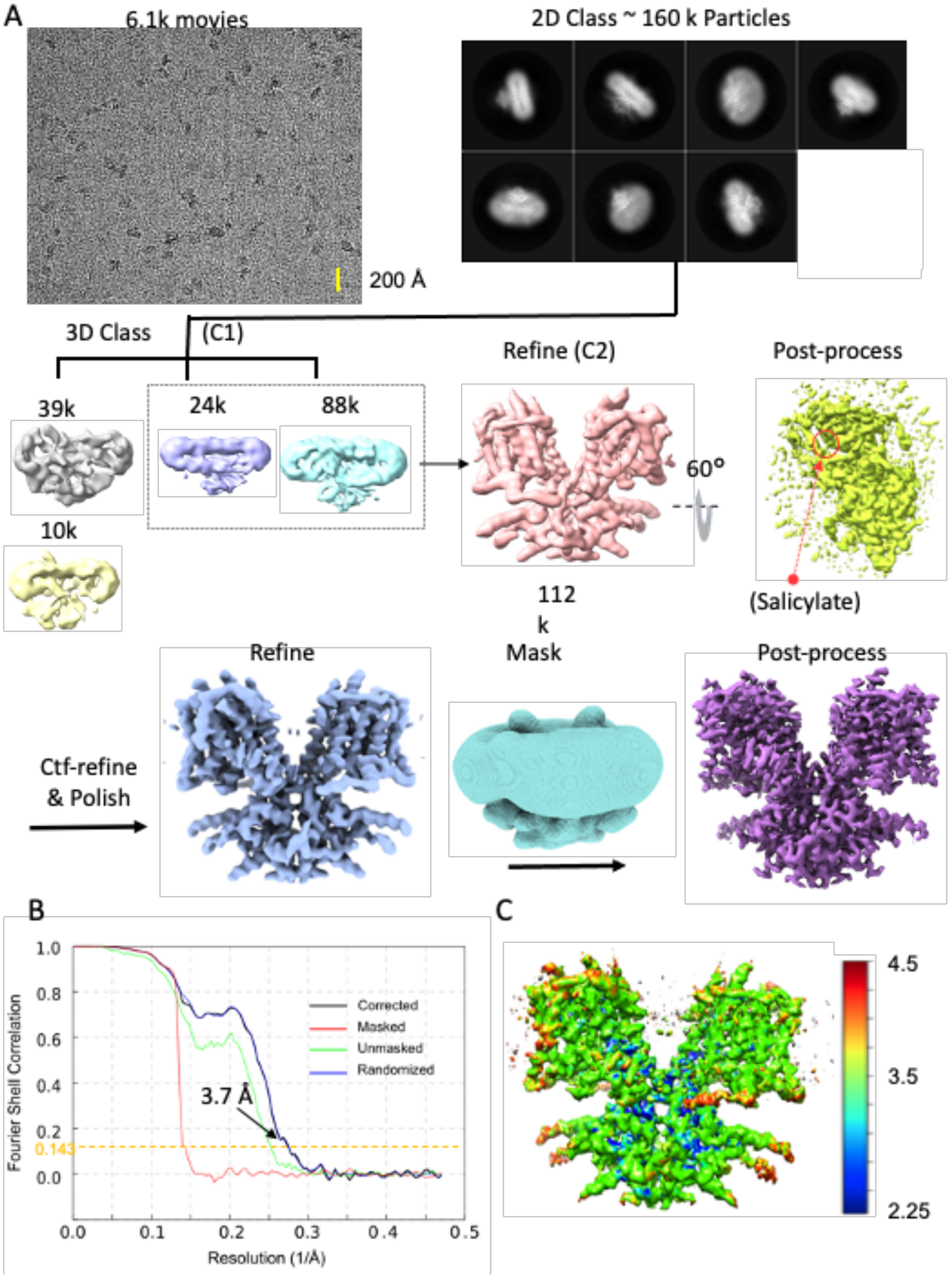


**Fig. S15.** Flow chart for the cryo-EM data processing and structure determination of the dolphin Prestin in the Inhibited II state ($SO_4^{2-}$ + Salicylate) (See Methods for details). The final reconstruction has a nominal resolution of 3.7 Å (at FSC=0.143). All the images in this figure were created in UCSF ChimeraX.

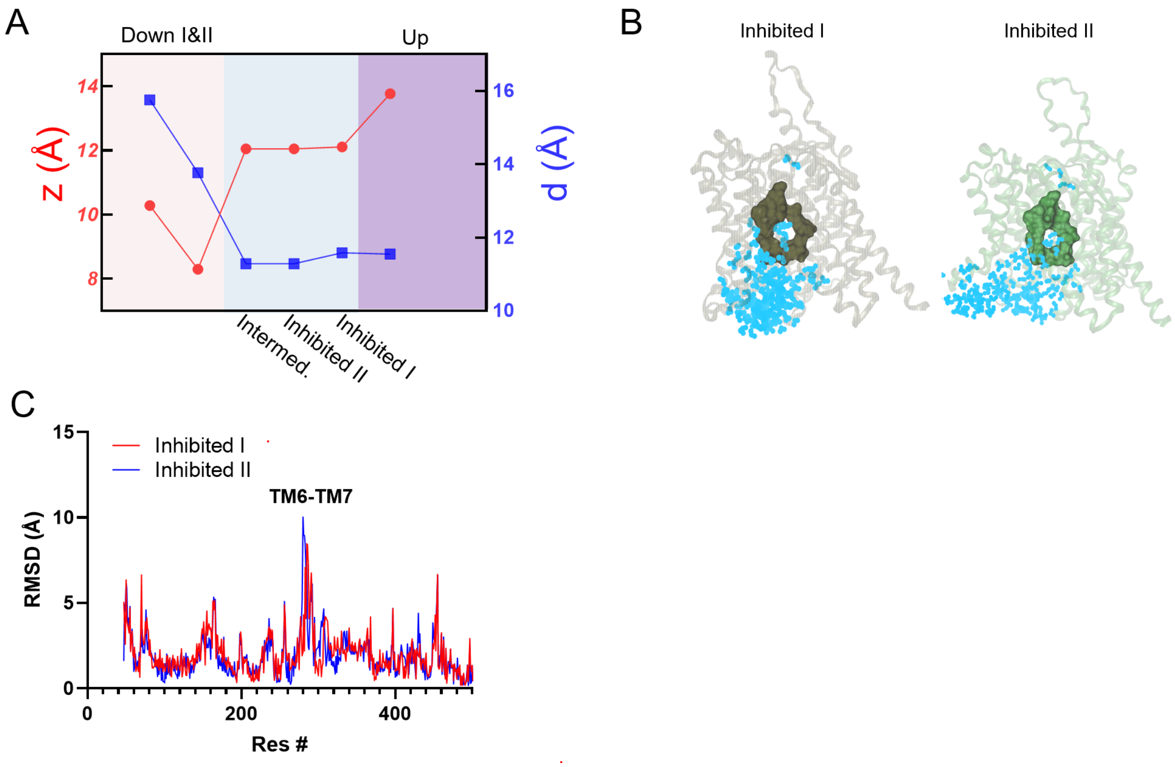


**Fig. S16.** Inhibited I & II (+ salicylate) states are closest to the Intermediate ($SO_4^{2-}$) state. (A) The vertical distance between R399 and V499 (as a measure for  the movement of the voltage sensor) , *z*, and the distance between F137 and R399, *d*, are calculated across states. When comparing the Inhibited states and the Intermediate state, differences between *z* and *d* are minimal, indicating a similar conformation of the voltage sensing and anion binding sites in the Inhibited and Intermediate states. (B) MD simulations shows that the anion-binding pocket (green) in inhibited I & II is not accessible by water (cyan).  The water molecules within 8 of the residues Q97, F101, F137 V397, S398, R399, E280 and E404 have been screened and illustrated using VMD. (C) RMSD calculation of Inhibited I & II (salicylate) states compared with the Intermediate ($SO_4^{2-}$) state. The major difference between them is in the TM6-TM7 helical dipole region as well as in the TM8 helix. The structures were aligned based on residue 460 to 505 (TM13-TM14).

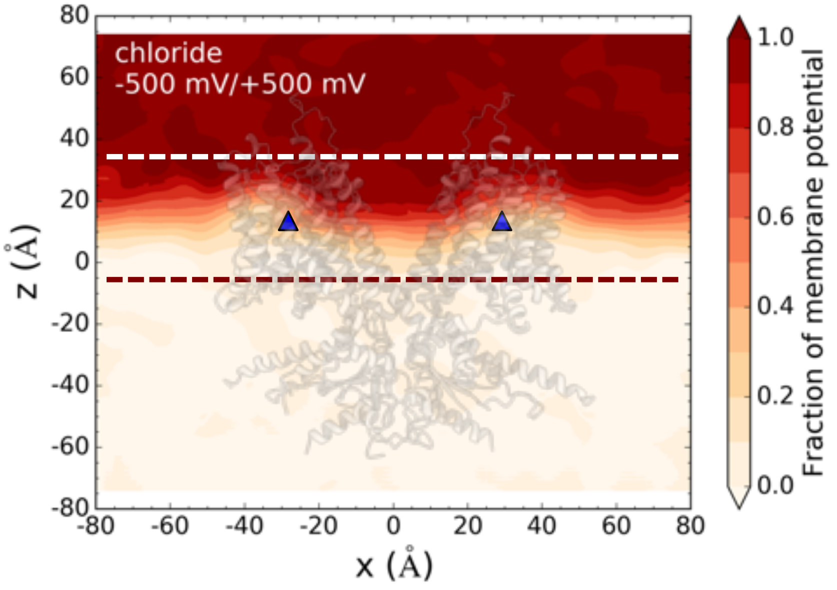


**Fig. S17.** The fraction of membrane potential around the binding sites of Up ($Cl^-$) state. The MD simulation box on the left and the corresponding 2-D fraction of membrane potential in the *x*-*z* plane crossing the central binding sites.

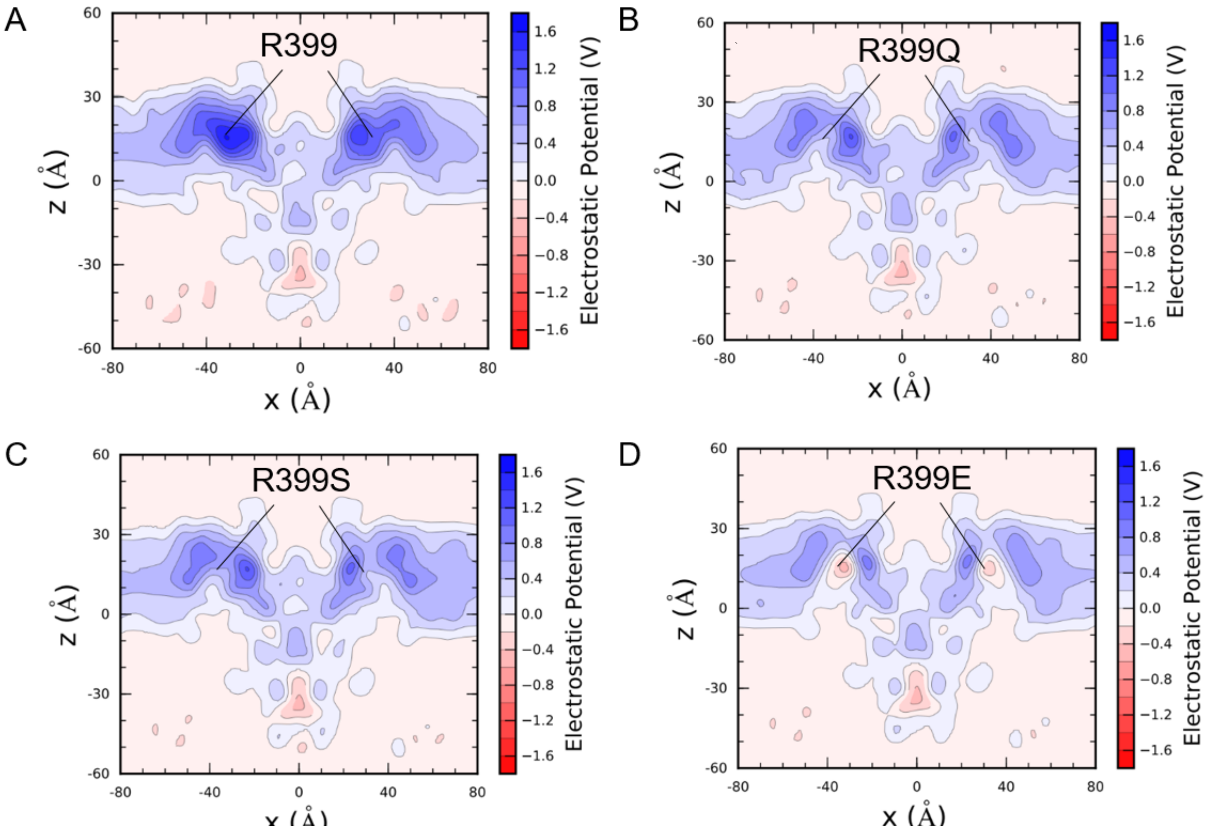


**Fig. S18.** 2D electrostatic calculations of Up state, without $Cl^-$ bound to either of the two binding sites, based on all-atom molecular dynamics simulation. (A-D) R399 in both monomers have been mutated to Q and S and E in different systems to see the contribution of R399 residue to the positive charge at the bilayer mid-plane. R399 mutation to polar residues shows that R399 has almost ~40% contribution the positive charge of the field at the bilayer mid-plane. The remainder likely comes from the TM3-TM10 helical dipole and other positive charges in this area.

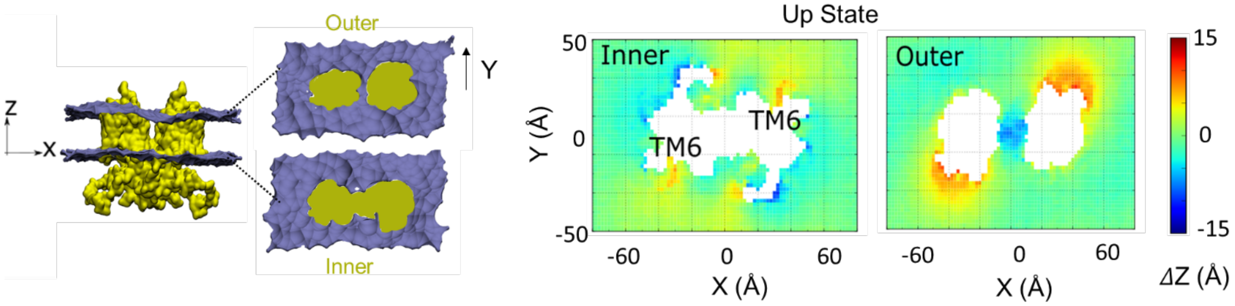


**Fig. 19.** MD simulation of Prestin (Up states) equilibrated in POPC lipid bilayers. The cross-sectional area of outer and inner monolayer with mapped leaflet coordinate in the Z direction (across the membrane thickness) using all-atom molecular dynamics simulations (1μs). The comparison was made between Up ($Cl^-$) and Inhibited II ($SO_4^{2-}$) states. The largest difference was observed at the location of the TM6 helix.

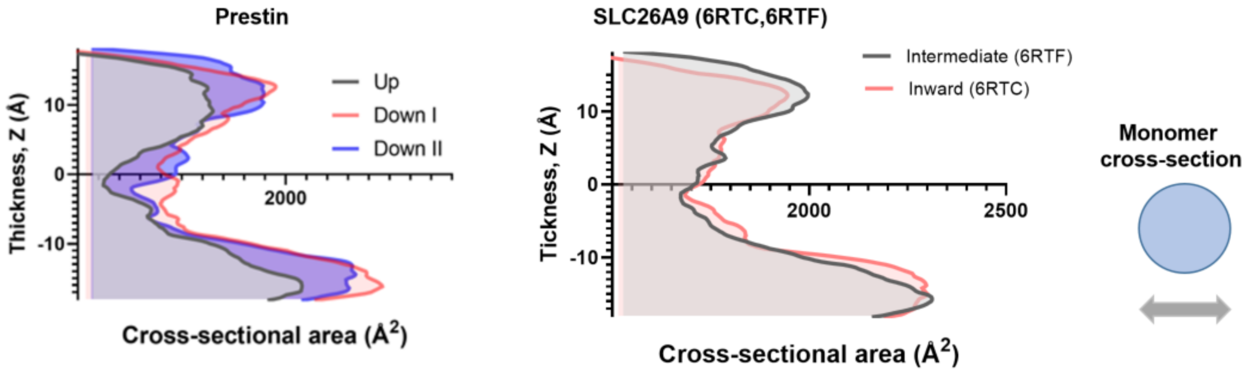


**Fig. S20.** Cross-sectional area calculations of the transmembrane domain of different dolphin Prestin structures compared to their corresponding structures of SLC26A9 along the hydrophobic thickness using CHARMM-membrane builder. Cross-sectional area change from Down to Occluded in Prestin and that of SLC26A9 from Inward-facing to Intermediate states (6RTC and 6RTF) per monomer. Note that prior to area calculation, the spatial arrangements of all the structures with respect to the hydrocarbon core of the lipid bilayer were first adjusted using the PPM server. The structures were aligned based on residues 460 to 550 (TM13-TM14).

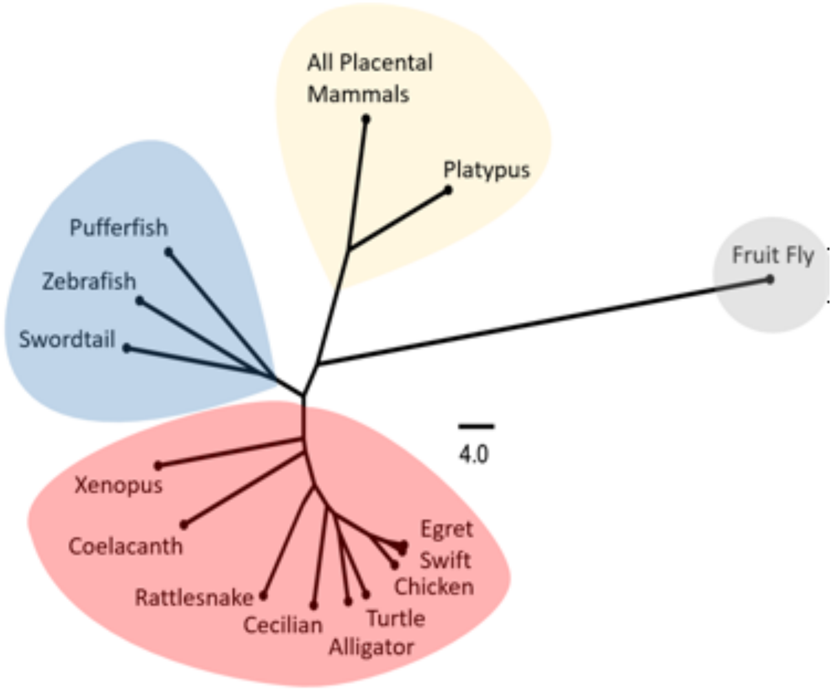


**Fig. S21.** Schematic phylogenetic tree of Prestin TM6 helix sequences, showing the nodes where ancestral Prestin genes have diverged.

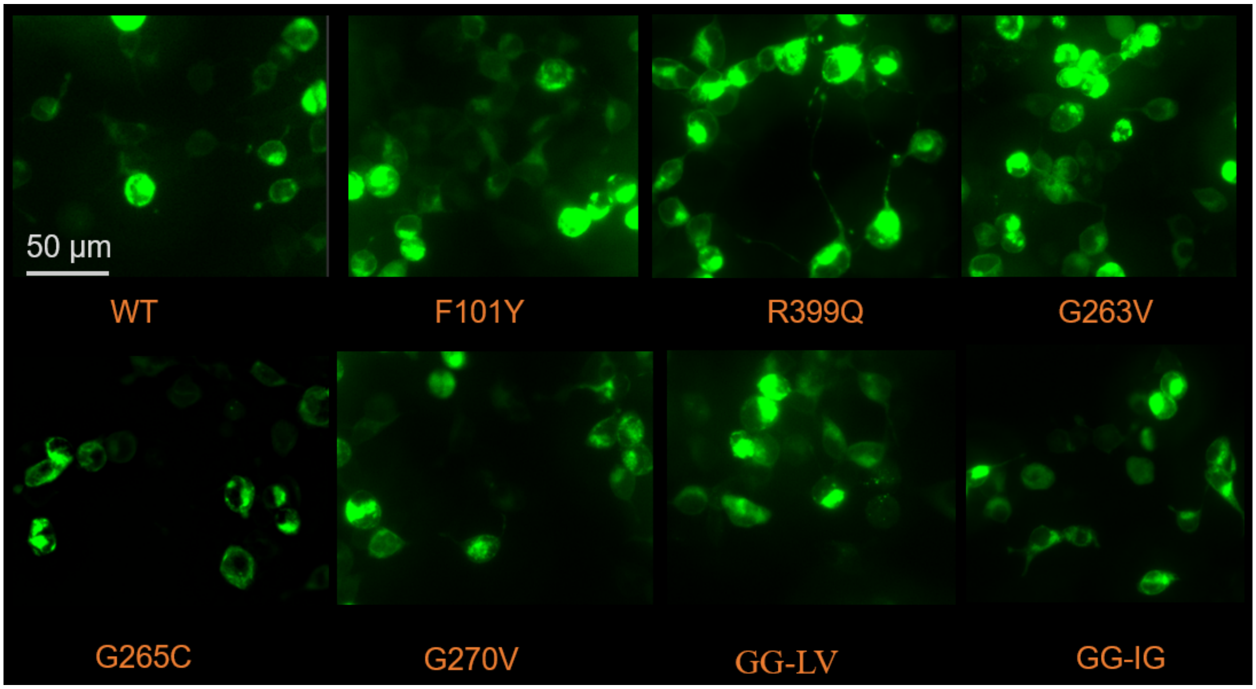


**Fig. S22.** Transient expression and membrane localization of wild type and mutant dolphin Prestin in HEK 293 cells. The grey solid line shows a scale bar of 50 µm. Prestin was over expressed in HEK 293 cells using the Prestin-C-EGFP pEG BacMam construct (see methods). Cellular localization of Prestin was detected by epifluorescence of prestin-EGFP (green) and images were captured by THUNDER microscopy (40X lens; Leica Microsystems). The cells were imaged between 32-36 hours post transfection (repeated 3 times). Both wild type and mutant Prestin are detected in the plasma membrane including in the phillopodia regions (which is most likely devoid of other cellular organelles). No visible difference was detected in terms of plasma membrane trafficking between the mutants and the wild type Prestin. Except that mutation G265C showed a relatively slower rate of expression (expresses post 30 h) compared to wild Prestin and other mutant (expresses post 24 h). Nevertheless, after this period, they are indistinguishable. Each transfection and imaging has been repeated at least three times.

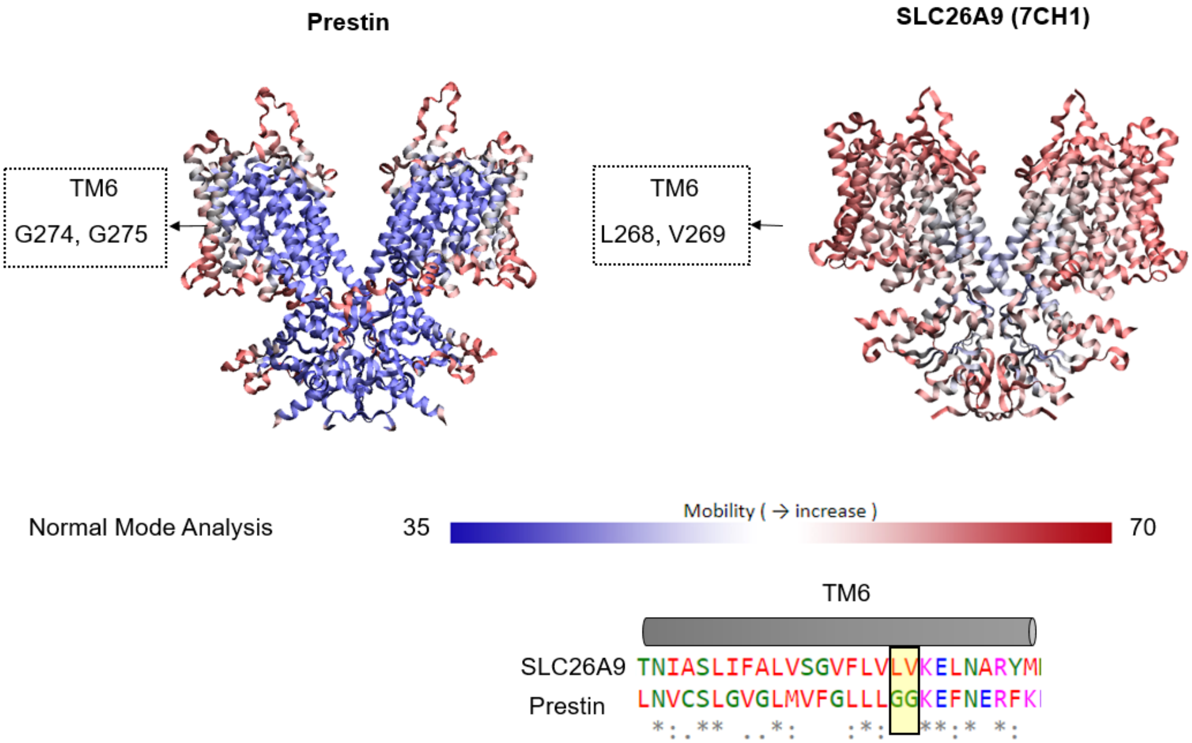


**Fig. S23.** Mobility and flexibility of Prestin versus SLC26A9**.** Degree of mobility of dolphin Prestin and human SLC26A9 (7CH1), mapped onto their corresponding structure using the DynOmics ENM server and normal mode analysis (*52*). This map shows that while TM6-TM7 region in Prestin has distinct higher mobility from the rest of Prestin structure, the corresponding helices in SLC26A9 have the same level of mobility as several other TM helices in the core and gate domain (indicative of a rigid body motion of the whole region).

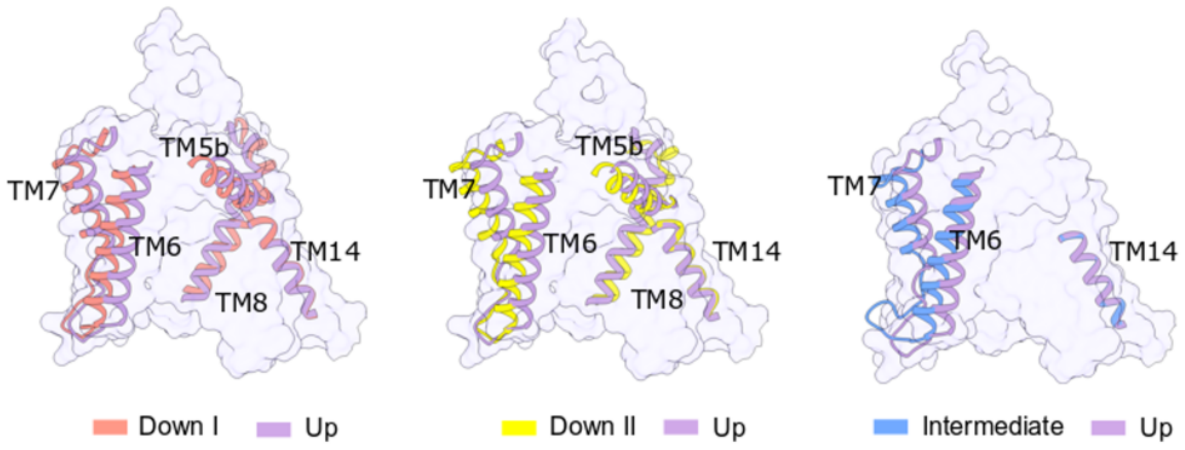


**Fig. S24** Upon the transition from Down I to Up state and the movement of the anion-binding site, the most obvious changes are seen in the Periphery helices TM5b, TM6-TM7, and TM8.

**Movie S1.**
The electromotility measurements of HEK 293 cells transfected with dolphin Prestin using whole-cell patch clamp electrophysiology. To evoke Prestin-mediated electromotility, The membrane potential was held at -70 mV; a 10-mV increase-in-amplitude voltage steps were applied up to the final steps which was from +150 mV to -140 mV (Fig. 1B). The pink square indicates the area that was chosen in our custom-written code to track the cellular displacements.

**Movie S2.**
Movement of the anion-binding site (i.e., the voltage sensor) from Down I to Down II conformation. For clarity, only one monomer has been shown. The anion-binding site has been highlighted in red and R399 has been shown in Stick representation and in yellow. The movies are made in UCSF ChimeraX.

**Movie S3.**
Structural changes from Down I to, Down II, Intermediate and Up conformation in series. For clarity, only one monomer has been shown. The anion-binding site has been highlighted in red and R399 has been shown in Stick representation and in yellow. The movies are made in UCSF ChimeraX.

**Movie S4.**
Structural changes from Down I to, Down II, Intermediate and Up conformation in series. The side front and top views of the dimer have been shown in one single frame. The anion-binding site has been highlighted in red and R399 has been shown in Stick representation and in yellow. The movies are made in UCSF ChimeraX.
